# The Encyclopedia of DNA Elements

**DOI:** 10.64898/2026.07.06.731365

**Authors:** The ENCODE Project Consortium, Timothy E. Reddy

**Affiliations:** Altius Institute for Biomedical Sciences

## Abstract

We present the Encyclopedia of DNA Elements (ENCODE), a reference map of the genomic basis of gene regulation. A product of more than two decades of systematic interrogation of genome function, ENCODE encompasses more than 16,000 genome-wide experiments, predominantly in primary cells and tissues, focused on three core layers of genome function. First, ENCODE now provides a catalog of gene regulatory elements. The catalog is based on a foundation of 5.3 million DNase I hypersensitive sites that delineate essentially all chromatin-accessible regulatory DNA in the human genome, as well as extensive maps of chromatin states, transcription factor occupancy, and nascent transcription, and systematic predictions of the functional consequences of non-coding genetic variants on regulatory element activity. Second, ENCODE expands the catalog of genes and transcripts, which now includes nearly 18,000 novel human long noncoding RNA genes, nearly 150,000 novel transcript isoforms, and genome-wide maps of transcript stability across cell types and time. Third, ENCODE now maps physical and functional interactions among regulatory elements and genes across more than 100 human tissues and cell lines at up to 10 bp resolution. Those studies reveal a vast network of interactions among millions of loop anchors across and links those interactions to gene expression. Through parallel studies in mice, ENCODE also provides extensive maps of gene regulatory elements, transcripts, and their interactions across the mouse postnatal development. Together, the Encyclopedia of DNA Elements provides a foundational framework for genome-focused studies of human and mouse biology.

## Introduction

The human genome is a sequence of three billion nucleotides that contain the heritable information required to create life.^1–3^ Encoded within the human genome is a vast collection of DNA elements that mediate genome function, most critically genes and the regulatory elements that control their expression (Supplementary Box 1). Following the publication of the first draft sequence of the human genome,^1,2^ the Encyclopedia of DNA Elements (ENCODE) Project was launched to identify and characterize those elements.

ENCODE pursued this goal through systematic studies of genome activity and function. The first phase (2003–2007) focused on 1% of the human genome.^4^ The second phase of ENCODE (2007–2012), leveraging the development of high-throughput short read-DNA sequencing, expanded to genome-wide studies in human^5^ and initiated parallel projects focused on mouse^6^ and other model organisms.^7,8^ The third phase (2012–2015) expanded studies of primary human and mouse cells and tissues^9^ while also broadening the experimental and computational approaches used.

Here, we report the results from the fourth and final phase of ENCODE. This phase greatly expanded studies to capture a more complete representation of human and mouse biology. Specifically, ENCODE 4 studied diverse primary cells and tissues representing all major human and mouse anatomical compartments; commonly used cell lines; human and mouse developmental time courses; and responses of human cells to environmental and genetic perturbations (Supplementary Box 2). Across those samples, we cataloged and characterized three primary aspects of gene regulation: (i) genes, (ii) gene regulatory elements, and (iii) the interactions among them (Fig. 1a).

**Figure 1.**
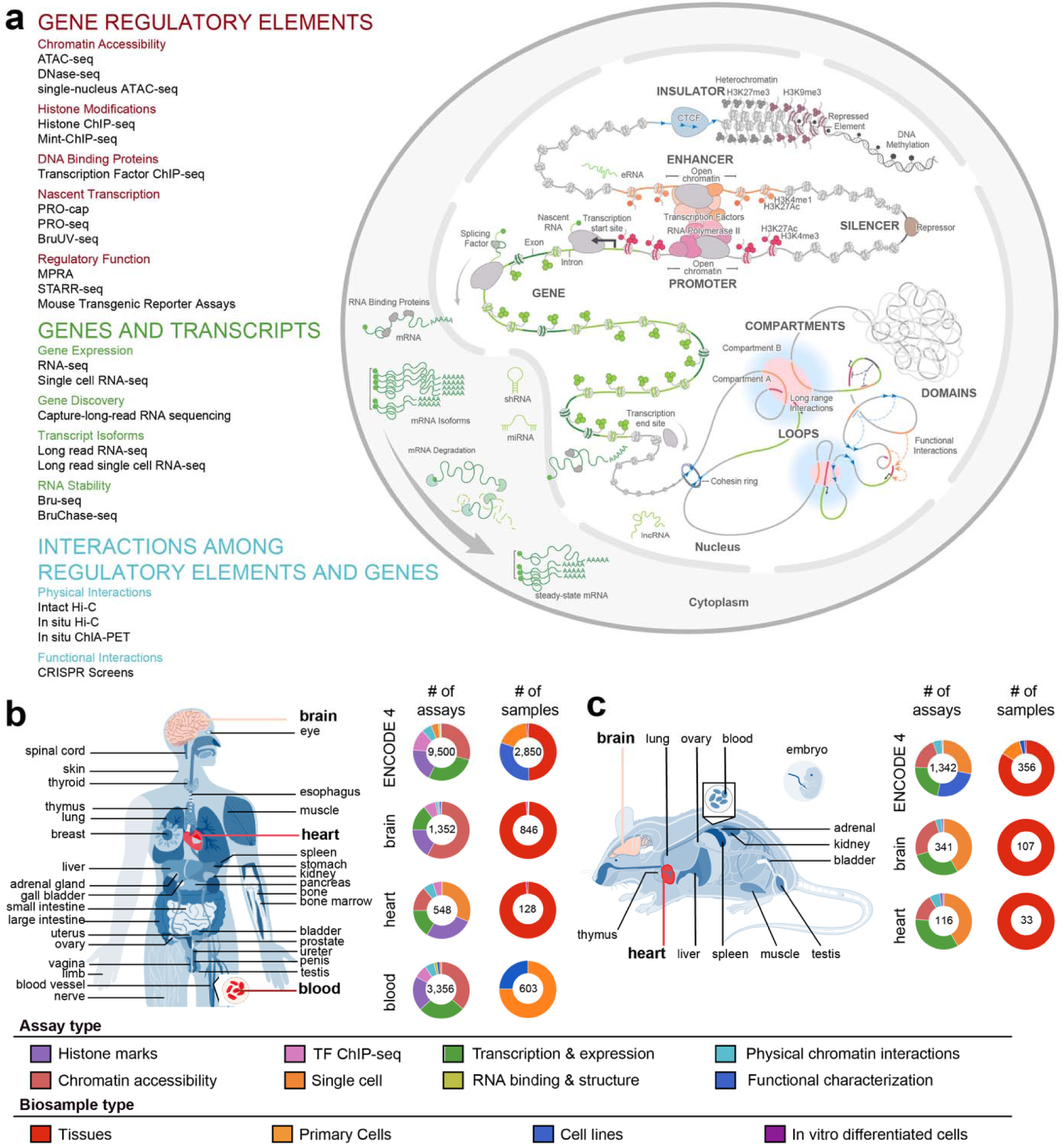
ENCODE 4 assays and samples. **a,** ENCODE 4 used a suite of complementary assays to measure three major components of gene regulation: (i) gene regulatory elements, (ii) genes and transcripts, and (iii) interactions among gene regulatory elements and genes. At right is a diagram of the different features of gene regulation those assays measured. **b and c,** ENCODE 4 assayed samples representing diverse human (**b**) and mouse (**c**) tissues. Pie charts indicate the distribution of assays and samples completed overall, and in specific deeply studied organs for each organism.

In total, ENCODE completed and released >16,000 genome-wide assays that probe various aspects of gene regulation. Of these, most (>9,500) are reported here for the first time (Supplementary Table 1). ENCODE assays studied >5,000 distinct biological samples including primary cells; primary tissues; in vitro differentiated pluripotent cells; and genetic or environmental perturbations thereof. We extensively coordinated data generation across labs from common samples, enabling numerous integrative and comparative assessments (Fig. 1b). In addition, the EN-TEx study^10^ enabled assessment of the functional contributions of non-coding variants by studying 30 tissues from four Genotype-Tissue Expression (GTEx) donors^11^ GTEx. We report here that ENCODE’s >16,000 assays enable annotation of tens of thousands of genes, hundreds of thousands of gene transcripts, millions of gene regulatory elements, and tens of millions of regulatory element interactions. We compiled these results into an Encyclopedia of DNA Elements to serve as a foundational reference for the study of human and mouse genomes function.

We present the Encyclopedia in five sections. First, we report comprehensive identification and systematic characterization of gene regulatory elements and the DNA sequences underlying them in the human genome. Second, we report expanded annotations of genes and transcripts, including their stability over time. Third, we annotate physical and functional interactions between genes and regulatory elements, and their impacts on gene expression. Fourth, we detail expansion of ENCODE annotations to the mouse genome. Finally, we describe key tools for accessing and using ENCODE data.

## Results

### Gene regulatory elements

From the outset, a major focus of ENCODE has been to identify and characterize gene regulatory elements encoded within the human genome sequence. Regulatory elements are characteristically accessible to nucleases and other enzymes, are bound by transcription factors, and may be flanked by histones with specific covalent modifications. Distal regulatory elements may also produce short unstable transcripts known as enhancer RNAs.

#### A comprehensive assessment of human chromatin accessibility

Genome-wide mapping of chromatin accessibility has been central to ENCODE’s effort to map the location of regulatory elements because it is a general marker of regulatory elements and can be deployed across a wide range of samples. During the fourth phase of ENCODE, we expanded maps of DNase I hypersensitive sites (DHSs) to encompass 4,159 distinct cell types, tissue types, and cell states (representing 3,125 conditions using reference ontologies) (<u>DNase-seq Datasets</u>). That is a ∼5.5-fold increase over previous ENCODE reports.^9^ The new samples included significantly expanded coverage of human fetal tissues^12^, adult brain, and immune cells. We also mapped chromatin accessibility over a subset of those samples using complementary bulk and single-cell ATAC-seq assays. The latter allowed us to infer chromatin accessibility for distinct cell types in their native environment inside heterogeneous tissues (Supplementary Table 2, Supplementary Figure 1).

The final ENCODE Encyclopedia catalog of human DHSs comprises 5.3 million high-confidence chromatin accessible sites within the uniquely mappable human genome sequence (http://regulome.org) (Fig. 2a).^13^

**Figure 2.**
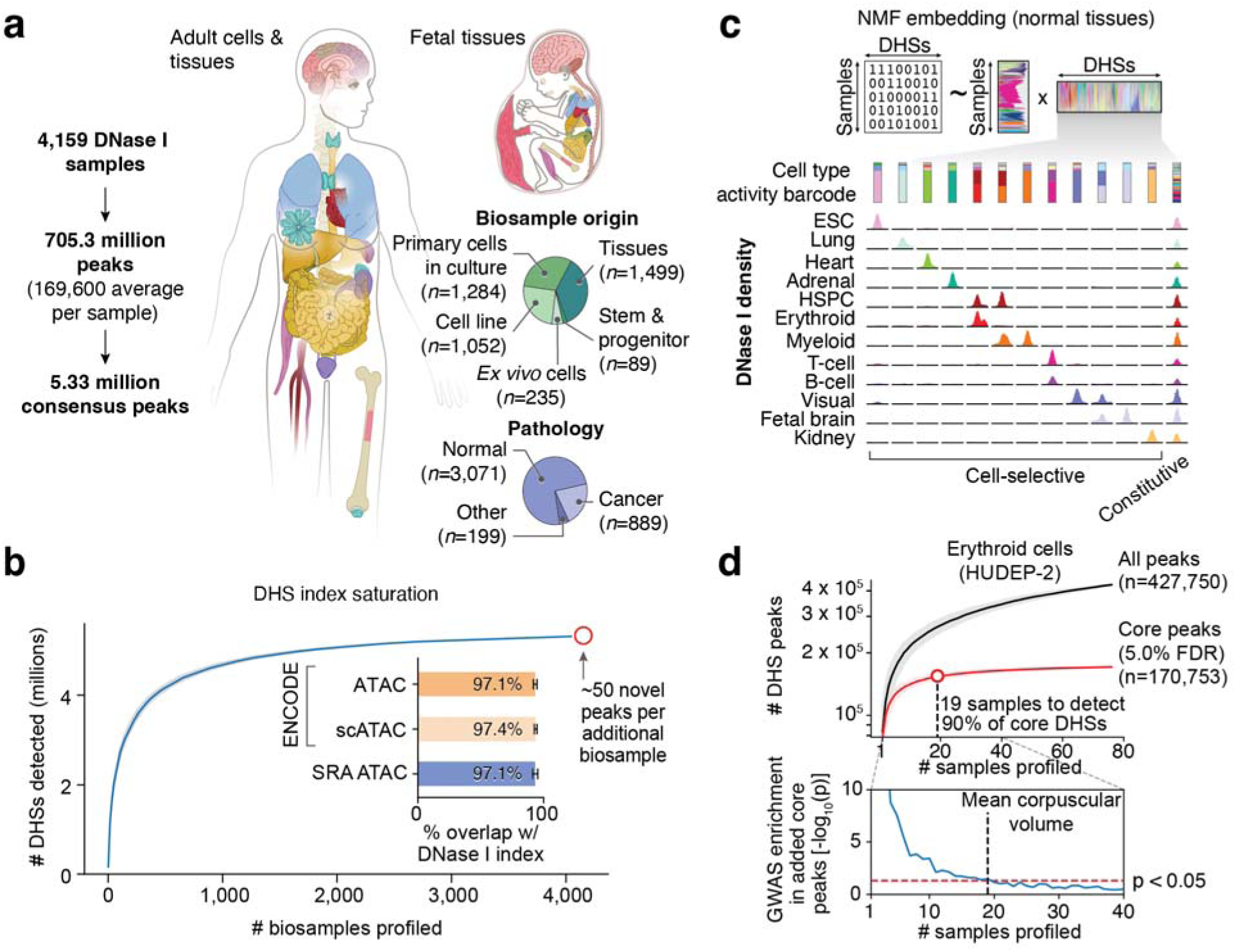
Chromatin accessibility across human cells and tissues. **a**, Construction of the reference index of chromatin accessibility. DNase I chromatin accessibility mapping was performed on 4,159 diverse samples yielding ∼705 million total peaks (∼170,000 per sample) and then combined to delineate an index of 5.33 million consensus regions. **d**, to visualize the cell selectivity of regulatory DNA, we created an activity barcode that represents the cell types and states in which each DHS is active. Specifically, non-negative matrix factorization was used to define cell-selective activity profiles for each index DHS (*n*=5.33 million) (top). Shown are the activity barcodes and normalized DNase I densities for 9 representative DHSs (bottom). **c**, A discovery curve showing the number of newly detected peaks as a function of the number of samples profiled. **d**, Saturation analysis of DNase I peaks for an individual cell type (erythroid HUDEP-2). Top: peak detection in HUDEP-2 cells as a function of number of replicates. Grey line indicates the mean of all peaks detected (adjusted p-value ≤ 0.01 in one or more samples) and the red line indicates the mean number of core peaks (adjusted *p*-value < 0.01). Shaded regions correspond to the 95% confidence interval. Bottom: Enrichment (log_10_ *p*-value) of mean corpuscular volume (MCV) trait associated genetic variation in newly added core DHS peaks after the profiling of additional HUDEP-2 samples.

Discovery of DHSs at new genomic positions dramatically diminished as new samples were profiled regardless of cell or tissue type, suggesting saturation of the DHS landscape. To assess further the completeness of the ENCODE DHS catalog, we compared it to elements identified from >20,000 uniformly processed publicly available non-ENCODE chromatin accessibility datasets. We found that nearly all elements from non-ENCODE datasets were represented within the ENCODE catalog despite many datasets originating from biological contexts not examined by ENCODE or from single cell assays. We estimate that a future chromatin accessibility experiment will annotate on average ∼50 new elements (Fig. 2b). As such, the 5.3 million elements in the ENCODE DHS catalog is, for most purposes, complete. This fulfills one of the key goals of ENCODE.^14^

#### Patterns of chromatin accessibility across cell types

Regulatory elements exhibit diverse activation patterns across cell types (http://regulome.org). Overall, we found that each reference DHS was accessible in a median of 25 cell contexts, while only ∼0.5% of elements were detected in ≥80% of samples. Most widely detected DHSs were promoters or sites bound by the chromatin regulator CTCF. Cells from related tissues share similar DHS landscapes, and most DHSs from normal cells and tissues could be assigned to one of ∼28 DHS landscape categories (Fig. 2c). In contrast, malignant cells frequently activated DHSs that were not typically accessible in their corresponding cell lineage of origin but rather were found in non-cognate cell lineages.

Many cell types were repeatedly sampled to capture donor- or condition-specific variation. Those deep analyses reveal that a core set of DHSs are recurrently activated within a given cell type context, whereas others are only detected sporadically (Fig. 2d). In any individual sample from a given cell type, only ∼50% of the core DHSs were observed. Distinguishing the core DHSs for a cell type typically required >20 distinct biological samples from that cell type. As such, studies based on only one or a few samples may considerably underestimate the extent of regulatory DNA within a given cell type.

#### Annotating regulatory elements using chromatin state

Covalent histone modifications are markers of genome function. We therefore assayed covalent histone modifications representing key aspects of gene regulation across hundreds of different tissues and cell types. Specifically, we measured the distribution of six histone modifications associated with enhancer or promoter activity (H3K27ac, H3K4me1, H3K4me3), gene bodies (H3K36me3), and facultative (H3K27me3) or constitutive (H3K9me3) heterochromatin across ∼450 different cell and tissue contexts using a combination of histone ChIP-seq and Mint-ChIP assays^15^ (<u>Mint-ChIP and Histone ChIP-seq Datasets</u>). Because of its major influence on chromatin structure, we also assayed CTCF occupancy in many of the same cell types using ChIP-seq (<u>CTCF</u> <u>ChIP-seq Datasets</u>). Using those datasets, we systematically annotated regulatory activity states across the human genome using the ChromHMM^16^ and Segway^17,18^ semi-supervised hidden Markov models (<u>ChromHMM and Segway Analyses</u>). We also used local histone modification patterns to assign many DHSs into functional categories of candidate cis-regulatory elements (cCREs, see below).

We also used histone modification data to identify repressive regulatory elements and structures. Silencers were systematically characterized using several approaches (Supplementary Figure 2).^19–24^ In ENCODE 4, we found that H3K27me3-rich genomic regions physically interact over long distances to form higher-order regulatory units with strong transcriptional repression, which we termed super-silencers.^25^

#### Genome binding profiles for most human transcription factors

Previous phases of ENCODE measured the occupancy of ∼500 transcription factors and chromatin-associated proteins via ChIP-seq.^26,27^ The final phase expanded those measurements to 1,100 proteins, representing ∼70% of human transcription factors (<u>Transcription Factor ChIP-seq Datasets</u>).^28^ For many newly studied proteins, including hundreds of zinc-finger transcription factors, antibodies with sufficient specificity for ChIP-seq do not exist. To circumvent that challenge, we introduced epitope tags into the endogenous coding sequences of 729 such transcription factors and performed ChIP-seq using antibodies against those epitope tags (Supplementary Table 3).^29^ Because epitope tagging and selection at the scale required for ChIP-seq is most feasible in immortalized cell lines, we concentrated those efforts on HepG2, a liver carcinoma cell line, and K562, a leukemia cell line.

The TF ChIP-seq datasets collectively identify ∼80,000 and 40,000 genomic regions TF-bound in HepG2 and K562 cells, respectively. We detected significant ChIP-seq signal over ∼10% of the human genome (Supplementary Figure 3) though, as detailed below, only a fraction of that signal is explained by sequence-specific DNA:protein interactions. ChIP-seq signal for different transcription factors commonly clusters within ∼100 bp genomic regions, and we found thousands of high-occupancy target sites with evidence of many transcription factors binding together.^30^ At many other genomic sites, we identified modules of transcription factors, such as the activator protein 1 (AP-1) transcription factors, that bind at the same sets of genomic locations.^31^ Some modules are shared across cell lines, whereas others were cell-type specific. However, cell-shared modules often bind different locations in different cell types. Similarly, the same genomic location often binds different modules of transcription factors in different cell lines. As discussed below, those differentially bound regions frequently coincide with chromatin loops, where co-binding patterns dictate whether the loop serves a primarily structural or regulatory purpose in each cell type.^32^

#### Identifying regulatory elements using nascent RNA

Promoters are defined by their ability to initiate RNA Polymerase II transcripts (protein-coding or non-coding), while enhancers may originate short transcripts that mark an active regulatory state.^33^ ENCODE mapped nascent transcripts using nuclear run-on sequencing (PRO-seq and PRO-cap)^34^ and BruUV-seq.^35^ Overall, we identified 715,296 transcriptionally active regulatory elements across 215 human tissue and cell samples using PRO-cap (<u>PRO-cap Datasets</u>).^36^ We also measured the effect of protein degradation and glucocorticoid treatments on transcription using PRO-cap and PRO-seq (<u>Targeted Degradation Datasets</u>, <u>Glucocorticoid Response Datasets</u>). The BruUV-seq assays measured transcription initiation in 16 human cell lines as part of a larger coordinated study of gene regulation in those lines (<u>Coordinated Cell Line</u> <u>Datasets</u>).^37,38^

Transcriptional initiation from regulatory elements corresponds to other indicators of regulatory activity. For example, in K562 cells, ∼80% of transcriptionally active regions tested had activity in high-throughput reporter assays.^36,39^ Moreover, such regions often exhibit both promoter and enhancer activity, and these activities are strongly correlated.^40^ Computational models of the sequence determinants of transcriptional initiation showed that promoter and enhancer activity share common sequence determinants^40^ and can also inform genetic contributions to health and disease.^36,38,41,42^ Further insights into regulatory element activity can be gleaned by evaluating the location and direction of transcription initiation sites within regulatory elements. As examples, transcription factors such as CTCF and ZNF143 may contribute to the direction of transcription from regulatory elements, and unidirectionally transcribed elements are evolutionarily younger and have weaker sequence constraint than divergently transcribed elements.^36,43^

#### Systematic categorization of candidate cis-regulatory elements using ENCODE data

Each of the above datasets gives complementary information about the location and activity of regulatory elements. In the previous phase of ENCODE, we developed a rule-based approach to categorize candidate cis-regulatory elements (cCREs) by assigning DNA segments containing DHSs into one of seven regulatory states based on proximity to transcription start sites, histone modification patterns, and transcription factor occupancy.^9^ In this final phase, we expanded that categorization to include ∼2.4 million elements (∼45% of annotated DHSs) from 1,679 samples based on updated criteria including nascent transcription (Supplementary Figure 4a) (<u>cCRE Analyses</u>).^24^ Promoter-like and enhancer-like cCREs had higher reporter assay activity than all other classes of cCREs.^44^ In addition, cCREs were most strongly enriched for activity in the cell type where they were identified, particularly for CRISPR perturbation assays (Supplementary Figure 4b).

#### Functional assessment of regulatory element activity

In its final phase, ENCODE initiated a systematic effort to probe the regulatory potential of DNA sequences using functional assays. We performed ∼600 large-scale functional characterization experiments, focusing on functional reporter assays and CRISPR perturbations spanning 35 human and mouse cell types and tissue (<u>Functional</u> <u>Characterization Datasets</u>). Reporter assays included genome-wide measurements of regulatory activity, coordinated dense studies of specific ∼1 Mb loci (<u>High-throughput</u> <u>Reporter Assay Datasets</u>), and *in vivo* whole-organism reporter assays of ∼300 individual regulatory elements in day 11.5 transgenic mouse embryos (<u>Transgenic</u> <u>Mouse Reporter Datasets</u>). The CRISPR perturbation studies included a coordinated multi-center analysis of 13 Mb of the human genome in five cell types^45^ as well as targeted studies of disease-associated regions^46^ and of the promoters of dosage-sensitive genes.^47^

Collectively, we report evidence of functional regulatory activity for tens of thousands of genomic elements,^45^ including >33,000 with activity in multiple assays in K562 cells.^44^ We found reporter assays to be more permissive for regulatory activity than CRISPR screens, exposing regulatory activity for DNA sequences that are found both in accessible chromatin in that cell type and in non-accessible DNA that coincided with DHSs in other cell types. In contrast, significant results from CRISPR interference screens were almost entirely limited to genomic regions with accessible chromatin, transcription factor binding, and reporter assay activity in the cell context tested. Those results are consistent with reporter assays measuring the potential of DNA sequences to regulate a nearby promoter, whereas CRISPR screens reflect additional constraints such as chromatin accessibility and the ability to physically reach the target promoter.^44,48,49^

#### Sequence determinants of regulatory element activity identified by predictive models

Regulatory DNA function is mediated by the cooperative binding of sequence-specific transcription factors. However, identifying the relative contribution of factor recognition sites to regulatory DNA activity has proven challenging. To do so, we trained convolutional neural networks that use local DNA sequences to predict base-resolution signal profiles for several assays of regulatory element activity including chromatin accessibility,^50^ transcription factor binding,^51–53^ nascent transcription, and activity in reporter assays (Fig. 3a).^54,55^ We trained separate models for each assay type in each tissue type using five-fold cross-validation. In total, we created models for 2,339 TF ChIP-seq experiments (including 1,261 datasets for 455 unique C2H2 zinc fingers), 1,143 DNase-seq experiments, 369 ATAC-seq experiments, 6 PRO-cap experiments, and 8 high-throughput reporter assay experiments in both human and mouse (Supplementary Table 4) (<u>ChromBPNet and BPNet Models</u>). Experiment-specific models accurately predict the position, strength, and shape of signal with base-pair resolution and in an assay- and cell-type-specific manner (Supplementary Figure 5).

**Figure 3.**
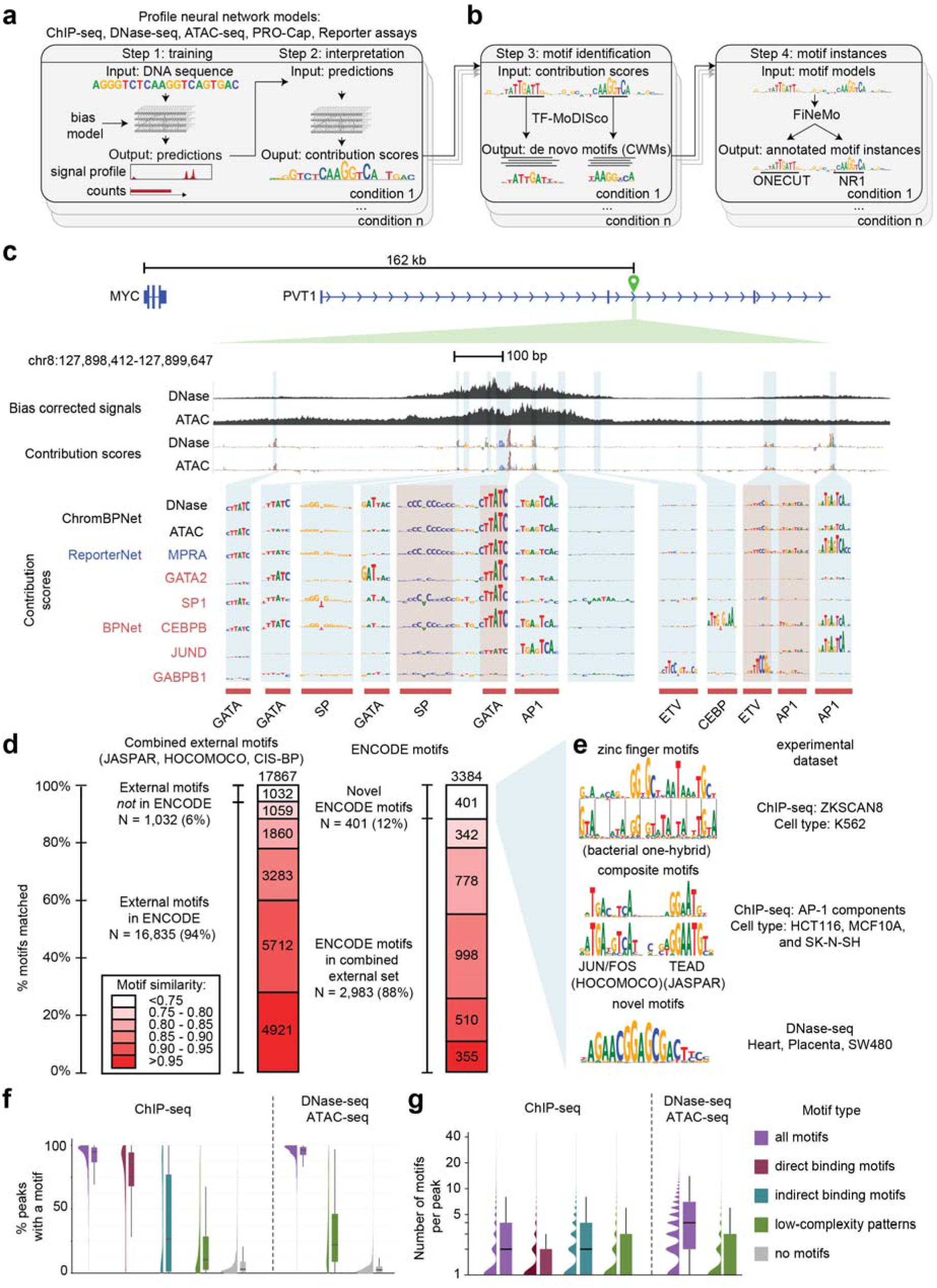
Predictive DNA sequence models and unified motif interpretation. **a,** A schematic of the sequence modeling and downstream analysis methods. Convolutional neural network models (BPNet: TF ChIP-seq, ChromBPNet: DNase/ATAC-seq, ReporterNet: Reporter assays, ProCapNet: PRO-cap) are trained per experiment to predict both total signal (count) and base-resolution profile shape from local DNA sequence, while explicitly modeling and correcting assay specific biases. **b,** Interpretation tools are used on trained models to get contributions of individual bases in the input sequences, summary of predictive motifs and the locations of motifs within the input sequences. DeepLIFT computes base-resolution contribution scores; TF-MoDISco clusters high-contribution subsequences into *de novo* contribution weight matrix (CWM) motifs; FiNeMo competitively scans contribution maps with CWM libraries to call motif instances; MotifCompendium clusters CWMs across models into a global motif index for cross-context comparison. **c,** Model-derived browser tracks at a CRISPRi-validated distal enhancer at the MYC locus in K562 (chr8:127,898,412–127,899,647; ∼162 kb downstream of the MYC promoter). From top to bottom: bias-corrected, base-resolution predicted DNase-seq and ATAC-seq profiles (ChromBPNet) with corresponding contribution maps. Insets compare contribution maps across DNase/ATAC (ChromBPNet), MPRA (ReporterNet), and TF ChIP-seq models (BPNet; e.g., GATA2, SP1, CEBPB, JUND, GABPB1), with high-impact motif instances annotated (e.g., GATA, SP, AP-1, ETV/ETS, CEBP). **d,** Overlap between our consolidated model-derived motif catalog and combined external motif databases (JASPAR 2026-Core Homo sapiens, HOCOMOCO V14-CORE Human, and CIS-BP V3-Homo sapiens) across similarity thresholds. Left bar plot shows that most external motifs map to our catalog (16,835/17,867; 94%), while a significant fraction of our motif catalog is not captured by the external databases (401/3,384; 12%). **e,** Examples of motifs in our catalog without strong matches in external databases, including candidate C2H2 zinc-finger motifs supported by a bacterial one-hybrid (B1H) recognition code (e.g. ZKSCAN8 ChIP-seq model in K562), composite motifs consistent with TF–TF co-binding (e.g., JUN/FOS–TEAD composite from AP-1 ChIP-seq models in HCT116, MCF10A and SK-N-SH), and novel motifs discovered from accessibility models (e.g., DNase-seq model in heart, placenta, and SW480 cells). **f,** Distribution (violin and box plots) of percentage of peaks containing at least one model-derived predictive motif instance, stratified by motif class (all: all types of motifs, direct: known cognate motif of a TF, indirect: non-cognate motif of a different TF, low-complexity motifs, or no motifs) across all ChIP-seq (left) and DNase/ATAC-seq (right) datasets. **g,** Distribution (violin and box plots) of the number of model-derived predictive motif instances per peak across assays (ChIP-seq, DNase/ATAC-seq) and motif classes (same as in **f**).

We next created a unified interpretation framework to identify predictive sequence features influencing these signals (Fig. 3b).^51–54,56–58^ We created three datasets per experiment: (i) predicted signal profiles corrected for experimental noise and biases such as sequence preferences for enzymes used in each assay, (ii) nucleotide-resolution contributions to those signals, and (iii) the effects of those contributions on DNA motifs derived from all models (Fig. 3c). Each dataset for each experiment is provided as a tracks that can be viewed on a genome browser. We illustrate how they can be used to dissect the DNA contributions to regulatory activity in Case Study 1.

#### The ENCODE DNA motif catalog

We next created a unified ENCODE catalog of DNA motifs that predict different aspects of regulatory element activity across cell contexts. To do so, we consolidated >280,000 motifs models into a non-redundant catalog of 3,384 motifs, explaining transcription factor binding and chromatin accessibility.^52^ The ENCODE catalog encompasses most motifs found in other compendia (Fig. 3d), and also adds many novel motifs, especially for zinc fingers, for transcription factors regulating chromatin state, and for cooperative binding between transcription factors (Fig. 3e, Supplementary Figure 6).^52,53,58^

The ENCODE motif catalog also gives new insights into the sequences critical for gene regulation. Nearly all (∼95%) of ChIP-seq or chromatin accessibility peaks contained at least one motif instance that was predicted to influence signal strength (Fig. 3f). Overall, ChIP-seq signal can be explained by fewer motifs than chromatin accessibility signal (median of ∼2 vs ∼4 per peak, Fig. 3g), and by motifs that are closer to the signal summit and explain a greater fraction of the signal at each site (Supplementary Figure 7a). Most peak-associated signals were best explained by aggregate motif instances vs. individual motifs (Supplementary Figure 7b-c). Low-complexity features also accounted for a substantial fraction of ChIP-seq and chromatin accessibility signal (Supplementary Figure 7c). Individual motif instances also provide testable hypotheses about the context-specific syntax of gene regulatory DNA sequences. For example, we identified and validated examples of non-additive effects on regulatory element activity arising from cooperative interactions between multiple motifs, and between motifs and flanking DNA sequence context (Case Study 2).

#### Estimating variant effects on regulatory element activity

Non-coding genetic variants are major contributors to human traits.^59–61^ ENCODE assayed samples from hundreds of donors. The use of high-throughput sequencing for readout in assays such as DNase-seq allows us to identify genetic variants in regulatory elements in those donors’ genomes. At heterozygous positions, we can also quantify assay signal that is specific to each allele. Complementing that approach, we also used synthetic reporter assays to test the effects of large numbers of selected non-coding variants.

Across both approaches, ENCODE experimentally tested the effects of ∼1.5 million common and low frequency variants. We observed significant (FDR < 0.05) allelic effects at 149,679 (10%). The expanded allele-specific chromatin accessibility assessments in ENCODE 4 informed 80% more GWAS catalog variants and accounted for 27% of all novel trait-to-top-sample associations discovered (Supplementary Figure 9c).^62^ To expand beyond those experimental measurements, we also computationally predicted 10 million variant effects using eQTL and deep learning models (Fig. 4a, Supplementary Figure 8, Supplementary Figure 9a). The computationally predicted effects were enriched 13-fold and 10-fold for variants with allele-specific reporter assay activity and DNase hypersensitivity, respectively (Fig. 4b) (Supplementary Tables 5 and 6).^63^ In total, ENCODE 4 assays and models provide evidence for allelic-specific activity at 5% and 16% of GWAS catalog variants, respectively (Supplementary Figure 9b) (Supplementary Table 7).

**Figure 4.**
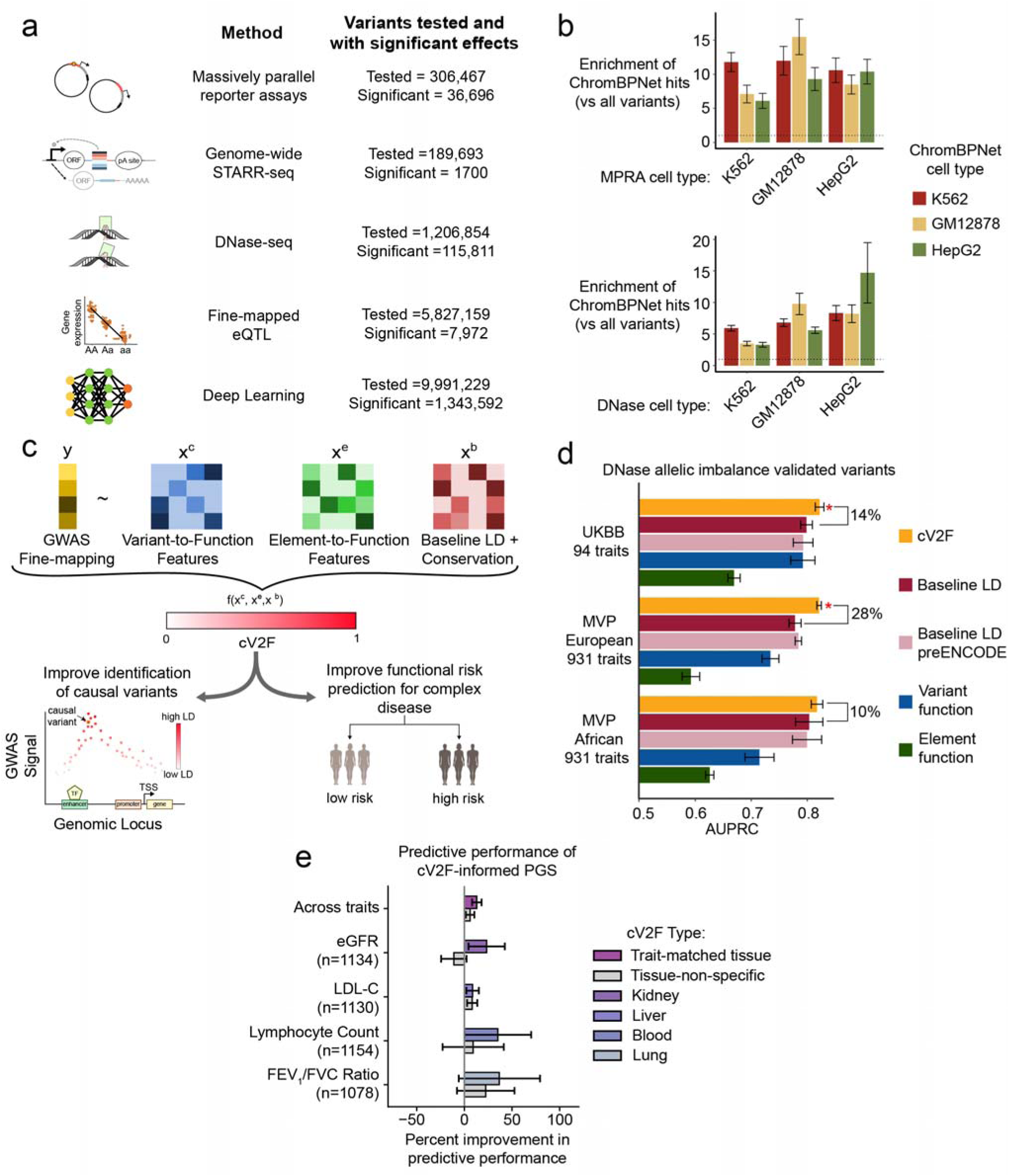
Effects of non-coding genetic variants on regulatory elements. **a,** Outline of the five different types of ENCODE Phase 4 variant function assays and models: (i) targeted massively parallel reporter assays, (ii) whole-genome STARR-seq, (iii) DNase I cleavage maps, (iv) AFGR eQTLs, and (v) ChromBPNet deep learning model. We report the proportion of common and low-frequency variants (1000 Genomes, minor allele count >=5) evaluated by each of these approaches and the proportion of the tested variants that were found to be significant. We report in parenthesis the number of samples corresponding to each of these assays or models. **b,** Cross-cell-type concordance of ChromBPNet predictions with experimental assays. Fold-enrichment of ChromBPNet hits (vs. all tested variants) among MPRA-significant variants (top) and DNase allelic imbalance variants (bottom) in K562, GM12878, and HepG2. Bar color denotes the ChromBPNet model cell type. Error bars denote 95% confidence intervals. The dotted line indicates no enrichment. **c**, Schematic of the cV2F model. Fine-mapped GWAS posterior inclusion probabilities (*y*) are modeled as a function of variant-to-function features, element-to-function features, and baseline-LD and conservation features (*x*). cV2F score can be used to prioritize variants for function in a cell-type agnostic and cell-type specific manner. **d,** Area under the precision and recall curve (AUPRC) of cV2F features, combining all variant-level functional assays and sequence-based models, evaluated against various sub-models trained only on variant-function or element-function features. Performance is evaluated using fine-mapped variants from UK Biobank (UKBB; 94 traits), and Million Veteran Program European-ancestry and African-ancestry (MVP European and MVP African; 931 traits) GWAS. Error bars denote 95% confidence intervals. Red asterisks denote significant improvement of cV2F over the next-best method (Leave-one-chromosome-out Jack-knife *P* < 0.05). **e,** Across four traits, tissue-matched polygenic risk scores doubled the average improvement over non-tissue-matched scores (6% to 13%, 95% CI: [3.3%, 23%], p=0.04). Numerical results are reported in Supplementary Tables 5-10.

Non-coding variants can contribute to phenotypes by impacting different aspects of gene regulation. We therefore created consensus variant to function (cV2F) scores that provide a single variant effect score for many traits (Supplementary Table 8) by learning optimal combinations of ENCODE datasets, sequence conservation and linkage disequilibrium (Fig. 4c) (<u>cV2F Scores</u>).^64^ Overall, ENCODE cV2F scores had a high precision-recall when predicting held-out UK Biobank trait associations (AUPRC = 0.82). The addition of ENCODE 4 data improved precision-recall by 14% on held-out UK Biobank fine-mapped variants and by 28% and 14% on Million Veterans Project fine-mapped associations in European-Americans and in African-Americans, respectively (Fig. 4d, Supplementary Table 9).^65–67^ Variant effect estimates and tissue-specific features were especially important for cV2F performance, and using tissue-matched cV2F in multi-ancestry polygenic prediction showed 13% improvement over the non-functional baseline in African ancestry individuals (Fig. 4e, Supplementary Figure 9f-g, Supplementary Table 10).^64,68,69^ Compared to existing approaches, cV2F scores had better concordance on held-out GWAS and reporter assay data (Supplementary Figure 9d-e); and increased the number of confidently fine-mapped UK Biobank trait variants by 14% (PIP > 0.95, N = 3,551 versus 3,107) (Supplementary Figure 9h). We detail example applications using cV2F to fine-map common and rare disease mechanisms in Case Study 3.

### Genes and transcripts

A second overarching goal of ENCODE 4 was to create and expand atlases of genes, gene expression, transcript isoform diversity, and transcript stability.

#### An expanded gene catalog

Cataloging all human and mouse genes, transcripts, and translations has been one of the major goals of ENCODE and of the offshoot GENCODE consortium. This phase of ENCODE improved and expanded the GENCODE gene catalog, primarily through the systematic application of capture-long-read RNA sequencing (CLS) across a wide-range of samples.^70,71^ Doing so substantially increased the number of annotated long non-coding RNAs (lncRNA) genes, adding 17,931 novel human genes (140,268 transcripts) and 22,784 novel mouse genes (136,169 transcripts). These novel lncRNA genes exhibit the features expected of bona fide transcriptional units even though they are poorly detected by bulk RNA-seq because their expression is highly restricted to specific cell populations. Many previously orphan promoters and enhancers map to those novel transcriptional units, as do tens of thousands of phenotype-associated variants. This new catalog increases the number of human disease-associated lncRNAs with mouse orthologs by 3-fold. Additionally, through mapping novel lncRNA transcripts in primate genomes, we predict hundreds of thousands of novel transcripts in other species, greatly enhancing lncRNA annotation in non-human primates.

#### Expanded maps of transcript isoforms via long-read RNA-seq

ENCODE 4 also expanded maps of isoform expression by profiling >400 full-length transcriptomes across a range of tissues and developmental stages using long0-read RNA-seq (<u>Long Read RNA-seq Datasets</u>).^72^ We detected at least one full-length transcript from 87% of annotated human protein coding genes in GENCODE v40 (Fig. 5a), with 89% expressing multiple transcripts. In total, we mapped >200,000 polyA+ transcripts, 35% of which have novel splicing compared to GENCODE v40 (Fig. 5b). In contrast, only 25% of lncRNAs and pseudogenes express multiple transcripts (Fig. 5c). Long-read RNA-seq datasets were also rich in allele-specific alternative splicing events.^73^

**Figure 5.**
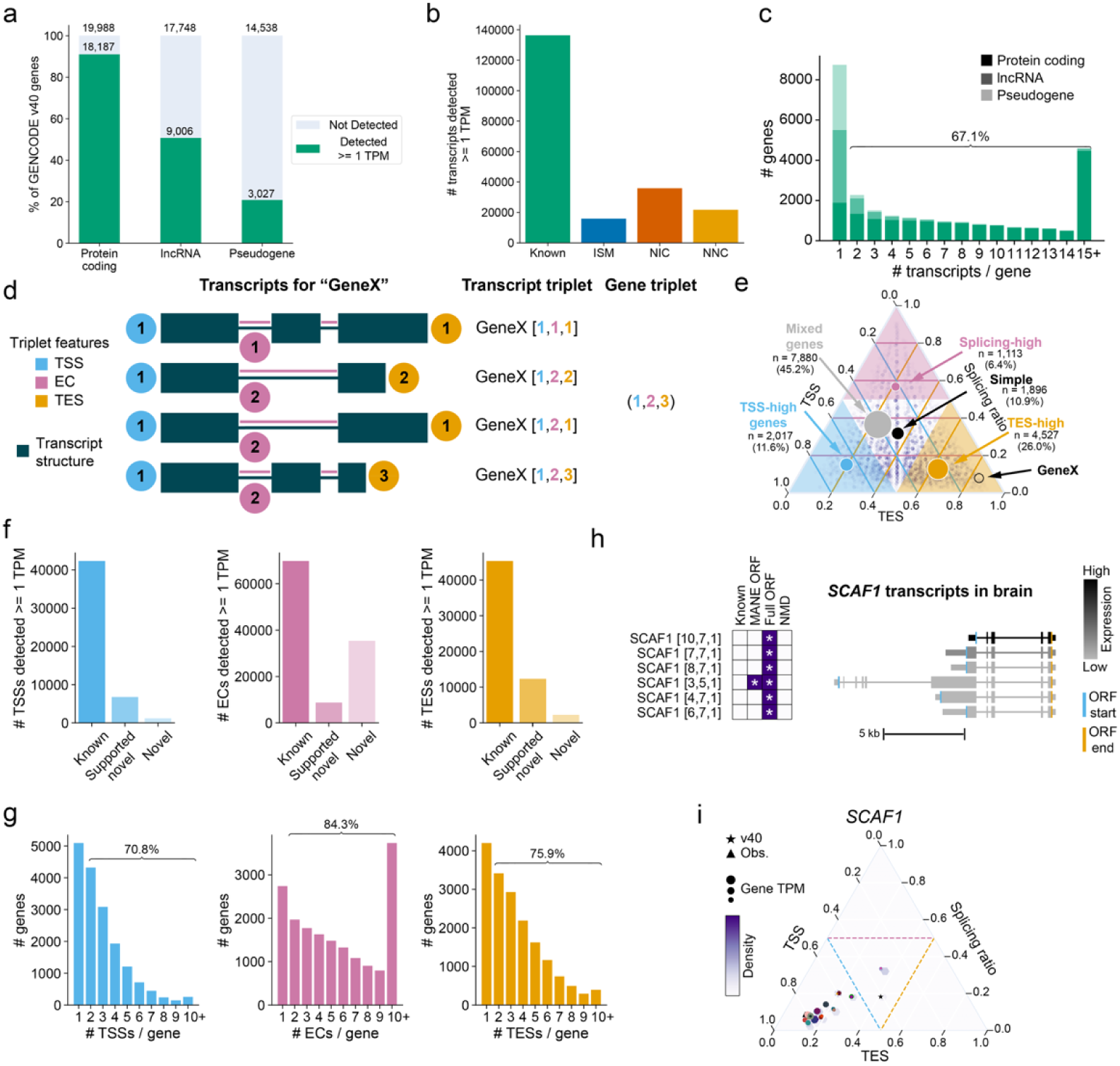
Drivers of transcriptional diversity. **a,** Percentage of GENCODE v40 genes by biotype detected (>= 1 TPM) in at least one human long read RNA-seq condition. **b,** Novelty characterization of detected in human long read RNA-seq. **c,** Transcripts per polyA gene (lncRNAs, pseudogenes, protein coding genes) detected (>= 1 TPM) in human ENCODE long read RNA-seq. **d,** Representation of structure and transcript / gene triplet convention for 4 different transcripts from the same gene based on the transcript start site (TSS), exon junction chain (EC), and transcript end site (TES) used. **e,** Layout of the gene structure simplex. Number of human protein-coding genes per sector indicated in density and in circles. GeneX coordinates from **d** also shown. Proportion of TSS usage is the blue axis (left), proportion of TES is the orange axis (bottom), and proportion of splicing ratio is the pink axis (right). Regions of the simplex are colored and labeled based on their sector category (TSS-high, splicing-high, TES-high). **f,** From left to right: TSSs, ECs, or TESs >= 1 TPM in human long-read RNA-seq from protein coding genes. Known features are annotated in GENCODE v29 or v40. Novel supported features are supported by the following: TSS: CAGE, RAMPAGE, PRO-cap, CA-H3K4me3 ENCODE cCREs, Pol II ChIP-seq, or long-read GTEx TSSs; EC: long-read GTEx ECs; TESs: PAS-seq, the PolyA Atlas, or long-read GTEx TESs. **g,** From left to right: Number of TSSs, ECs, or TESs >= 1 TPM per protein coding gene in human long read RNA-seq for e, TSSs, f, ECs, g, TESs. **h,** Gene structure simplex for *SCAF1*, where each colored point represents a gene triplet in a condition. Density of points also shown. **i,** Transcripts of *SCAF1* detected (>= 1 TPM) in brain.

We categorized and visualized each transcript according to its transcript start site (TSS), exon junction chain (EC), and transcript end site (TES) (Fig. 5d-e), and created a transcript visualization interface for those results (<u>Long Read RNA-seq Data Viewer</u>). Most protein coding genes have multiple TSSs, ECs, and TESs (Fig. 5f-g). For 44% of protein-coding genes, transcriptional diversity is biased towards variation specifically in either the TSS, EC, or TES (Fig. 5e).^72^ For example, transcripts of the splicing factor *SCAF1* preferentially differ based on TSS usage (Fig. 5h-i). Together, these results indicate constraints within transcripts for certain protein coding genes.

#### A catalog of RNA turnover rates across human genes

RNA is degraded at variable rates, impacting steady-state gene expression and isoform abundance.^74,75^ Variation in isoform stability has also been linked to disease.^76^ To measure RNA stability genome-wide, we used RNA Bru-seq and BruChase-seq assays across 16 widely-used cell lines that were coordinately grown, distributed, and profiled at the same time points by several additional types of experimental assays (<u>Deeply</u> <u>Profiled Cell Line Datasets</u>).^77,78^ We measured RNA degradation shortly after transcription (0 – 2 h) and over a later time interval (2 – 6 hr) (Fig. 6a-b).^79^ Nascent RNA sequencing reads mapped to ∼20% of the genome in individual cell lines, and to ∼48% of the genome collectively across all 16 cell lines.^37^

**Figure 6.**
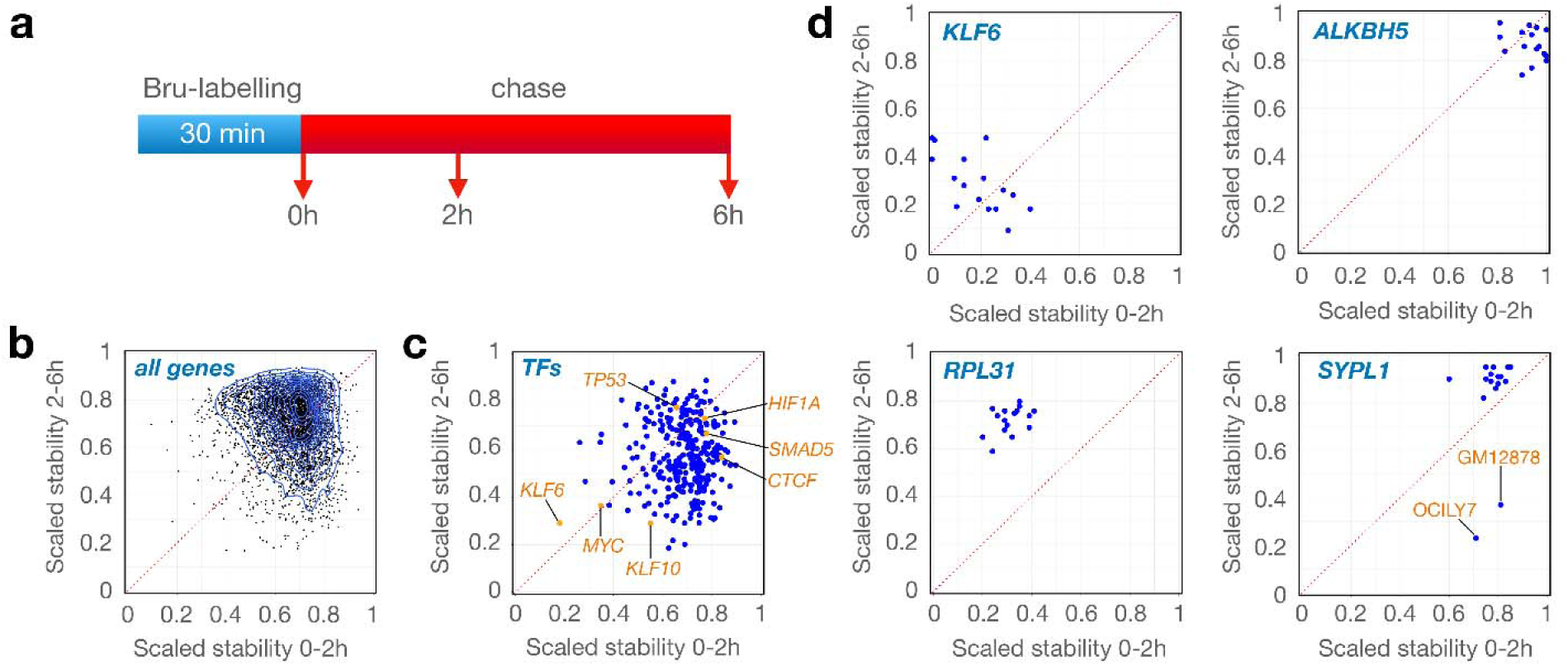
Stability of transcripts across time and cell models. **a,** Experimental outline of the Bru-seq and BruChase-seq techniques. **b**, Scaled stability scores for the 0-2 h period (x-axis) and for the 2-6 h period (Y-axis) for transcripts from 4415 genes expressed >0.5 RPKM across all 16 cell lines. **c,** Stability scores of transcripts coding for transcription factors with each dot representing an average degradation score across 16 cell lines. **d,** Examples of transcripts with exceptionally low stability (KLF6), high stability (ALKBH5), low initial stability but then high stability (RPL31) and cell type-specific regulation of stability (SYPL1).

RNA turnover rates varied widely across genes, time, and cell types and particularly among transcription factors (Fig. 6c). Of those, *KLF6* and *ALKBH5* are consistently among the most unstable and stable transcripts, respectively, across both time and cell lines (Fig. 6d). Some genes showed consistent patterns across cell types, such as the ribosomal protein transcript *RPL31* that degrades rapidly and specifically in early time points. Others such as *SYPL1* were highly variable between cell types (Fig. 6d). More generally, isoforms from the same gene often had different stability, indicating prevalent post-transcriptional isoform selection.^46^ Genetic factors also likely contribute to some of that variation in RNA turnover rates.^76,80^

#### Binding and functional profiles for human RNA binding proteins

As a step towards understanding how RNA binding proteins contribute to transcript life cycles, we systematically characterized their interactions with RNA using eCLIP and their functional effects using gene knockdowns.^81^ In total, the Encyclopedia includes eCLIP profiles for 286 RNA binding proteins in at least one cell line (<u>eCLIP Datasets</u>) and RNA-seq after knockdown for 536 (<u>RBP Knockdown RNA-seq Datasets</u>). For 216 RNA binding proteins, we generated both an eCLIP experiment and a corresponding knockdown experiment, allowing for both RNA binding and functional to be queried. The final ENCODE dataset includes a deep investigation of >100 zinc-finger proteins that revealed substantially altered splicing and polyadenylation site usage, as well as novel multi-functional RNA binding proteins.^82^ Deep-learning models on eCLIP profiles can be used to prioritize disease variants and uncover RNA binding protein-related mechanisms of pathogenesis.^83^

### Interactions among regulatory elements and genes

The third major focus of ENCODE 4 was mapping the physical and functional interactions among genes and gene regulatory elements.

#### Mapping physical interactions genome-wide at element and motif resolution

To map physical interactions genome-wide, ENCODE sequenced DNA:DNA proximity-ligation libraries generated via a variety of protocols. Together, those datasets expand upon previous consortium efforts by many-fold across a range of metrics. For example, the Encyclopedia contains 4-fold more sequenced molecules from proximity-ligation libraries, including 39-fold more that were generated using non-sequence-specific nucleases which enable higher resolution^84–86^.

For high-resolution physical interaction mapping, we developed intact Hi-C (Fig. 7a).^84^ We used intact Hi-C to create an atlas of physical interactions across 100 human cell and tissue types. These included high-density physical interaction maps comprising ∼90 billion sequencing reads each from reference cell lines (K562, HepG2, HCT-116) (Fig. 7b), and ∼350 billion sequencing reads across 40 lymphoblastoid cell lines (LCLs) derived from 40 donors (Fig. 7c) (<u>LCL Intact Hi-C Datasets</u>). High-density maps resolve the endpoints of long-range interactions to within 10 bp – sufficient to identify interactions between individual regulatory elements – and detect most loops shorter than 250 kb. Nearly all anchors overlap an ENCODE DHS. Maps from different donors allowed us to quantify allele-specific physical interactions in LCLs, and to identify ∼1,000,000 point-to-point loops connecting 138,049 distinct loop anchors (<u>Combined</u> <u>LCL Intact Hi-C Map</u>). We also resolved homolog-specific interactions by phasing genetic variants across chromosome-length haploblocks (<u>Phased Genotype Datasets</u>), enabling us to quantify allele-specific physical interactions for millions of single nucleotide polymorphisms and to relate allele-specific interactions to transcription factor binding motifs (Fig. 7c) (Case Study 4).

**Figure 7.**
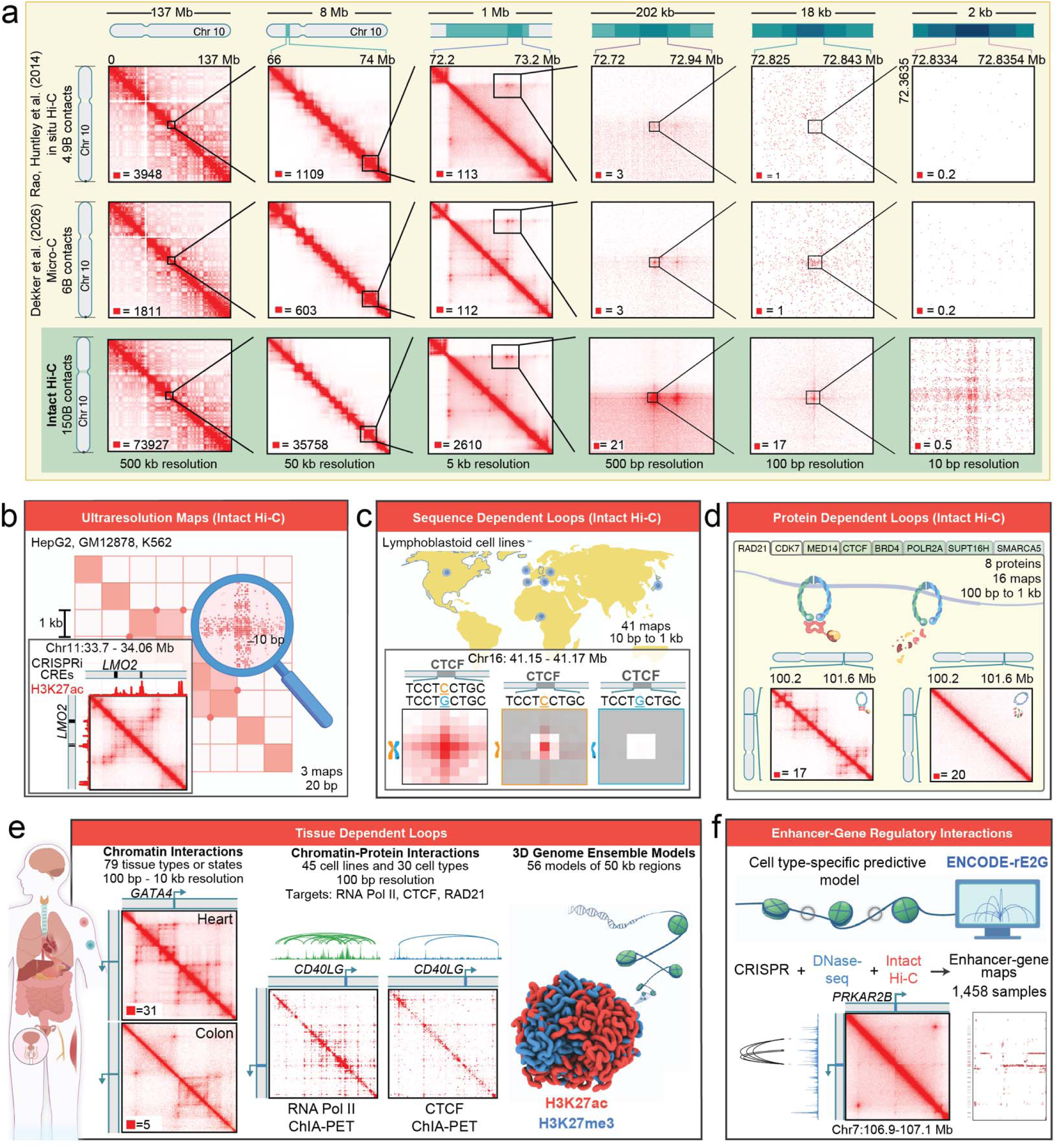
High resolution maps of physical interactions. **a,** We compare our Intact Hi-C contact map at chromosome 10 in lymphoblastoid cells (bottom row, ∼150 B contacts) to other extant datasets: In situ Hi-C (top row, ∼4.9 B contacts);^126^ the 4D Nucleome’s deepest 3D map, based on Micro-C (middle row, ∼6 B contacts).^127^ By contrast with prior methods, intact Hi-C resolves physical interactions at sub-element (<100 bp) and motif level resolutions (10 bp) not seen in prior maps. **b,** Ultradeep intact Hi-C maps (<20 bp) in HepG2, K562, and HCT-116 resolve physical interactions between individual CRISPRi-validated regulatory elements (inset: a contact map for the LMO2 locus in K562 cells is shown, along with corresponding H3K27ac ChIP-Seq data and CRISPRi functional CREs for the LMO2 promoter). **c**, We generated and phased intact-HiC maps from 41 LCLs to generate haplotype-specific contact matrices and probe allele-specific nuclear architecture. Inset: a single SNP on chromosome 9 (41.15–41.17 Mb) disrupts a CTCF motif, causing loss of looping. **d.** We identified protein-dependent loops by auxin-inducible degradation of eight proteins. Contact matrices for a representative locus (chromosome 10: 100.2–101.6 Mb) before (left) and after (right) RAD21 degradation are shown, with an observed loss of loop domains. **e,** Tissue specific interactions were studied with three approaches. Left: With intact Hi-C, we assayed 79 different human tissues, including tissue-specific physical interactions. Contact matrices at the *GATA4* locus (chromosome 8: 11.14–12.03 Mb) from heart and colon show several heart-specific loops to the *GATA4* promoter. Middle: ChIA-PET in 45 cell lines and 30 primary cell types (134 maps, 100 bp) reveals cell-type-specific interactions mediated by POLR2A, CTCF, and RAD21. Right: Ensemble models of genome organization reveal the 3D compaction of epigenetic marks. **f,** ENCODE-rE2G predictions integrating CRISPRi, DNase-seq, and Intact Hi-C across 369 samples reveals networks of regulatory and physical interactions.

#### Protein-associated and protein-dependent physical interactions

To assess the relationship between physical interactions and specific chromatin associated proteins, we measured the effect of degrading eight major regulatory factors (RNA Polymerase II, RAD21, CTCF, CDK-7, SUPT16H, BRD4, MED14, and SMARCA5) using an auxin-inducible degron system in HCT-116 human colorectal carcinoma cells (Fig. 7d) (<u>Targeted Degradation Datasets</u>). We report that, while many loops depend on cohesin (85%) and CTCF (78%), a fraction do not. Degradation of RNA Pol II effaces loops anchored at RNA Pol II binding sites (26% lost) more than those anchored at CTCF (11% lost), and those RNA Pol II loops were often strongly dependent on cohesin. Together, these results indicated that recruiting RNA Pol II gives an element the ability to communicate at a distance via cohesin extrusion. In contrast, loops anchored at AP1 motifs are more robust to degradation of both CTCF and cohesin, suggesting there are additional mechanisms of loop formation.

#### A broad map of tissue-dependent physical interactions

We also created a catalog of physical interactions maps spanning >100 human cell and tissue types using intact Hi-C and ChIA-PET each in ∼80 cell or tissue types (Fig. 7e) (<u>Physical Interaction Datasets</u>). The intact Hi-C maps revealed millions of physical interactions between individual DNA elements, while the ChIA-PET maps revealed context-specific physical interactions associated with CTCF and RNA Pol II.^87–89^ We also created reference intact Hi-C maps for organs and developmental lineages, we aggregated results from the component cells and tissues (<u>Intact Hi-C Body Map</u> <u>Datasets</u>). For instance, we created a heart-specific Hi-C map containing 34 billion contacts, spanning data from multiple individuals and all four of the heart’s chambers. Together, the catalog of physical interactions across all cell and tissue samples includes 11 million loops at <100 bp resolution, sufficient to resolve physical interactions between individual DNA elements in primary tissues. Finally, for a subset of genomic loci, we created ensemble models of genome architecture at specific loci using ChIP-seq data.^90,91^

#### Regulatory interactions are associated with physical interactions in the human genome

A common hypothesis is that long-range regulatory interactions correspond to physical interactions between regulatory elements, although generalized experimental evidence is lacking. We thus compared Hi-C physical interactions with element-to-gene regulatory interactions mapped using CRISPR perturbations. We found that most long-range regulatory interactions correspond to physical interactions. For instance, of the regulatory interactions detected in full tiling CRISPRi screening experiments in K562,^45^ up to 92% of perturbations with regulatory impact corresponded to physical loops (Supplementary Figure 10). In a similar tiling experiment that only targeted DHSs in HCT-116, 100% of 30 mapped regulatory interactions corresponded to physical loops, whereas pairs of loci not found to exhibit regulatory interactions corresponded to physical interactions only 30% of the time.^84,92^ Together, these results strongly support a model in which physical interactions among regulatory elements are critical contributors to their regulatory function.

#### Measuring and predicting regulatory interactions between enhancers and promoters

To evaluate more broadly the regulatory function of DNA elements, we developed a model (ENCODE-rE2G) of predicted functional regulatory interactions genome-wide and across multiple cell types.^93^ We trained ENCODE-rE2G on 10,356 element-gene pairs, including 471 experimentally verified regulatory connections that had been assayed with CRISPR screens in ENCODE and in external studies.^44,93^ We then predicted regulatory interactions genome-wide and in additional cell types using a series of supervised classifiers. Overall, 75% of the predicted regulatory interactions occurred within 100 kb of the target promoter. On average, the ENCODE-rE2G model predicted 5.9 regulatory interactions per gene, and 1.6 regulatory interactions per distal regulatory element. Based on the ENCODE-rE2G models, we predicted and validated in specific cases (i) that the promoters of housekeeping genes are less sensitive to distal regulatory elements^49^ and (ii) that groups of enhancers located in close physical proximity often interact super-additively.^44,93^ We extended ENCODE-rE2G to predict regulatory interactions in the absence of physical interaction studies by using 1,458 of the 4,159 ENCODE DNase-seq experiments to predict >92 million regulatory interactions (<u>ENCODE-rE2G Predictions</u>).

#### Integrative biological insights using ENCODE data

Together, the ENCODE datasets can provide detailed insights into gene regulatory elements, gene expression, and how regulatory elements and genes interact. To illustrate this potential in the context of dynamic cell systems, we provide case studies showing how ENCODE data can be integrated to provide insights into intestinal crypt cell differentiation (Case Study 5), B-cell trans-differentiation (Case Study 6), and kidney cell differentiation (Case Study 7).

### Gene regulation in mouse and across evolution

Mice are an invaluable model organism, enabling studies that are not possible in humans. Therefore, the annotation of mouse genes, regulatory elements, and the interactions between them was a fourth key focus of ENCODE.

#### Mapping regulatory elements in mouse using chromatin accessibility

The final phase of ENCODE expanded mapping of DHSs in mouse cell and tissue types by ∼15-fold over MouseENCODE.^6^ Mouse DHSs were mapped in 578 samples from distinct cell and tissue types and states or ages of which 114 represent fetal or embryonic states and 448 represent post-natal states (<u>Mouse DNase-seq Datasets</u>). Collectively, these maps yielded an index of 2.6 million mouse DHSs. Based on the relative sizes of the mouse and human genomes and densities of DHSs in each, we estimate that the mouse index is ∼61% complete. ∼1 million mouse elements from 367 samples (∼37% of annotated DHSs) were categorized using the cCRE framework.

#### Gene regulatory dynamics across mouse postnatal development

We focused on changes in gene regulation during mouse postnatal development, extending previous ENCODE studies of prenatal development. To enable allele specific and parent of origin analyses, we used C57BL6/J x CAST/EiJ F1 hybrid mice (Fig. 8a). We analyzed seven time points representing childhood (post-natal days 4, 10, 14 and 25), puberty (post-natal day 36), young adulthood (2 months), and senior adulthood (18 months). We mapped chromatin accessibility by DNase-seq across these time points in 13 different tissues. We also profiled gene expression by bulk and single-cell RNA-seq in five core tissues (adrenal gland, heart, skeletal muscle, left cerebral cortex, and hippocampus) across the same time intervals. We additionally mapped single cell chromatin accessibility at post-natal day 14 and 2 months; and CTCF occupancy plus the H3K27ac and H3K4me3 covalent histone modifications at 2 months (Fig. 8a). All post-natal experiments were performed on equivalent time point samples from four animals, two from each sex.

**Figure 8.**
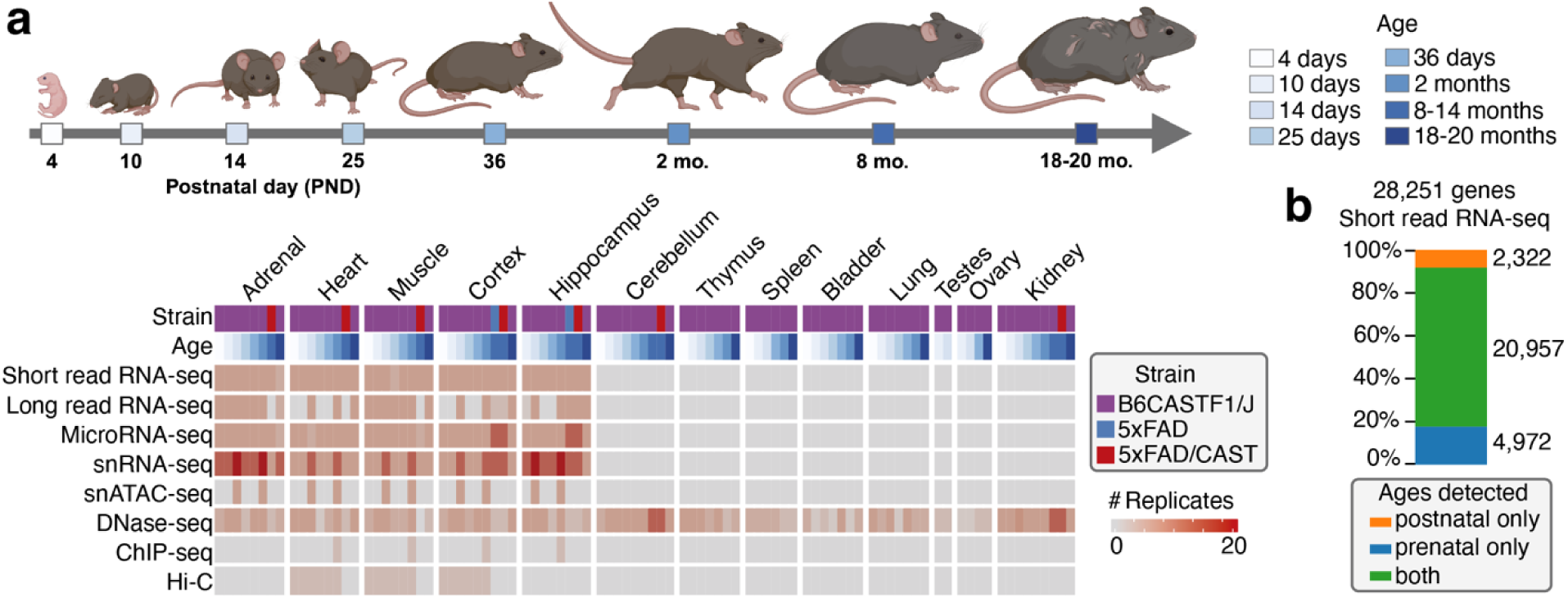
Changes in gene regulation across mouse postnatal development. **a,** Postnatal mouse time course spanning juvenile (PND4–PND25), puberty (PND36), and adult stages (2 mo., 8 mo., 18–20 mo.) and data types collected in a wide variety of tissues including adrenal gland, heart, skeletal muscle, cerebral cortex, hippocampus, thymus, spleen, bladder, lung, gonads, and kidney at various time points. **b,** Out of 28,251 genes detected in the short-read RNA-seq data, 2,322 genes are only expressed postnatally.

In the five core tissues, 4,982 (18%) of 28,251 genes were selectively expressed prenatally, and 2,322 (8%) genes were selectively expressed after birth (Fig. 8b). Long read RNA-seq data revealed many instances of isoform switching between biological contexts, an example of which is detailed in Case Study 8. We also identified cell-type specific regulatory elements from pseudobulked single nucleus ATAC-seq (Supplementary Figure 11).

#### Conservation of regulatory elements between human and mouse

Genome sequence similarity between species is commonly used to estimate evolutionary conservation in protein-coding regions. Extending that principle to gene regulatory elements has been challenging because the corresponding DNA sequences mutate rapidly and the relationship between sequence and function is less clear. We identified about 550,000 DHS in both species with evidence of sequence conservation. That represents 10% of human DHS and 21% of mouse DHS, noting that the mouse DHS annotation is approximately 61% complete.^13^

As an alternative approach, we evaluated promoter-proximal functional conservation between human and mouse using computationally matched DNase, ATAC-seq, and histone ChIP-seq datasets (Supplementary Figure 12). We completed this analysis in two complementary ways. We tested for functional conservation between 2.2 million human-mouse regulatory element pairs matched based on sequence similarity. We also evaluated functional conservation across the regions flanking the promoters of orthologous genes. For that analysis, we tested ∼180 million pairs of elements (Fig. 9a) (<u>Functional Conservation Analyses</u>). To estimate functional conservation, we then tested if each human-mouse regulatory element pair had a similar pattern of activity across matched datasets (Fig. 9b, Supplementary Figure 12a). We identified 300,000 (FDR ≤ 10%) and 3.1 million (FDR ≤ 25%) pairs of elements from the core and extended set, respectively, with evidence of functional conservation. Those pairs cover 20% of the human DHSs that were included in this analysis. Pairs of elements with conserved constitutive activity were enriched in promoter-like and proximal enhancer-like cCRE annotations, whereas pairs of regulatory elements with conserved tissue-specific activity were enriched in distal enhancer-like cCREs. Many pairs of sequence-matched elements with functional conservation were only annotated as cCREs in human. Transferring those annotations to mouse identified additional putative regulatory elements supported by orthogonal activity signatures (Supplementary Figure 12b-c). In addition, transferring orthologous loop anchors from human to mouse resulted in loop predictions with ∼9-fold higher validation rates than unprioritized loops (Fig. 9c). We also show that functional conservation can support interpretation of noncoding variants associated with human traits and disease (Fig. 9d-e).^94^

**Figure 9.**
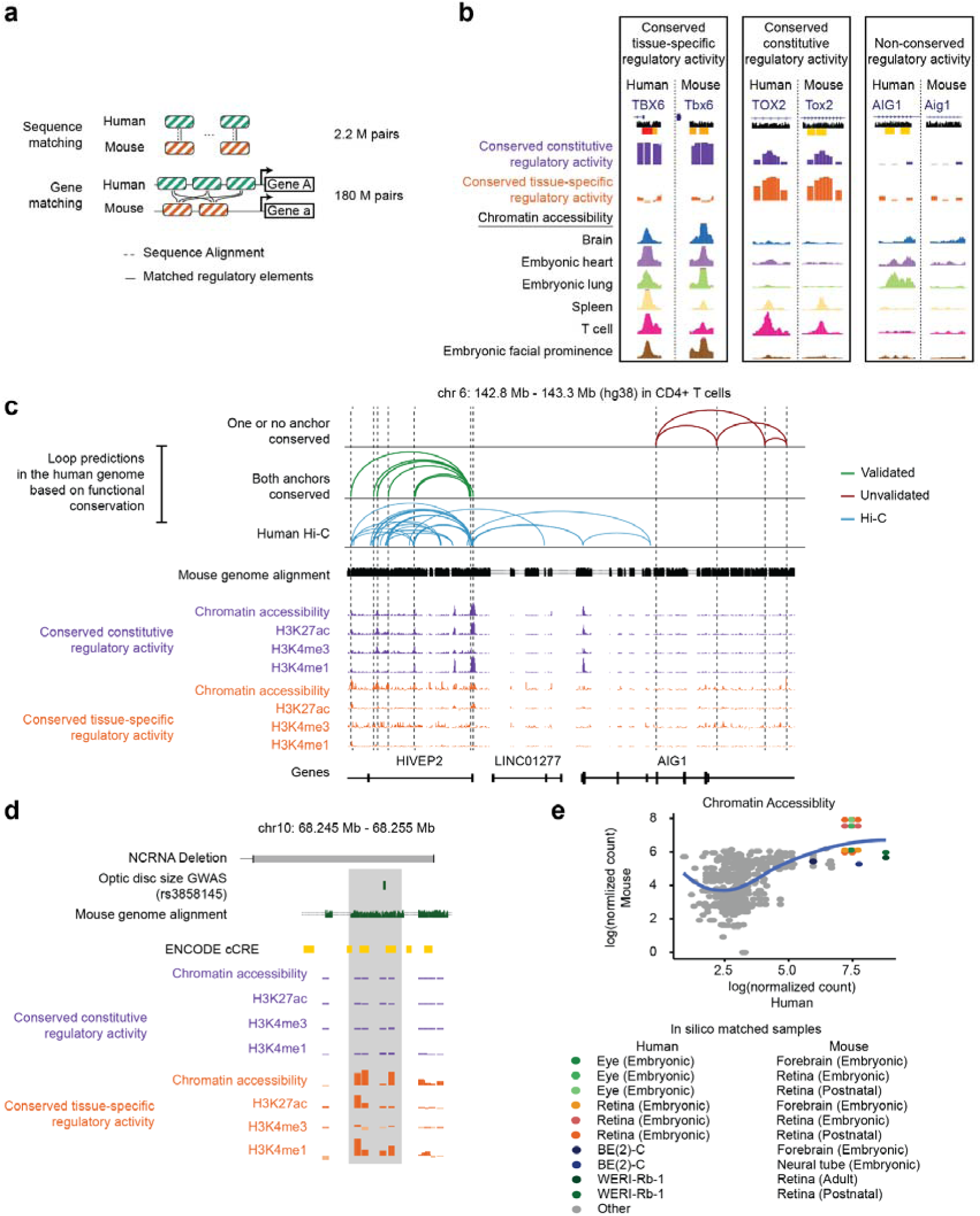
Functional conservation of gene regulatory elements between human and mouse. **a,** Overview of our functional conservation analysis approach. A core set of 2.2 M regulatory element pairs was defined by sequence homology and an extended set of 180 M by gene homology. DNase-seq, ATAC-seq and ChIP-seq profiles were co-embedded into a shared low-dimensional space to generate in silico-matched samples. **b,** Genome browser examples of human-mouse regulatory element pairs (REPs) showing conserved tissue-specific accessibility (TOX2), conserved constitutive chromatin accessibility (TBX6), or no functional conservation (AIG1). **c.** Human chromatin loops from in situ Hi-C (bottom) and loops predicted by transfer from mouse ENCODE Hi-C (top, middle) in CD4⁺ T cells. Examples with one or no conserved anchors (top) versus two conserved anchors (middle) are shown. Green loops are validated by Hi-C, whereas red loops are not. Conservation-based predictions exhibit higher validation rates, highlighting the value of conservation for improving cross-species transfer of Hi-C loop annotations **d,** Human enhancer containing a variant linked to congenital retinal nonattachment. The element has high functional conservation scores of tissue-specific activity between species for chromatin accessibility, H3K27ac and H3K4me1; reporter assays in mice from a previous study confirm retinal regulatory activity.^128^ **e,** Chromatin accessibility signals for the element highlighted in (**d**) across *in silico* matched human and mouse samples demonstrate conserved retinal accessibility. Retina-related samples are highlighted by color.

We also modeled regulatory element evolution at a subset of DHSs by training promoter and enhancer gkm-SVM sequence models of 1,644 and 244 human and mouse chromatin accessibility datasets (<u>gkm-SVM Models</u>). Across 45 pairs of matched human and mouse samples, we observed rapid evolution of enhancers at the DNA sequence level and high variation in enhancer conservation rate by tissue type, even though transcription factors represented in the sequence models are well-matched in all human and mouse tissue pairs.^95^

#### Contribution of transposable elements to the evolution of regulatory elements

Transposable elements are a substantial evolutionary source of regulatory elements in the human and mouse genomes.^96–104^ Based on ENCODE 4 annotations, 35% of DHSs and ∼33% of cCRE-categorized elements substantially overlap transposable elements. Overlapping elements have similar reporter-assay activity, patterns of covalent histone modifications, and enrichment for genetic associations as elements that do not overlap transposable elements.^104,105^ We estimate that 37% of human-lineage specific cCREs and 28% of mouse-lineage-specific cCREs overlap transposable elements. Transcription factor binding motifs in those elements are typically ancestral, with the notable exception of short interspersed nuclear elements.

#### Chromatin loops over development and evolution

Finally, we sought to evaluate the evolution of human nuclear architecture more broadly. To that end, we developed a variant of Hi-C, PaleoHi-C, for use with ancient samples;^106^ and demonstrated that the chromatin loops in human skin are evident in a 52,000-year-old skin sample from a woolly mammoth. We also measured physical interactions across bilaterian evolution via analyses of seven different organisms (human, mouse, sea lamprey, arctic lamprey, sea urchin, ciona, and fruit fly) (Supplementary Figure 13) (Supplementary Table 11). Patterns of looping within the vertebrate clade are consistent with cohesin-mediated extrusion arrested by CTCF in an orientation-specific fashion. However, this phenomenon is not seen outside the vertebrates (i.e., in sea urchin, ciona, and fruit fly), consistent with observations of a recent study.^107^

## Accessing the Encyclopedia of DNA Elements

The Encyclopedia of DNA Elements is available in multiple forms: reference datasets, visualization tools for exploration by human readers, and via Model Context Protocol (MCP) servers to facilitate exploration via artificial intelligence agents.

Primary experimental results, integrative analyses, and computational predictions are available at the ENCODE Portal (https://www.encodeproject.org/).^108^ The portal enables searching and filtering via structured metadata associated with every dataset from all phases of the ENCODE project. The portal also provides links to specific sample collections of broad interest.

Visual investigation of human and mouse gene regulation is facilitated by several resources. The ENCODE DHS catalog can be explored via a web-based tool (https://www.regulome.org). An expanded version of the SCREEN tool (https://screen.wenglab.org/) enables users to search genes, cCRE annotations, and the underlying experimental datasets. SCREEN also enables users to search for elements in specific cell or tissue types and compare the compendium of elements found in any cell or tissue.^24^ Factorbook (https://www.factorbook.org/) provides an ENCODE-integrated catalog of transcription factor binding specificities and regulatory roles. Factorbook also includes a comprehensive collection of motifs and candidate transcription factor binding sites derived from large-scale ChIP-seq data. A UC Santa Cruz Genome Browser Track Hub facilitates browsing ENCODE data and analyses (<u>UCSC Browser Track Hub</u>).^109^ We also developed applications to visualize and query functional conservation scores (<u>Functional Conservation Query Tool</u>, <u>Functional</u> <u>Conservation Track Hub</u>), transcript isoform structure (<u>Long Read RNA-seq Data</u> <u>Viewer</u>),^72^ physical interactions in the genome (<u>Juicebox</u>),^110^ and three-dimensional models of genome structure (<u>Spacewalk</u>).^111^

Finally, to facilitate the use of artificial intelligence systems to explore ENCODE data, we developed MCP servers for both the broad ENCODE dataset (ENCODE MCP Server) and for physical interaction datasets (Juicebox MCP Server).

## Discussion

The Encyclopedia of DNA Elements was created as a public resource to support biomedical research. ENCODE is the product of hundreds of investigators working together across disciplines towards the common vision of understanding where functional elements are encoded in the human genome and how they are used. The Encyclopedia includes thousands of assays and analyses that are standardized, organized, and documented to facilitate broad use. ENCODE and public resources like it have proven to be valuable guides to novel mechanistic studies of the human genome.^112–116^ ENCODE also provides essential training data for machine learning models in biology^117–124^ which are sure to expand in the coming years.

Given ENCODE’s stated goal of identifying and characterizing all the functional elements in the human genome, it is natural to consider the question of completeness: in terms of genes, in terms of regulatory elements, and in terms of the interactions between them.

For genes, we currently have confidence that there are between 19,000 and 20,000 protein-coding genes in the human genome.^125^ That estimate has not grown appreciably despite thousands of gene expression studies. However, ENCODE 4 and other studies have identified many thousands of novel transcripts and lncRNAs as well as substantial variability in transcript stability that remains to be understood. Although the pace of annotating new transcript isoforms and non-coding RNAs may be moderating, abundant opportunities for discovery remain.

For regulatory elements, several lines of evidence indicate that the human genome encodes approximately 5.3 million accessible chromatin elements across all tissues and cell types. To annotate the potential function of those elements, we also generated detailed maps of histone modifications, transcription factor binding, and nascent transcription across diverse samples, and integrated those data with computational models. Intriguingly, large-scale functional studies reported here demonstrate that similar combinations of transcription factors and flanking histone modifications are associated with disparate effects on target gene expression. We expect that a better understanding of how regulatory elements interact will be a key to resolving that discrepancy.

For physical interactions, a measure of completeness has been achieved for a few cell types such as B-lymphoblastoid cells. In those, we now have sufficient resolution to detect interactions between individual regulatory elements, and the number of detected physical interactions over distances <250 kb appears to have plateaued after hundreds of billions of sequencing reads. Those maps reveal an extensive and complex network of regulatory elements, the importance of which we are only beginning to understand. Expanding high-resolution physical interaction maps to a broader range of biological contexts and developing a conceptual understanding of how those interactions impact gene regulation remain important challenges.

Taken together, the Encyclopedia of DNA Elements is a foundational resource for studies of the human and mouse genomes and the role of gene regulation in traits and disease. Throughout the development of that resource, many questions have emerged about how DNA elements function alone and in combination to drive transcription. Indeed, “the more we learn about the human genome, the more there is to explore.”^1^

## Methods

Detailed methods for each experiment and the associated primary analyses are available on the ENCODE portal (https://www.encodeproject.org/). The information can be accessed via a unique accession number for each experiment. The information includes standardized information about the source and preparation of the biological sample; detailed experimental and analytical protocols; quality control reports; raw sequencing data; and primary processed data such as gene expression quantification, peak calls, and genome browser tracks (<u>UCSC Browser Track Hub</u>). In some cases, experiments are further organized into series to represent more complex experimental designs. Those series include developmental and differentiation time courses, treatment time and dose courses, environmental or genetic perturbations; and collections of disease versus control. Additional methods can be found in companion papers referenced in each section.

## Author Contributions

### ENCODE Consortium Contributions

#### Data Production Leads^†^

Gabriela Balderrama-Guttierez, Charles Epstein, Eva Gega, Dimos Gkountaroulis, Jessica M. Halow, Mark Mackiewicz, Brian Magnuson, Ragini Mahajan, E. Christopher Partridge, Michelle T. Paulsen, Rajinder K. Kaul, Suhas S. P. Rao, Richard Sandstrom, Muhammad S. Shamim, Derya Unutmaz, Ishwarya Venkata Narayanan, Annika Weimer, Brian Williams

#### Functional Characterization Leads^†^

Vikram Agarwal, You Chen, Ilias Georgakopoulos-Soares, Fumitaka Inoue, Martin Kircher, Betty Liu, Georgi Marinov, Eyal Metzl-Raz, Mauricio I. Paramo, Len A. Pennacchio, Steven K. Reilly, Xingjie Ren, Sagar R. Shah, Manman Shi, Zohar Shipony, Keith Siklenka, Ryan Tewhey, Nate Tippens, Josh Tycko, Axel Visel, Xiaoyu Yang, David Yao, Li Yao, Junke Zhang

#### Analysis Center Leads^†^

Michael A. Beer, Jason D. Chobirko, Ziwei Chen, Kushal K. Dey, Alan Y. Du, Ivy Evergreen, Ting Fu, Vivian Hecht, Elaine Huang, Alireza Karbalayghareh, Daniel S. Kim, Eran Kotler, Soumya Kundu, Johannes Linder, Kristy S. Mualim, Anusri Pampari, Aman Patel, Giovanni Quinones-Valdez, Vivekanandan Ramalingam, Austin Wang, Chang M. Yun, Xiaoyu Zhuo

#### Working Group and Subgroup Leads

ENCODE established working groups (i) to coordinate data collection and analysis both across the consortium and for specific types of assays, and (ii) to improve visibility and usability of the Encyclopedia. All working groups and subgroups are listed in alphabetical order.

##### • Analysis Working Group

***The Analysis Working Group coordinated integrative analyses across the consortium and provided a forum for developing, refining, and evaluating computational approaches used in the flagship study. The group reviewed analytical results, contributed biological and technical interpretation across data types, and helped synthesize subgroup findings into the broader ENCODE resource and manuscript.***

Nadav Ahituv, Mark Gerstein, Christina S. Leslie, Michael P. Snyder, Zhiping Weng

##### • Biosamples Coordination Working Group

***The Biosamples Coordination Working Group organized the collection and sharing of samples across ENCODE; coordinated assays on common samples; and organized and assembled metadata on ENCODE samples.***

Nina P. Farrell, Elizabeth Gaskell, Eva Gega, Rajinder K. Kaul, Mats Ljungman, Brian A. Williams, Annika Weimer

##### • Functional Characterization Working Group

***The Functional Characterization Working Group coordinated targets for functional characterization assays and developed analyses to compare results across those assays.***

Keith Siklenka, Galip Gürkan Yardımcı

##### • Mouse Working Group

***The Mouse Working Group organized collection and sharing of mouse samples and developed cross species analyses.***

Aviva Presser Aiden, Gabriela Balderrama-Guttierez, M. A. Bender, Mark Groudine, Jessica M. Halow, Hongkai Ji, Rajinder K. Kaul, Shaiza Pasha, Elisabeth Rebboah, Fairlie Reese, Narges Rezaie, Diane Trout, Brian Williams, Barbara J. Wold, Ali Mortazavi

##### • Nuclear Architecture Working Group

***The Nuclear Architecture Working Group coordinated assays and analyses of the physical interactions in the human and mouse genomes.***

Erez Lieberman Aiden, William S. Noble

**3D Tracing Subgroup:** Dimos Gkountaroulis, Anat G. Vivante, Galip Gürkan Yardımcı, Guy Nir

**Body Atlas Subgroup:** Ragini Mahajan, Muhammad S. Shamim

**ChIA-PET Subgroup:** Eva Gega

**Diploid Analysis Subgroup:** Muhammad S. Shamim, Olga Dudchenko

**Element to Gene Subgroup:** Andreas R. Gschwind, Kristy S. Mualim, Maya U. Sheth, Jesse M. Engreitz

**Intact Hi-C Subgroup:** Suhas S. P. Rao, Huiya Gu

**Modeling & Prediction Subgroup:** Ritambhara Singh, Gamze Gürsoy

**Physical Simulation Subgroup:** Vinícius G. Contessoto, Antonio B Oliveira Junior, Michele Di Pierro

**Ultra-Resolution Subgroup:** Suhas S. P. Rao, M. Jordan Rowley, Ragini Mahajan

**ZooCode Subgroup:** Olga Dudchenko

##### • Outreach Working Group

***The Outreach Working Group created resources to improve community awareness and use of ENCODE data, including via public ENCODE Users meetings.***

Jennifer Jou, Jason Hilton, Meenakshi S. Kagda, Jill E. Moore, Richard Sandstrom

##### • Personalized Genomes Working Group†

***The Personalized Genomes Working Group developed analyses to improve the assembly and analysis of human genome sequences using ENCODE data.***

Erez Lieberman Aiden, Mark Gerstein

**EN-TEx Subgroup:**

Beatrice Borsari, Olga Dudchenko, Charles B. Epstein, Timur Galeev, Jiahao Gao, Gamze Gürsoy, Tianxiao Li, Jason Liu, Ragini Mahajan, Joel Rozowsky, Kun Xiong, Jinrui Xu, Yucheng T. Yang

**Hi-C Subgroup:**

Olga Dudchenko, Gamze Gürsoy, Ragini Mahajan

^†^Authors are listed alphabetically by last name

##### • Registry of cCREs Working Group

***The Registry of cCREs Working Group developed a catalog of candidate cis regulatory elements from ENCODE data by applying a common set of criteria across ENCODE datasets.***

Jill E. Moore, Zhiping Weng

##### • RNA Working Group

***The RNA Working Group coordinated assays of gene expression and evaluated the adoption of emerging technologies such as long-read RNA-seq.***

Fairlie Reese, Brian A. Williams, Gabriela Balderrama-Gutierrez, Dana Wyman, Muhammed H. Çelik, Karan Bedi, Elisabeth Rebboah, Narges Rezaie, Diane Trout, Milad Razavi-Mohseni, Yunzhe Jiang, Beatrice Borsari, Mats Ljungman, Roderic Guigó, Barbara J. Wold, Ali Mortazavi

##### • Search and User Interface Working Group^†^

***The Search and User Interface Working Group worked with major data repositories including the Gene Expression Omnibus to enable interactive viewing and analysis of ENCODE datasets.***

Erez Lieberman Aiden, Anshul Kundaje

^†^Authors are listed alphabetically by last name

##### • Single Cell Analysis Working Group

***The Single Cell Analysis Working Group coordinated the collection and analysis of single cell assays of gene expression and chromatin accessibility.***

William J. Greenleaf, Anshul Kundaje

**Cell Type Annotation Subgroup:** Austin Wang, Julia M. Schaepe, Winston R. Becker, Yu Fu, Benjamin R. Doughty, Elisabeth Rebboah, Benjamin E. Parks, Eran Kotler, Salil S. Deshpande, Jennifer Jou, Fairlee Reese, Narges Rezaie, Meng Wang, Jill E. Moore, Idan Gabdank, Benjamin C. Hitz, Hongkai Ji, Michael P. Snyder, Zhiping Weng, Ali Mortazavi, William J. Greenleaf, Anshul Kundaje

**Pseudobulk and Integrative Analysis Subgroup:** Zhicheng Ji, Wenpin Hou, Austin Wang, Diane Trout, Anshul Kundaje

**Single-nucleus and Single-cell ATAC-seq Pipeline Subgroup:** Austin Wang, Surag Nair, Benjamin E. Parks, Caleb Lareau, William J. Greenleaf, Anshul Kundaje

**Single-nucleus and Single-cell RNA-seq Pipeline Subgroup:** Diane Trout, Ali Mortazavi

##### • Transcription Factor Coordination Working Group

***The Transcription Factor Coordination Working Group coordinated the prioritization of transcription factor tagging via ChIP-seq.***

Annika Weimer, Mark Mackiewicz

#### National Human Genome Research Institute Program Management^†^

***Sarah Anstice, Afia Asare, Stephanie Calluori, Elise A. Feingold, Daniel A. Gilchrist, Carolyn M. Hutter, Stephanie A. Morris, Michael J. Pazin, Ella K. Samer***

^†^Authors are listed alphabetically by last name

(The role of the National Human Genome Research Institute Program Management in the preparation of this paper was limited to coordination and scientific management of the ENCODE Consortium.)

#### Principal investigators^†^

Nadav Ahituv, Erez Lieberman Aiden, Michael C. Bassik, Michael A. Beer, Bradley E. Bernstein, J. Michael Cherry, Maria Ciofani, Barak Cohen, Gregory E. Crawford, Cedric Feschotte, Charles A. Gersbach, Mark Gerstein, William J. Greenleaf, Charles Lee, Christina S. Leslie, John T. Lis, Mats Ljungman, Eric M. Mendenhall, Jennifer R. Moran, Ali Mortazavi, Richard M. Myers, Len A. Pennacchio, Alkes Price, Jonathan K. Pritchard, Soumya Raychaudhuri, Timothy E. Reddy, Bing Ren, Yijun Ruan, Pardis C. Sabeti, Yin Shen, Jay Shendure, Michael P. Snyder, John A. Stamatoyannopoulos, Ryan Tewhey, Axel Visel, Ting Wang, Zhiping Weng, Barbara J. Wold, Xinshu Xiao, Haiyuan Yu’

^†^Authors are listed alphabetically by last name

### Manuscript Preparation

**Figure 1**

Eva Gega, Jill E. Moore

**Figure 2**

Sergey Abramov, Alexandr Boytsov

**Figure 3**

Vivekanandan Ramalingam, Chang M. Yun, Georgi Marinov, Vivian Hecht, Anusri Pampari, Aman Patel

**Figure 4**

Kushal K. Dey, Tabassum Fabiha, Berk Turhan, Yosuke Tanigawa, Xiaohe Tian, Charles E. Breeze

**Figure 5**

Fairlie Reese

**Figure 6**

Karan Bedi

**Figure 7**

Ragini Mahajan, Suhas S. P. Rao, Andreas R. Gschwind, Maya U. Sheth, Antonio B Oliveira Junior, Vinícius G. Contessoto, Xiaotao Wang

**Figure 8**

Elisabeth Rebboah

**Figure 9**

Weixiang Fang, Hongkai Ji

**Case Study 1**

Vivekanandan Ramalingam, Vivian Hecht, Anusri Pampari, Aman Patel, Ziwei Chen, Kelly Cochran

**Case Study 2**

Vivekanandan Ramalingam, Ziwei Chen, Johannes Linder, Ryan Tewhey

**Case Study 3**

Ivy Evergreen, Soumya Kundu, Tabassum Fabiha, Thahmina Ali, Kushal. K. Dey

**Case Study 4**

Ragini Mahajan, Suhas S. P. Rao

**Case Study 5**

Winston R. Becker, Vivian Hecht

**Case Study 6**

Beatrice Borsari, Silvia González-López, Sílvia Pérez-Lluch, Marina Ruiz-Romero

**Case Study 7**

Ryuji Morizane

**Case Study 8**

Elisabeth Rebboah, Fairlie Reese

#### Writing Group^†^

Erez Lieberman Aiden, Kushal K. Dey, William J. Greenleaf, Roderic Guigó, Benjamin C. Hitz, Hongkai Ji, Anshul Kundaje, Mats Ljungman, Jill E. Moore, Ali Mortazavi, Timothy E. Reddy, Michael P. Snyder, John A. Stamatoyannopoulos

^†^Authors are listed alphabetically by last name

### The ENCODE Project Consortium Author List

#### Mapping Centers

##### Altius Institute for Biomedical Sciences^†^

Aleksandra Abisheva, Sergei Abramov, Daniel Bates, Michael A. Bender, Jordan Bloom, Jennifer Bolin, Alexandr Boytsov, Madeline Brannon, Michael Buckley, Sergey Bushuev, Hua Cao, Andres Castillo, Allison J. Cote, Henry Cryst, Morgan Diegel, Siobhan Duffy, Douglas Dunn, Anas Fathul, Mark Frerker, Alister P. W. Funnell, Grigorios Georgolopoulos, Alexander A. Gimelbrant, Erika Giste, Jacob Greene, Mark Groudine, Jessica M. Halow, R. Scott Hansen, Matt E. Hartman, Eric Haugen, Alicia Hill-Force, Richard Humbert, John Hutchinson, Sean Ibarrientos, Mineo Iwata, Sam John, Audra Johnson, Ericka M. Johnson, Rajinder K. Kaul, Marlowe Keller, Cailin Kelly, Tanya Kutyavin, Kristen Lee, Joel Lenox, Matthew T. Maurano, Asia Mendelevich, Wouter Meuleman, Alexander Muratov, Leslie Myatt, Jemma Nelson, Shane Neph, Fidencio Neri, Etaane Neumann, Eric D. Nguyen, Alessandra Odonne Sullivan, Ericka Otterman, Thalia Papayannopoulou, Sadie Patraw, Jocelynn Pearl, Dmitry Penzar, Tamara Pestina, Nikoleta Psatha, Konstantin Queitsch, Alex P. Reynolds, Eric Rynes, Peter J. Sabo, Minerva E. Sanchez, Richard S. Sandstrom, Anthony Shafer, Nurislam Shaikhutdinov, Kyle Siebenthall, John A. Stamatoyannopoulos, George Stamatoyannopoulos, Sandra Stehling-Sun, Andrew B. Stergachis, Shamil R. Sunyaev, Athanasios Teodosiadis, Cameron Trader, Piper M. Treuting, Ben Tudor, Ben Van Biber, Jeff Vierstra, Shinny Vong, Olivia Waltner, Hao Wang, Amanda Watts, Robert Welikson, Matthew S. Wilken, Janghee Woo, Yongqi Yan

##### Baylor College of Medicine

Suhas S. P. Rao^1^, Ragini Mahajan^1^, Dimos Gkountaroulis^1^, Muhammad S. Shamim^1^, Neva C. Durand^2^, Huiya Gu^2^, Per A. Adastra^3^, Galina V. Aglyamova^3^, Ivan D. Bochkov^3^, Saul Godinez Pulido^3^, Shengqi Hang^3^, Shaiza Pasha^3^, Daphne Qin^3^, Mahdi Sadr^3^, Neev Shaw^3^, Mayu Shibata^3^, David Weisz^3^, Alyssa Blackburn^4^, Ishawnia Christopher^4^, Alexander Ellis^4^, Elizabeth L. Hilgert^4^, Su-Chen Huang^4^, Brian Q. Huynh^4^, Ruqayya Khan^4^, Kyle T. Knightly^4^, John A. Krause^4^, Natalie M. Linde^4^, Namita Mitra^4^, Moshe Olshansky^4^, M. Jordan Rowley^4^, Elena Stamenova^4^, Jeffrey Zhong^4^, Daniel B. Aguila Sifuentes^5^, Ohida Binte Amin^5^, Iman S. Asfaw^5^, Rohan Banerjee^5^, Sanjit S. Batra^5^, Xi Chen^5^, Zane L. Colaric^5^, Caitlyn N. Czinder^5^, Allen Dong^5^, Yossi Eliaz^5^, Alexander Elkin^5^, Renzo Espinoza^5^, Cynthia P. Estrada^5^, Thera Fu^5^, Anish Ganti^5^, Hannah L. Harris^5^, Marie Hoeger^5^, Yiwei Jiang^5^, Veda Kadakia^5^, Achyuth Kalluchi^5^, Jongwon D. Kim^5^, Valeriya Korchina^5^, Justin Lee^5^, Michael D. Lee^5^, Li Li^5^, Zoey Ling^5^, Audrey F. Lu^5^, Christopher G. Lui^5^, Michael A. Mancini^5^, Ragib Mostofa^5^, Nathaniel T. Musial^5^, Michael Q. Ngo^5^, Dat T. Nguyen^5^, Guy Nir^5^, Sarah K. Nyquist^5^, Arina D. Omer^5^, Melanie Pham^5^, Amanda N. Potts^5^, Asad Raheem^5^, Mohammad N. Rahman^5^, Sakib Reza^5^, Timothy E. Reznicek^5^, Sheikh Russell^5^, Venny Santosa^5^, Jinyuan Sun^5^, Anat G. Vivante^5^, Evan Vu^5^, Manrong Wu^5^, Weijie Yao^5^, Omer Barad^5^, David Bogumil^5^, Nika Iremadze^5^, Ariel Jamovich^5^, Sarah Pollack^5^, Zohar Shipony^5^, Masato T. Kanemaki, Aviva Presser Aiden, Olga Dudchenko, Erez Lieberman Aiden

^1^Equal contributions
^2^Equal contributions
^3^Equal contributions
^4^Equal contributions
^5^Equal contributions

##### Broad Institute of Harvard and MIT

Nina P. Farrell, Robbyn Issner, Cassandra M. White, Lauren Anderson, Siddarth Wekhande, Eugenio Mattei, Shivani Chinnappan, Michael J. Bennett, Samridhi Banskota, Guillermo Barreto Corona, Alexandra M. Ham, Lauren D. Walter, Joseph J. Raymond, Nicholas H. Nelson, Vivian Hecht, Kevin Dong, Ryuji Morizane, Charles B. Epstein, Noam Shoresh, Elizabeth Gaskell, Chad Nusbaum, Bradley E. Bernstein

##### California Institute of Technology and University of California, Irvine

Brian A. Williams, Fairlie Reese, Elisabeth Rebboah, Narges Rezaie, Muhammed H. Çelik, Heidi Liang, Jasmine S. Sakr, Gabriela Balderrama-Gutierrez, Elnaz Abdollahzadeh, Dana Wyman, Sorena Rahmanian, Rabi Murad, Kate Williams, Klebea C. Sohn, Magdalena Gantuz, Cassandra J. McGill, Diane Trout, Sean Upchurch, Henry Amrhein, Ken McCue, Igor Antoshechkin, Vijaya Kumar, Libera Berghella, Rahma Elsiesy, John Allman, Anthony Linares, Claire K. Zhang, Linjing (Lynn) Fang, Chi (Jina) H. Yun, Long Cai, Laura G. Reinholdt, David A. Bennett, Barbara J. Wold, Ali Mortazavi

##### HudsonAlpha Institute for Biotechnology and University of Alabama in Huntsville

E. Christopher Partridge, Mark Mackiewicz, Daniel Savic, Belle A. Moyers, Kim M. Newberry, Scott Newberry, Amy S. Nesmith, Sarah K. Meadows, Megan M. McEown, Laurel A. Brandsmeier, Jacob Loupe, Timley A. Watkins, Michael J. Betti, Eric M. Mendenhall, Richard M. Myers

##### Stanford University

Yiing Lin, Shin Lin, Annika K. Weimer, Jessika Adrian, Minyi Shi, Emma Monte, Lixia Jiang, Xinqiong Yang, Shannon M. White, Tao Wang, Alec Victorsen, Jie Zhai, Meng Wang, Cory W. Holgren, Jay X. J. Luo, Konor N. von Kraut, Mary Kasparian, Sai Zhang, Johnathan Cooper-Knock, Kevin White, Barbara E. Stranger, Howard Y. Chang, William J. Greenleaf, Michael P. Snyder

##### The Jackson Laboratory for Genomic Medicine

Eva Gega, D’Andre Conaway, Eric T. Loucks, Kun Zhu, Ping Wang, Byoungkoo Lee, Daniel Capurso, Xiaotao Wang, Lina Kozhaya, Sheng Li, Chia-Lin Wei, Derya Unutmaz^1^, Yijun Ruan^1^, Charles Lee^1^

^1^Equal contributions

##### University of Michigan

Ishwarya Venkata Narayanan, Ariel McShane, Karan Bedi, Brian Magnuson, Michelle T. Paulsen, Mario Ashaka, Nina T. Yang, Hailey Blinkiewicz, Mats Ljungman

#### Functional Characterization Centers

##### Broad Institute

Steven K. Reilly, Sager J. Gosai, James R. Xue, Jacob C. Ulirsch, Layla Siraj, Hilary K. Finucane, Alan Gutierrez, Ava Mackay-Smith, Gina M. Butler, Adrianne D. Gladden, Dustin Griesemer, Elizabeth A. Brown, Masahiro Kanai, Pardis C. Sabeti

##### Cornell University

Sagar R. Shah, You Chen, Alden King-Yung Leung, Mauricio I. Paramo, Abdullah Ozer, Zhou Zhou, Li Yao, Junke Zhang, Nathaniel Tippens, Jin Liang, Yiyang Jin, John T. Lis, Haiyuan Yu

##### Duke University

Keith Siklenka, Kuei-Yueh Ko, Katherine Dura, Joshua D. Wheaton, Morgan E. Parker, Stephanie A. Snyder, Naren U. Mehta, Alexandra M. Miggelbrink, Alexias Safi, Lexi Bounds, Tyler Klann, Maria ter Weele, Alex Barrera, Kari Strouse, Graham Johnson, Charles A. Gersbach, Gregory E. Crawford, Maria Ciofani, Timothy E. Reddy

##### Lawrence Berkeley National Laboratory

Diane E. Dickel, Len A. Pennacchio, Axel Visel

##### Stanford University

David Yao, Josh Tycko, Zohar Shipony, Georgi K. Marinov, Eyal Metzl-Raz, Winston R. Becker, Betty B. Liu, Kaitlyn Spees, Nimit Jain, Hua Bai, Arwa Kathiria, Aradhana Aradhana, Benjamin R. Doughty, Julia M. Schaepe, Derek C. Chen, Soon il Higashino, Samuel H. Kim, Sarah E. Pierce, Jeffrey M. Granja, Amy Chen, Benjamin E. Parks, Lacramioara Bintu, Michael C. Bassik, William J. Greenleaf

##### Tempus AI, Inc

Sophia C. Gaynor-Gillett, Lijun Cheng, Manman Shi, Yasuhiro Kyono, Martha Wall, Megan Spector, Mary Flaherty, Donghoon Lee, Jing Zhang, Jason Liu, Zhanlin Chen, Çağatay Dursun, Mark Gerstein, Kevin P. White, Jennifer R. Moran

##### The Jackson Laboratory

Hannah B. Dewey, Rodrigo I. Castro, Susan Kales, Natalia Fuentes, Niketa Nerurkar, Daniel Berenzy, John C. Butts, Kousuke Mouri, Ryan Tewhey

##### University of California, San Francisco and University of California, San Diego

Xingjie Ren, Xiaoyu Yang, Lazaros Lataniotis, Bingkun Li, Maya Asami Takagi, Jerry Lee, Lenka Maliskova, Tsz Wai Tam, Charles Cai, Ian R. Jones, Vivek J. J. Narayan, Cooper Beaman, Poshen Chen, Guoqiang Li, Kai Zhang, Bin Li, Emma Rooholfada, Mengchi Wang, Lina Zheng, Wei Wang, Bing Ren, Yin Shen

##### University of California, San Francisco and University of Washington

Vikram Agarwal, Chengyu Deng, Ryder Easterlin, Molly Gasperini, Ilias Georgakopoulos-Soares, M. Grace Gordon, Fumitaka Inoue, Aidan Keith, Jason C. Klein, Anat Kreimer, Dianne Laboy Cintron, Weiyu Li, Beth K. Martin, Max Schubach, Ajuni Sohota, Chenling Xiong, Jingjing Zhao, Martin Kircher, Jay Shendure, Nadav Ahituv

#### Computational Analysis Centers

##### Harvard University Brigham and Women’s Hospital

Kushal K. Dey, Tiffany Amariuta, Alkes Price, Soumya Raychaudhuri

##### Johns Hopkins University

Jin Woo Oh^1^, Milad Razavi-Mohseni^1^, Dustin Shigaki^1^, Wang Xi^1^, Michael A. Beer

^1^Equal contributions. Authors are listed alphabetically by last name.

##### Memorial Sloan Kettering Cancer Center

Alireza Karbalayghareh, Sneha Mitra, Rui Yang, Vianne Gao, William S. Noble, Christina S. Leslie

##### Stanford University

Vivekanandan Ramalingam^1^, Chang M. Yun^1^, Vivian Hecht^1^, Aman Patel^1^, Anusri Pampari^1^, Ziwei Chen^1^, Johannes Linder^1^,Soumya Kundu^1^, Ivy Evergreen^1^, Austin Wang^1^, Daniel S. Kim^1^, Eran Kotler^1^, Kristy S. Mualim^1^, Georgi K. Marinov^2^, Kelly Cochran^2^, Abhimanyu Banerjee^2^, Surag Nair^2^, Salil S. Deshpande^2^, Zahoor Zafrulla^2^, Riya Sinha^2^, Alex M. Tseng^3^, Amr Alexandari^3^, Mahfuza Sharmin^3^, Avanti Shrikumar,^3^ Jacob M. Schreiber^3^, Caleb Lareau^3^, Andrew R. Marderstein^3^, Marianne K. DeGorter^3^, Page C. Goddard^3^, Paul Flicek^4^, Stephen B. Montgomery^4^, Jonathan K. Pritchard^4^, Anshul Kundaje

^1^Equal contributions
^2^Equal contributions
^3^Equal contributions
^4^Equal contributions

##### University of California, Los Angeles

Ting Fu, Elaine Huang, Giovanni Quinones-Valdez, Kofi Amoah, Ling Zhang, Zhiheng Liu, Jonatan Hervoso, Xinshu Xiao

##### Washington University and Cornell University

Xiaoyu Zhuo^1^, Jason D. Chobirko^1^, Alan Y. Du^1^, Wanqing Shao, Kara Quaid, Mayank N. K. Choudhary, Jiawei Shen, Daofeng Li, Barak A. Cohen^2^, Cedric Feschotte^2^, Ting Wang^2^

^1^Equal contributions
^2^Equal contributions

#### Data Coordination Center

Benjamin C. Hitz, Idan Gabdank, Cricket A. Sloan, Philip Adenekan, Pedro R. Assis, Jessica Au, Ulugbek K.Baymuradov, Carrie Davis, Keenan Graham, Jason Hilton, Otto Jolanki, Jennifer Jou, Meenakshi S. Kagda, Bonita R. Lam, Jin-Wook Lee, Khine Lin, Casey Litton, Yun Hai Luo, Stuart R. Miyasato, Zachary Myers, Katherina Onate, Matt Simison, Emma Spragins, J. Seth Strattan, Paul Sud, Forrest Y. Tanaka, Chris Thomas, Ingrid A. Youngworth, J. Michael Cherry

#### Data Analysis Center

Gregory R. Andrews^1^, Beatrice Borsari^1^, Charles E. Breeze^1^, Christopher J. F. Cameron^1^, Shaimae I. Elhajjajy^1^, Kaili Fan^1^, Jonathan Fisher^1^,Yu Fu^1^, Timur Galeev^1^, Mingshi Gao^1^, Jiahao Gao^1^, Silvia González-López^1^, Gamze Gürsoy^1^, Natalie L. Haas^1^, Yunzhe Jiang^1^, Mansi Khandpekar^1^, Daniel S. Kim^1^, Matthew C. Lacadie^1^, Donghoon Lee^1^, Tianxiao Li^1^, Jason Liu^1^, Susanna Liu^1^, Georgi K. Marinov^1^, Jair Meza^1^, Jill E. Moore^1^, Fabio Navarro^1^, Vasilis Ntasis^1^, Ramil Nurtdinov^1^, Sílvia Pérez-Lluch^1^, Nishigandha Phalke^1^, Huong Phan^1^, Henry E. Pratt^1^, Thomas M. Reimonn^1^, Ferran Reverter^1^, Joel Rozowsky^1^, Marina Ruiz-Romero^1^, Jacob M. Schreiber^1^, Nicole A. Shedd^1^, Yosuke Tanigawa^1^, Xiaohe Tian^1^, Alex M. Tseng^1^, Austin Wang^1^, Jun Wang^1^, Eve S. Wattenberg^1^, Valentin Wucher^1^, Kun Xiong^1^, Jinrui Xu^1^, Yucheng T. Yang^1^, Galip Gürkan Yardımcı^1^,Xuezhu Yu^1^, Suchen Zheng^1^, Roderic Guigó^2^, Manolis Kellis^2^, Anshul Kundaje^2^, William S. Noble^2^, Mark Gerstein^3^, Zhiping Weng^3^

^1^Equal contributions. Authors are listed alphabetically by last name.
^2^Equal contributions. Authors are listed alphabetically by last name.
^3^Equal contributions. Authors are listed alphabetically by last name.

#### Affiliate Members

##### Brown University

Nitya Thakkar, Ritambhara Singh

##### Carnegie Mellon University

Kyle Xiong, Yang Zhang, Jian Ma

##### Center for Theoretical Biological Physics, Rice University, BioScience Research Collaborative

Vinícius G. Contessoto, Antonio B. Oliveira Junior, Esteban Dodero-Rojas, Matheus F. Mello, Sumitabha Brahmachari, Ryan R. Cheng, Michele Di Pierro, Peter G. Wolynes, José N. Onuchic

##### Cold Spring Harbor Laboratory

Alexander Dobin

##### Emory University School of Medicine

Michael H. Nichols, Victor G. Corces

##### Icahn School of Medicine at Mount Sinai

Guo-Cheng Yuan

##### Johns Hopkins Bloomberg School of Public Health

Weixiang Fang, Chaoran Chen, Boyang Zhang, Yi Wang, Ruzhang Zhao, Wenpin Hou, Zhicheng Ji, Weiqiang Zhou, Hongkai Ji

##### Nanyang Technological University and Cancer Science Institute

Ying Zhang, Kaijing Chen, Melissa J. Fullwood

##### Northwestern University

Yu Luan, Ye Hou, Huijue Lyu, Juan Wang, Qixuan Wang, Josiah Hiu-yuen Wong, Yihao Fu, Xintong Chen, Feng Yue

##### Simon Fraser University

Marjan Farahbod, Aboud Diab, Mehdi Foroozandeh, Ishan Goel, Habib Daneshpajouh, Maxwell W. Libbrecht

##### Sloan Kettering Cancer Center

Tabassum Fabiha, Berk Turhan, Thahmina A. Ali, Kushal K. Dey

##### Stanford University

Andreas R. Gschwind, Kristy S. Mualim, Maya U. Sheth, Evelyn Jagoda, Philine Guckelberger, Benjamin R. Doughty, Dulguun Amgalan, Chad J. Munger, Judhajeet Ray, Glen Munson, Vasundhara Singh, Helen Kang, Hank Jones, X. Rosa Ma, Joseph Nasser, Charles P. Fulco, Drew T. Bergman, Tri C. Nguyen, Jesse M. Engreitz

##### University of California, Berkeley

Tal Ashuach, Alyssa Morrow, Nir Yosef

##### University of California, San Diego

Gene W. Yeo

##### University of Connecticut Health Center

Sara Olson, Xintao Wei, Lijun Zhan, Brenton R. Graveley

##### National Human Genome Research Institute^†^

Sarah Anstice, Afia Asare, Stephanie Calluori, Elise A. Feingold, Daniel A. Gilchrist, Carolyn M. Hutter, Stephanie A. Morris, Michael J. Pazin, Ella K. Samer

(National Human Genome Research Institute contributions were limited to coordination and scientific management of the ENCODE Consortium.)

^†^Authors are listed alphabetically by last name

## Acknowledgements

This work was supported by National Human Genome Research Institute ENCODE Program grants to Bradley E. Bernstein and Chad Nusbaum (UM1 HG009390); Erez Lieberman Aiden (UM1 HG009375); Mats Ljungman (UM1 HG009382); Richard M. Myers and Eric Mendenhall (UM1 HG009411); Yijun Ruan and Charles Lee (UM1 HG009409); Michael P. Snyder (UM1 HG009442); John A. Stamatoyannopoulos (UM1 HG009444); Barbara J. Wold and Ali Mortazavi (UM1 HG009443); Nadav Ahituv and Jay Shendure (UM1 HG009408); William J. Greenleaf and Michael C. Bassik (UM1 HG009436); John T. Lis and Haiyuan Yu (UM1 HG009393); Len A. Pennacchio and Axel Visel (UM1 HG009421); Timothy E. Reddy, Maria Ciofani, Gregory E. Crawford and Charles A. Gersbach (UM1 HG009428); Pardis C. Sabeti (UM1 HG009435); Yin Shen and Bing Ren (UM1 HG009402); Kevin White (UM1 HG009426); Michael A. Beer (U01 HG009380); Christina S. Leslie (U01 HG009395); Alkes Price and Soumya Raychaudhuri (U01 HG009379); Jonathan K. Pritchard (U01 HG009431); Ting Wang, Barak A. Cohen and Cedric Feschotte (U01 HG009391); Xinshu Xiao (U01 HG009417); J. Michael Cherry (U24 HG009397); and Zhiping Weng and Mark Gerstein (U24 HG009446).

Additional support was provided by the National Human Genome Research Institute (R01 HG009518 and R01 HG010889 awarded to Hongkai Ji, U24 HG009889 awarded to Brenton R. Graveley and Gene W. Yeo, R01 HG014008 and R00 HG012293 awarded to Kushal K. Dey); by the National Cancer Institute (U01 CA271830 to Sheng Li); by the National Institute of General Medical Sciences (U24 HG009889, R35 GM133562 and R35 GM162228 to Sheng Li); by the National Institute on Aging (R56 AG071766 to Sheng Li); by the National Research Foundation, Singapore under its AI Singapore Programme (AISG Award No: AISG3-GV-2023-014 awarded to Melissa J. Fullwood) and under its NMRC OF-YIRG program (Project No. MOH-001988-00 awarded to Ying Zhang); by the Ministry of Education, Singapore under its Academic Research Fund Tier 1 (RG38/23 awarded to Melissa J. Fullwood); and by the Blood Cancer United (Career Development Award 1398-25 to Sheng Li).

Research conducted at the E.O. Lawrence Berkeley National Laboratory was performed under U.S. Department of Energy Contract DE-AC02-05CH11231 with the University of California.

The Religious Orders Study and Rush Memory and Aging Project (ROSMAP) is supported by P30 AG10161, P30 AG72975, R01 AG17917, R01 AG015819, U01 AG072572, and U01 AG046152.

We gratefully acknowledge Tom Gingeras and Michael Schatz for their leadership roles in the EN-TEx project as members of the ENCODE 3 consortium.

## References

1 Lander, E. S. et al. Initial sequencing and analysis of the human genome. Nature 409, 860–921 (2001). 10.1038/35057062

2 Venter, J. C. et al. The sequence of the human genome. Science 291, 1304–1351 (2001). 10.1126/science.1058040

3 Nurk, S. et al. The complete sequence of a human genome. Science 376, 44–53 (2022). 10.1126/science.abj6987

4 Birney, E. et al. Identification and analysis of functional elements in 1% of the human genome by the ENCODE pilot project. Nature 447, 799–816 (2007). 10.1038/nature05874

5 An integrated encyclopedia of DNA elements in the human genome. Nature 489, 57–74 (2012). 10.1038/nature11247

6 Stamatoyannopoulos, J. A. et al. An encyclopedia of mouse DNA elements (Mouse ENCODE). Genome Biol 13, 418 (2012). 10.1186/gb-2012-13-8-418

7 Gerstein, M. B. et al. Integrative analysis of the Caenorhabditis elegans genome by the modENCODE project. Science 330, 1775–1787 (2010). 10.1126/science.1196914

8 Roy, S. et al. Identification of functional elements and regulatory circuits by Drosophila modENCODE. Science 330, 1787–1797 (2010). 10.1126/science.1198374

9 Moore, J. E. et al. Expanded encyclopaedias of DNA elements in the human and mouse genomes. Nature 583, 699–710 (2020). 10.1038/s41586-020-2493-4

10 Rozowsky, J. et al. The EN-TEx resource of multi-tissue personal epigenomes & variant-impact models. Cell 186, 1493–1511.e1440 (2023). 10.1016/j.cell.2023.02.018

11 The GTEx Consortium atlas of genetic regulatory effects across human tissues. Science 369, 1318–1330 (2020). 10.1126/science.aaz1776

12 Companion manuscript ENC4P103 (“The Human Fetal Regulome And Transcriptome”) is in preparation. Please refer to the included abstract.

13 Companion manuscript ENC4P11 (“A Comprehensive Catalog Of Accessible DNA Elements Encoded By The Human Genome”) is in preparation. Please refer to included abstract list.

14 The ENCODE (ENCyclopedia Of DNA Elements) Project. Science 306, 636–640 (2004). 10.1126/science.1105136

15 van Galen, P. et al. A Multiplexed System for Quantitative Comparisons of Chromatin Landscapes. Mol Cell 61, 170–180 (2016). 10.1016/j.molcel.2015.11.003

16 Ernst, J. & Kellis, M. ChromHMM: automating chromatin-state discovery and characterization. Nat Methods 9, 215–216 (2012). 10.1038/nmeth.1906

17 Chan, R. C. W. et al. Segway 2.0: Gaussian mixture models and minibatch training. Bioinformatics 34, 669–671 (2018). 10.1093/bioinformatics/btx603

18 Farahbod, M. et al. Integrative chromatin state annotation of 234 human ENCODE4 cell types using Segway. Genome Res (2025). 10.1101/gr.280633.125

19 Huang, D., Petrykowska, H. M., Miller, B. F., Elnitski, L. & Ovcharenko, I. Identification of human silencers by correlating cross-tissue epigenetic profiles and gene expression. Genome Res 29, 657–667 (2019). 10.1101/gr.247007.118

20 Doni Jayavelu, N., Jajodia, A., Mishra, A. & Hawkins, R. D. Candidate silencer elements for the human and mouse genomes. Nat Commun 11, 1061 (2020). 10.1038/s41467-020-14853-5

21 Pang, B. & Snyder, M. P. Systematic identification of silencers in human cells. Nat Genet 52, 254–263 (2020). 10.1038/s41588-020-0578-5

22 Cai, Y. et al. H3K27me3-rich genomic regions can function as silencers to repress gene expression via chromatin interactions. Nat Commun 12, 719 (2021). 10.1038/s41467-021-20940-y

23 Ngan, C. Y. et al. Chromatin interaction analyses elucidate the roles of PRC2-bound silencers in mouse development. Nat Genet 52, 264–272 (2020). 10.1038/s41588-020-0581-x

24 Moore, J. E. et al. An expanded registry of candidate cis-regulatory elements. Nature (2026). 10.1038/s41586-025-09909-9

25 Zhang, Y. et al. Super-silencer perturbation by EZH2 and REST inhibition leads to large loss of chromatin interactions and reduction in cancer growth. Nat Struct Mol Biol 32, 137–149 (2025). 10.1038/s41594-024-01391-7

26 Partridge, E. C. et al. Occupancy maps of 208 chromatin-associated proteins in one human cell type. Nature 583, 720–728 (2020). 10.1038/s41586-020-2023-4

27 Wang, J. et al. Sequence features and chromatin structure around the genomic regions bound by 119 human transcription factors. Genome Res 22, 1798–1812 (2012). 10.1101/gr.139105.112

28 Lambert, S. A. et al. The Human Transcription Factors. Cell 172, 650–665 (2018). 10.1016/j.cell.2018.01.029

29 Savic, D. et al. CETCh-seq: CRISPR epitope tagging ChIP-seq of DNA-binding proteins. Genome Res 25, 1581–1589 (2015). 10.1101/gr.193540.115

30 Moyers, B. A. et al. Characterization of human transcription factor function and patterns of gene regulation in HepG2 cells. Genome Res 33, 1879–1892 (2023). 10.1101/gr.278205.123

31 Seo, J. et al. AP-1 subunits converge promiscuously at enhancers to potentiate transcription. Genome Res 31, 538–550 (2021). 10.1101/gr.267898.120

32 White, S. M. et al. Comparative atlas of genome-wide chromatin-associated protein co-occupancy. bioRxiv, 2024.2012.2017.628199 (2024). 10.1101/2024.12.17.628199

33 Core, L. J. et al. Analysis of nascent RNA identifies a unified architecture of initiation regions at mammalian promoters and enhancers. Nat Genet 46, 1311–1320 (2014). 10.1038/ng.3142

34 Kwak, H., Fuda, N. J., Core, L. J. & Lis, J. T. Precise maps of RNA polymerase reveal how promoters direct initiation and pausing. Science 339, 950–953 (2013). 10.1126/science.1229386

35 Magnuson, B. et al. Identifying transcription start sites and active enhancer elements using BruUV-seq. Sci Rep 5, 17978 (2015). 10.1038/srep17978

36 Shah, S. R. et al. The regulatory landscape of nascent transcription in human health and disease. bioRxiv (2025). 10.1101/2025.09.24.676871

37 McShane, A. et al. Characterizing nascent transcription patterns of PROMPTs, eRNAs, and readthrough transcripts in the ENCODE4 deeply profiled cell lines. bioRxiv (2024). 10.1101/2024.04.09.588612

38 Cochran, K. et al. Dissecting the cis-regulatory syntax of transcription initiation with deep learning. bioRxiv (2024). 10.1101/2024.05.28.596138

39 Zhang, J. et al. Comprehensive evaluation of diverse massively parallel reporter assays to functionally characterize human enhancers genome-wide. Genome Biol 26, 378 (2025). 10.1186/s13059-025-03828-8

40 Paramo, M. I. et al. Dual promoter-enhancer activities reflect a unified regulatory logic. Nat Commun 17 (2026). 10.1038/s41467-026-68780-y

41 Dudnyk, K., Cai, D., Shi, C., Xu, J. & Zhou, J. Sequence basis of transcription initiation in the human genome. Science 384, eadj0116 (2024). 10.1126/science.adj0116

42 He, A. Y. & Danko, C. G. Dissection of core promoter syntax through single nucleotide resolution modeling of transcription initiation. bioRxiv (2024). 10.1101/2024.03.13.583868

43 Chen, Y. et al. Directionality of transcriptional regulatory elements. bioRxiv, 2024.2012.2001.621925 (2024). 10.1101/2024.12.01.621925

44 Companion manuscript ENC4P06 (“Functional Characterization Of Regulatory Elements In The Human Genome”) is in preparation. Please refer to included abstract list.

45 Yao, D. et al. Multicenter integrated analysis of noncoding CRISPRi screens. Nat Methods 21, 723–734 (2024). 10.1038/s41592-024-02216-7

46 Yang, X. et al. Functional characterization of gene regulatory elements and neuropsychiatric disease-associated risk loci in iPSCs and iPSC-derived neurons. bioRxiv, 2023.2008.2030.555359 (2023). 10.1101/2023.08.30.555359

47 Ren, X. et al. CRISPR tiling deletion screens reveal functional enhancers and allelic compensation effects (ACE) on SIN3A transcription. Nat Commun 17 (2026). 10.1038/s41467-026-70933-y

48 Siklenka, K. et al. Enhancer hubs govern chromatin topology and Th17 cell identity. bioRxiv (2026). 10.64898/2026.04.02.715458

49 Bergman, D. T. et al. Compatibility rules of human enhancer and promoter sequences. Nature 607, 176–184 (2022). 10.1038/s41586-022-04877-w

50 Pampari, A. et al. ChromBPNet: bias factorized, base-resolution deep learning models of chromatin accessibility reveal cis-regulatory sequence syntax, transcription factor footprints and regulatory variants. bioRxiv (2025). 10.1101/2024.12.25.630221

51 Avsec, Ž., et al. Base-resolution models of transcription-factor binding reveal soft motif syntax. Nat Genet 53, 354–366 (2021). 10.1038/s41588-021-00782-6

52 Companion manuscript ENC4P03 (“A Unified Lexicon Of Predictive DNA Sequence Motifs From ENCODE Transcription Factor Binding And Chromatin Accessibility Assays”) is in preparation. Please refer to the included abstract.

53 Companion manuscript ENC4P38 (“Deep Learning The Alternate Recognition Modes Of Human C2H2 Zinc Finger Transcription Factors”) is in preparation. Please refer to included abstract list.

54 Agarwal, V. et al. Massively parallel characterization of transcriptional regulatory elements. Nature 639, 411–420 (2025). 10.1038/s41586-024-08430-9

55 Companion manuscript ENC4P93 (“ReporterNet Predicts Regulatory Activity For Sequences In Diverse Reporter Assays And Enables Systematic Comparative Analysis Of High-throughput Reporter Assays”) is in preparation. Please refer to included abstract list.

56 Shrikumar, A., Greenside, P. & Kundaje, A. in International conference on machine learning. 3145–3153 (PMlR).

57 Shrikumar, A. et al. Technical note on transcription factor motif discovery from importance scores (TF-MoDISco) version 0.5. 6.5. arXiv preprint arXiv:1811.00416 (2018).

58 Companion manuscript ENC4P39 (“Decoding Predictive Motif Lexicons And Syntax From Deep Learning Models Of Transcription Factor Binding Profiles”) is in preparation. Please refer to included abstract list.

59 Zhang, F. & Lupski, J. R. Non-coding genetic variants in human disease. Hum Mol Genet 24, R102–110 (2015). 10.1093/hmg/ddv259

60 Lappalainen, T. & MacArthur, D. G. From variant to function in human disease genetics. Science 373, 1464–1468 (2021). 10.1126/science.abi8207

61 Finucane, H. K. et al. Heritability enrichment of specifically expressed genes identifies disease-relevant tissues and cell types. Nat Genet 50, 621–629 (2018). 10.1038/s41588-018-0081-4

62 Sollis, E. et al. The NHGRI-EBI GWAS Catalog: knowledgebase and deposition resource. Nucleic Acids Res 51, D977–d985 (2023). 10.1093/nar/gkac1010

63 Siraj, L. et al. Functional dissection of complex and molecular trait variants at single nucleotide resolution. bioRxiv (2024). 10.1101/2024.05.05.592437

64 Fabiha, T. et al. A consensus variant-to-function score to functionally prioritize variants for disease. bioRxiv, 2024.2011.2007.622307 (2024). 10.1101/2024.11.07.622307

65 Gaziano, J. M. et al. Million Veteran Program: A mega-biobank to study genetic influences on health and disease. J Clin Epidemiol 70, 214–223 (2016). 10.1016/j.jclinepi.2015.09.016

66 Verma, A. et al. Diversity and scale: Genetic architecture of 2068 traits in the VA Million Veteran Program. Science 385, eadj1182 (2024). 10.1126/science.adj1182

67 Bycroft, C. et al. The UK Biobank resource with deep phenotyping and genomic data. Nature 562, 203–209 (2018). 10.1038/s41586-018-0579-z

68 Tanigawa, Y. & Kellis, M. Power of inclusion: Enhancing polygenic prediction with admixed individuals. Am J Hum Genet 110, 1888–1902 (2023). 10.1016/j.ajhg.2023.09.013

69 Tian, X. et al. PRISM: ancestry-aware integration of tissue-specific genomic annotations enhances the transferability of polygenic scores. bioRxiv (2025). 10.1101/2025.11.13.688144

70 Lagarde, J. et al. High-throughput annotation of full-length long noncoding RNAs with capture long-read sequencing. Nat Genet 49, 1731–1740 (2017). 10.1038/ng.3988

71 Perteghella, T. et al. The GENCODE CLS project: massively expanding the lncRNA catalog through capture long-read RNA sequencing. bioRxiv, 2024.2010.2029.620654 (2026). 10.1101/2024.10.29.620654

72 Reese, F. et al. The ENCODE4 long-read RNA-seq collection reveals distinct classes of transcript structure diversity. bioRxiv (2023). 10.1101/2023.05.15.540865

73 Quinones-Valdez, G., Amoah, K. & Xiao, X. Long-read RNA-seq demarcates cis-and trans-directed alternative RNA splicing. Nat Commun 16, 9603 (2025). 10.1038/s41467-025-64605-6

74 Faucillion, M. L., Johansson, A. M. & Larsson, J. Modulation of RNA stability regulates gene expression in two opposite ways: through buffering of RNA levels upon global perturbations and by supporting adapted differential expression. Nucleic Acids Res 50, 4372–4388 (2022). 10.1093/nar/gkac208

75 Dowdle, M. E. & Lykke-Andersen, J. Cytoplasmic mRNA decay and quality control machineries in eukaryotes. Nat Rev Genet 26, 463–478 (2025). 10.1038/s41576-024-00810-1

76 Huang, E. et al. Genetic variants affecting RNA stability influence complex traits and disease risk. Nat Genet 57, 2578–2588 (2025). 10.1038/s41588-025-02326-8

77 Paulsen, M. T. et al. Coordinated regulation of synthesis and stability of RNA during the acute TNF-induced proinflammatory response. Proc Natl Acad Sci U S A 110, 2240–2245 (2013). 10.1073/pnas.1219192110

78 Paulsen, M. T. et al. Use of Bru-Seq and BruChase-Seq for genome-wide assessment of the synthesis and stability of RNA. Methods 67, 45–54 (2014). 10.1016/j.ymeth.2013.08.015

79 Bedi, K. et al. Isoform and pathway-specific regulation of post-transcriptional RNA processing in human cells. bioRxiv (2024). 10.1101/2024.06.12.598705

80 Fu, T. et al. Massively parallel screen uncovers many rare 3’ UTR variants regulating mRNA abundance of cancer driver genes. Nat Commun 15, 3335 (2024). 10.1038/s41467-024-46795-7

81 Van Nostrand, E. L. et al. A large-scale binding and functional map of human RNA-binding proteins. Nature 583, 711–719 (2020). 10.1038/s41586-020-2077-3

82 Gosztyla, M. L. et al. Integrated multi-omics analysis of zinc-finger proteins uncovers roles in RNA regulation. Mol Cell 84, 3826–3842.e3828 (2024). 10.1016/j.molcel.2024.08.010

83 Her, H. L. et al. Comprehensive RNA-binding protein analyses and deep learning uncover genetic constraints and disease associations in protein-RNA interfaces. Cell Syst, 101588 (2026). 10.1016/j.cels.2026.101588

84 Companion manuscript ENC4P10 (“An Atlas Of Nuclear Architecture Resolves Element-to-element Chromatin Loops In Over 60 Human Tissues”) is in preparation. Please refer to included abstract list.

85 Hsieh, T. H. et al. Mapping Nucleosome Resolution Chromosome Folding in Yeast by Micro-C. Cell 162, 108–119 (2015). 10.1016/j.cell.2015.05.048

86 Slobodyanyuk, E., Cattoglio, C. & Hsieh, T. S. Mapping Mammalian 3D Genomes by Micro-C. Methods Mol Biol 2532, 51–71 (2022). 10.1007/978-1-0716-2497-5_4

87 Wang, P. et al. Chromatin topology reorganization and transcription repression by PML-RARα in acute promyeloid leukemia. Genome Biol 21, 110 (2020). 10.1186/s13059-020-02030-2

88 Chai, H. et al. ChIATAC is an efficient strategy for multi-omics mapping of 3D epigenomes from low-cell inputs. Nat Commun 14, 213 (2023). 10.1038/s41467-023-35879-5

89 Wang, P. et al. In situ Chromatin Interaction Analysis Using Paired-End Tag Sequencing. Curr Protoc 1, e174 (2021). 10.1002/cpz1.174

90 Oliveira Junior, A. B., Contessoto, V. G., Mello, M. F. & Onuchic, J. N. A Scalable Computational Approach for Simulating Complexes of Multiple Chromosomes. J Mol Biol 433, 166700 (2021). 10.1016/j.jmb.2020.10.034

91 Di Pierro, M., Cheng, R. R., Lieberman Aiden, E., Wolynes, P. G. & Onuchic, J. N. De novo prediction of human chromosome structures: Epigenetic marking patterns encode genome architecture. Proc Natl Acad Sci U S A 114, 12126–12131 (2017). 10.1073/pnas.1714980114

92 Guckelberger, P. et al. Cohesin-mediated 3D contacts tune enhancer-promoter regulation. bioRxiv (2024). 10.1101/2024.07.12.603288

93 Gschwind, A. R. et al. An encyclopedia of enhancer-gene regulatory interactions in the human genome. bioRxiv (2023). 10.1101/2023.11.09.563812

94 Fang, W. et al. Quantifying Functional Conservation of Human and Mouse Regulatory Elements via FUNCODE. bioRxiv, 2024.2010.2031.620766 (2024). 10.1101/2024.10.31.620766

95 Oh, J. W. & Beer, M. A. Gapped-kmer sequence modeling robustly identifies regulatory vocabularies and distal enhancers conserved between evolutionarily distant mammals. Nat Commun 15, 6464 (2024). 10.1038/s41467-024-50708-z

96 Rebollo, R., Romanish, M. T. & Mager, D. L. Transposable elements: an abundant and natural source of regulatory sequences for host genes. Annu Rev Genet 46, 21–42 (2012). 10.1146/annurev-genet-110711-155621

97 Chuong, E. B., Elde, N. C. & Feschotte, C. Regulatory activities of transposable elements: from conflicts to benefits. Nat Rev Genet 18, 71–86 (2017). 10.1038/nrg.2016.139

98 Bourque, G. et al. Ten things you should know about transposable elements. Genome Biol 19, 199 (2018). 10.1186/s13059-018-1577-z

99 Sundaram, V. & Wysocka, J. Transposable elements as a potent source of diverse cis-regulatory sequences in mammalian genomes. Philos Trans R Soc Lond B Biol Sci 375, 20190347 (2020). 10.1098/rstb.2019.0347

100 Fueyo, R., Judd, J., Feschotte, C. & Wysocka, J. Roles of transposable elements in the regulation of mammalian transcription. Nat Rev Mol Cell Biol 23, 481–497 (2022). 10.1038/s41580-022-00457-y

101 Lawson, H. A., Liang, Y. & Wang, T. Transposable elements in mammalian chromatin organization. Nat Rev Genet 24, 712–723 (2023). 10.1038/s41576-023-00609-6

102 Zhang, X. O., Pratt, H. & Weng, Z. Investigating the Potential Roles of SINEs in the Human Genome. Annu Rev Genomics Hum Genet 22, 199–218 (2021). 10.1146/annurev-genom-111620-100736

103 Zhang, X. O., Gingeras, T. R. & Weng, Z. Genome-wide analysis of polymerase III-transcribed Alu elements suggests cell-type-specific enhancer function. Genome Res 29, 1402–1414 (2019). 10.1101/gr.249789.119

104 Du, A. Y., Chobirko, J. D., Zhuo, X., Feschotte, C. & Wang, T. Regulatory transposable elements in the encyclopedia of DNA elements. Nat Commun 15, 7594 (2024). 10.1038/s41467-024-51921-6

105 Du, A. Y. et al. Functional characterization of enhancer activity during a long terminal repeat’s evolution. Genome Res 32, 1840–1851 (2022). 10.1101/gr.276863.122

106 Sandoval-Velasco, M. et al. Three-dimensional genome architecture persists in a 52,000-year-old woolly mammoth skin sample. Cell 187, 3541–3562.e3551 (2024). 10.1016/j.cell.2024.06.002

107 Che, Y. et al. The evolution of high-order genome architecture revealed from 1,000 species. Cell (2026). 10.1016/j.cell.2026.03.042

108 Kagda, M. S. et al. Data navigation on the ENCODE portal. Nat Commun 16, 9592 (2025). 10.1038/s41467-025-64343-9

109 Casper, J. et al. The UCSC Genome Browser database: 2026 update. Nucleic Acids Res 54, D1331–D1335 (2026). 10.1093/nar/gkaf1250

110 Companion manuscript ENC4P88 (“Juicer 2.0 Analyzes And Visualizes Contact Mapping Experiments At Base-pair Resolution”) is in preparation. Please refer to included abstract list.

111 Companion manuscript ENC4P89 (“Spacewalk Provides A Cloud-based Visualization System For Microscopy Of The Genome”) is in preparation. Please refer to included abstract list.

112 Boyle, A. P. et al. Annotation of functional variation in personal genomes using RegulomeDB. Genome Res 22, 1790–1797 (2012). 10.1101/gr.137323.112

113 Fishilevich, S. et al. GeneHancer: genome-wide integration of enhancers and target genes in GeneCards. Database (Oxford) 2017 (2017). 10.1093/database/bax028

114 Schubach, M., Maass, T., Nazaretyan, L., Röner, S. & Kircher, M. CADD v1.7: using protein language models, regulatory CNNs and other nucleotide-level scores to improve genome-wide variant predictions. Nucleic Acids Res 52, D1143–d1154 (2024). 10.1093/nar/gkad989

115 Wang, K., Li, M. & Hakonarson, H. ANNOVAR: functional annotation of genetic variants from high-throughput sequencing data. Nucleic Acids Res 38, e164 (2010). 10.1093/nar/gkq603

116 Zhang, J. et al. An integrative ENCODE resource for cancer genomics. Nat Commun 11, 3696 (2020). 10.1038/s41467-020-14743-w

117 Xu, M., Yamamoto, T. & Kato, T. In vivo striatal dopamine release by M1 muscarinic receptors is induced by activation of protein kinase C. J Neurochem 54, 1917–1919 (1990). 10.1111/j.1471-4159.1990.tb04891.x

118 Steijger, T. et al. Assessment of transcript reconstruction methods for RNA-seq. Nat Methods 10, 1177–1184 (2013). 10.1038/nmeth.2714

119 Pardo-Palacios, F. J. et al. Systematic assessment of long-read RNA-seq methods for transcript identification and quantification. bioRxiv (2023). 10.1101/2023.07.25.550582

120 Zargari, A. et al. DeepSea is an efficient deep-learning model for single-cell segmentation and tracking in time-lapse microscopy. Cell Rep Methods 3, 100500 (2023). 10.1016/j.crmeth.2023.100500

121 Zhou, J. et al. Deep learning sequence-based ab initio prediction of variant effects on expression and disease risk. Nat Genet 50, 1171–1179 (2018). 10.1038/s41588-018-0160-6

122 Avsec, Ž., et al. Effective gene expression prediction from sequence by integrating long-range interactions. Nat Methods 18, 1196–1203 (2021). 10.1038/s41592-021-01252-x

123 Avsec, Ž., et al. Advancing regulatory variant effect prediction with AlphaGenome. Nature 649, 1206–1218 (2026). 10.1038/s41586-025-10014-0

124 Linder, J., Srivastava, D., Yuan, H., Agarwal, V. & Kelley, D. R. Predicting RNA-seq coverage from DNA sequence as a unifying model of gene regulation. Nat Genet 57, 949–961 (2025). 10.1038/s41588-024-02053-6

125 Mudge, J. M. et al. GENCODE 2025: reference gene annotation for human and mouse. Nucleic Acids Res 53, D966–d975 (2025). 10.1093/nar/gkae1078

126 Rao, S. S. et al. A 3D map of the human genome at kilobase resolution reveals principles of chromatin looping. Cell 159, 1665–1680 (2014). 10.1016/j.cell.2014.11.021

127 Dekker, J. et al. An integrated view of the structure and function of the human 4D nucleome. Nature 649, 759–776 (2026). 10.1038/s41586-025-09890-3

128 Ghiasvand, N. M. et al. Deletion of a remote enhancer near ATOH7 disrupts retinal neurogenesis, causing NCRNA disease. Nat Neurosci 14, 578–586 (2011). 10.1038/nn.2798

