## Supplementary Information for "The Encyclopedia of DNA Elements"

### Supplementary Boxes

#### Supplementary Box 1 | Glossary of ENCODE Terms

**ENCODE:** The Encyclopedia of DNA Elements

**DNA Element:** A DNA sequence in the human genome that either possesses a particular function (a functional element) or has been nominated as functional based on an experimental assay. The two major types of DNA elements ENCODE studied were **genes** and **regulatory elements.**

**Gene**: A DNA sequence that is transcribed into a functional RNA. There are two types of genes: (i) protein-coding genes, where the resultant RNA is translated into a protein by the ribosome; and (ii) non-coding RNA genes, which are not translated into proteins.

**Transcription start site (TSS):** the genomic position where the RNA polymerase II begins making the RNA transcript of a gene. Many genes have multiple TSSs.

**Regulatory Element:** Modular DNA element that contributes to the control of gene expression. Major classes of regulatory elements include **promoters**, **enhancers**, **silencers**, and **insulators**. Regulatory elements primarily impact the transcriptional state or potential of one or more genes lying on the same physical DNA molecule the regulatory element, generally in close linear proximity.

**Promoter:** A DNA element near the TSS of a gene where the basic protein machinery responsible for transcription of that gene assemble.

**Enhancer:** A short (typically < 500 bp) modular DNA element, distinct from the promoter, that increases gene expression. Enhancers are bound by transcription factors and act on target promoters that are often at far away in the linear distance of the genome.

**Silencer:** A DNA element that actively reduces gene expression. Like enhancers, they can act over long distances in the linear genome sequence.

**Insulator:** A DNA element that directly impedes the physical interaction of distal regulatory elements (e.g., enhancers) with a target promoter when interposed between them. Typically, insulators are bound by the protein CTCF.

**DNase hypersensitive site (DHS)**: A DNA element that is hypersensitive to cleavage in the context of native chromatin by the non-specific endonuclease DNase I. DHSs represent nucleosome-free regions and encompass DNA elements that are accessible to other nucleases or to transposases.

**Transcription factor binding site (TFBS)**: A short DNA element (typically < 20bp) that is the recognition site for a sequence-specific DNA-binding protein known as a transcription factor to bind.

**Enhancer RNA (eRNA)**: Small unstable directional RNA transcript produced by the recruitment of RNA polymerase to some enhancers by the occupying transcription factors. Some eRNAs may modulate the function of their cognate element.

**Candidate cis-regulatory element (cCRE)**: An ENCODE annotation anchored on DHSs that categorize a given DNA element into one of seven functional classes based on genomic position, histone modification pattern, transcription factor occupancy pattern, and the presence of nascent RNA transcription.

**Loop anchor:** A DNA element that lies at the base of a long-range physical interaction.

#### Supplementary Box 2 | Overview of ENCODE 4 Sample Collections

ENCODE organized coordinated studies on several sample collections. Major ENCODE collections are described here.

- **Human primary tissue samples:** To detail gene regulatory state broadly across the human body, we measured gene expression, chromatin state, and chromatin interactions across 122 different tissues or cell types derived from 319 human donors. Those assays include the ENTEx effort, detailed below, to measure gene regulatory state across 30 human tissues from four GTEx donors consented for full open data release^1-3^; analyses of human tissues obtained from surgical resections, cadavers; and analyses of >50 different immune cell types and states collected from human donors.
- [Epigenomes from four individuals (EN-TEx)](https://www.encodeproject.org/entex-matrix/?type=Experiment&status=released&internal_tags=ENTEx): The goal of the ENTEx collection is to establish maps of regulatory activity across the human body. The ENTEx effort completed 1567 assays of gene expression and chromatin state across 31 tissues from 4 post-mortem donors. Tissues from the same 4 donors were also studied in the GTEx project, and samples were collected for GTEx and ENTEx contemporaneously and from matched tissue sites.^2,3^ The collection includes assays of gene expression, transcription start site use, bulk and single cell chromatin accessibility, histone modification, and genomic occupancy of CTCF and RNA Pol II.
- **Population scale studies of gene regulatory state in primary tissues**: To investigate variation in gene regulatory state across donors and disease state, ENCODE 4 completed several population-scale studies. The Rush Alzheimer’s disease study analyzed samples from 120 donors with varying levels of cognitive impairment due to Alzheimer’s disease. The study measured chromatin accessibility, histone modifications, gene expression, and long-range chromatin interactions in dorsolateral prefrontal cortex; and measured chromatin accessibility in samples of the head of the caudate nucleus and of the posterior cingulate gyrus. ENCODE also completed single cell analyses of heart ventricles from 71 donors.
- [Epigenomic profiling of human immune cells:](https://www.encodeproject.org/immune-cells/?type=Experiment&replicates.library.biosample.donor.organism.scientific_name=Homo+sapiens&biosample_ontology.cell_slims=hematopoietic+cell&biosample_ontology.classification=primary+cell&control_type!=*&status!=replaced&status!=revoked&status!=archived&biosample_ontology.system_slims=immune+system&biosample_ontology.system_slims=circulatory+system&config=immune) The goal of the immune cell collection is to detail differences in regulatory state between diverse types and states of human immune cells including T cells, B cells, myeloid cells, and natural killer cells. In total, the collection comprises >50 different sorted, differentiated, and stimulated human immune cells from >80 donors; including >7 cell types from donors with multiple sclerosis and matched controls. The assays completed include measurements of bulk and single cell chromatin accessibility, histone modifications, gene expression, and long-range physical interactions.
- [Uniform assay reference](https://www.encodeproject.org/deeply-profiled-uniform-batch-matrix/?type=Experiment&control_type!=*&status=released&replicates.library.biosample.biosample_ontology.term_id=EFO:0002106&replicates.library.biosample.biosample_ontology.term_id=EFO:0001203&replicates.library.biosample.biosample_ontology.term_id=EFO:0006711&replicates.library.biosample.biosample_ontology.term_id=EFO:0002713&replicates.library.biosample.biosample_ontology.term_id=EFO:0002847&replicates.library.biosample.biosample_ontology.term_id=EFO:0002074&replicates.library.biosample.biosample_ontology.term_id=EFO:0001200&replicates.library.biosample.biosample_ontology.term_id=EFO:0009747&replicates.library.biosample.biosample_ontology.term_id=EFO:0002824&replicates.library.biosample.biosample_ontology.term_id=CL:0002327&replicates.library.biosample.biosample_ontology.term_id=CL:0002618&replicates.library.biosample.biosample_ontology.term_id=EFO:0002784&replicates.library.biosample.biosample_ontology.term_id=EFO:0001196&replicates.library.biosample.biosample_ontology.term_id=EFO:0001187&replicates.library.biosample.biosample_ontology.term_id=EFO:0002067&replicates.library.biosample.biosample_ontology.term_id=EFO:0001099&replicates.library.biosample.biosample_ontology.term_id=EFO:0002819&replicates.library.biosample.biosample_ontology.term_id=EFO:0009318&replicates.library.biosample.biosample_ontology.term_id=EFO:0001086&replicates.library.biosample.biosample_ontology.term_id=EFO:0007950&replicates.library.biosample.biosample_ontology.term_id=EFO:0003045&replicates.library.biosample.biosample_ontology.term_id=EFO:0003042&replicates.library.biosample.internal_tags=Deeply%20Profiled): The goal of the uniform assay reference is to enable systematic and integrative comparisons of gene regulatory state across highly consistent and uniformly grown cell lines. To do so, ENCODE 4 centralized growth of cells from 16 cell lines that are commonly used in biomedical research. Cells were then distributed for assays of bulk and single cell chromatin accessibility, gene expression, nascent transcription, RNA degradation, and long-range chromatin interactions.^4,5^
- [Targeted degradation of proteins:](https://www.encodeproject.org/degron-matrix/?type=Experiment&control_type!=*&status=released&internal_tags=Degron) The goal of the targeted degradation collection is to determine the role of eight proteins on the physical structure and regulatory state of the human genome. The collection used an auxin-inducible degron system to rapidly degrade key proteins involved in aspects of transcription or chromatin structure – CTCF, RAD21, BRD4, CDK7, POL2RA, SMARCA5, MED14, and SUPT16H – in HCT116. After degradation of each protein, measurements of chromatin accessibility, histone modifications, gene expression, nascent transcription, and long-range chromatin interactions were completed.^6,7^
- [African functional genomics resource](https://github.com/smontgomlab/AFGR): The goal of the African functional genomics resource is to measure chromatin accessibility and gene expression across the global human population. The collection includes RNA-seq on 599 lymphoblastoid cell lines derived from 1000 Genomes and HapMap3 donors across six African population samples, and ATAC-seq on a 100-sample subset. The resulting resource provides a matched set of transcriptomic and chromatin accessibility profiles, quantitative trait loci, and observed and predicted allelic effects while expanding the global diversity of human cell line data in ENCODE.
- [Mouse postnatal developmental time series](https://www.encodeproject.org/mouse-development-matrix/?type=Experiment&status=released&related_series.@type=OrganismDevelopmentSeries&replicates.library.biosample.organism.scientific_name=Mus+musculus): A major focus of ENCODE has been to measure changes in gene regulation across mouse development. Previous phases of ENCODE have measured gene regulatory state in cell lines, embryos, adults and over prenatal development. ENCODE 4 added measurements across postnatal development from birth to 20 months of age. Together with previous phases of ENCODE, the full lifespan of the mouse is now represented. Assays in ENCODE 4 focused on bulk and single cell gene expression measurements, including via long read RNA-seq, as well as bulk and single nucleus chromatin accessibility measurements.^8^
- [Stem cell differentiation studies:](https://www.encodeproject.org/stem-cell-matrix/?type=Experiment&replicates.library.biosample.donor.accession=ENCDO222AAA&status=released&control_type!=*) The goal of the stem cell differentiation collection is to measure changes in gene regulatory state as embryonic stem cells and induced pluripotent stem cells differentiate into other cell types and into organoids. The differentiated cell types reflect the three major cell lineages: mesoderm, endoderm and ectoderm. The collection focused on measuring chromatin accessibility, histone modifications, and gene expression across 16 differentiated cell types and two organoid models (brain and kidney).
- B cell trans-differentiation studies: The goal of the B cell trans-differentiation collection is to investigate the relationship between chromatin and transcriptional dynamics during a differentiation process from a lineage committed cell type (B-cells to macrophages). The collection focused on measuring RNA states for the whole cell, and the nuclear and cytosolic fractions, ribosome occupancy, protein content, and chromatin states. For more details, see Case Study 6.

### Supplementary Figures

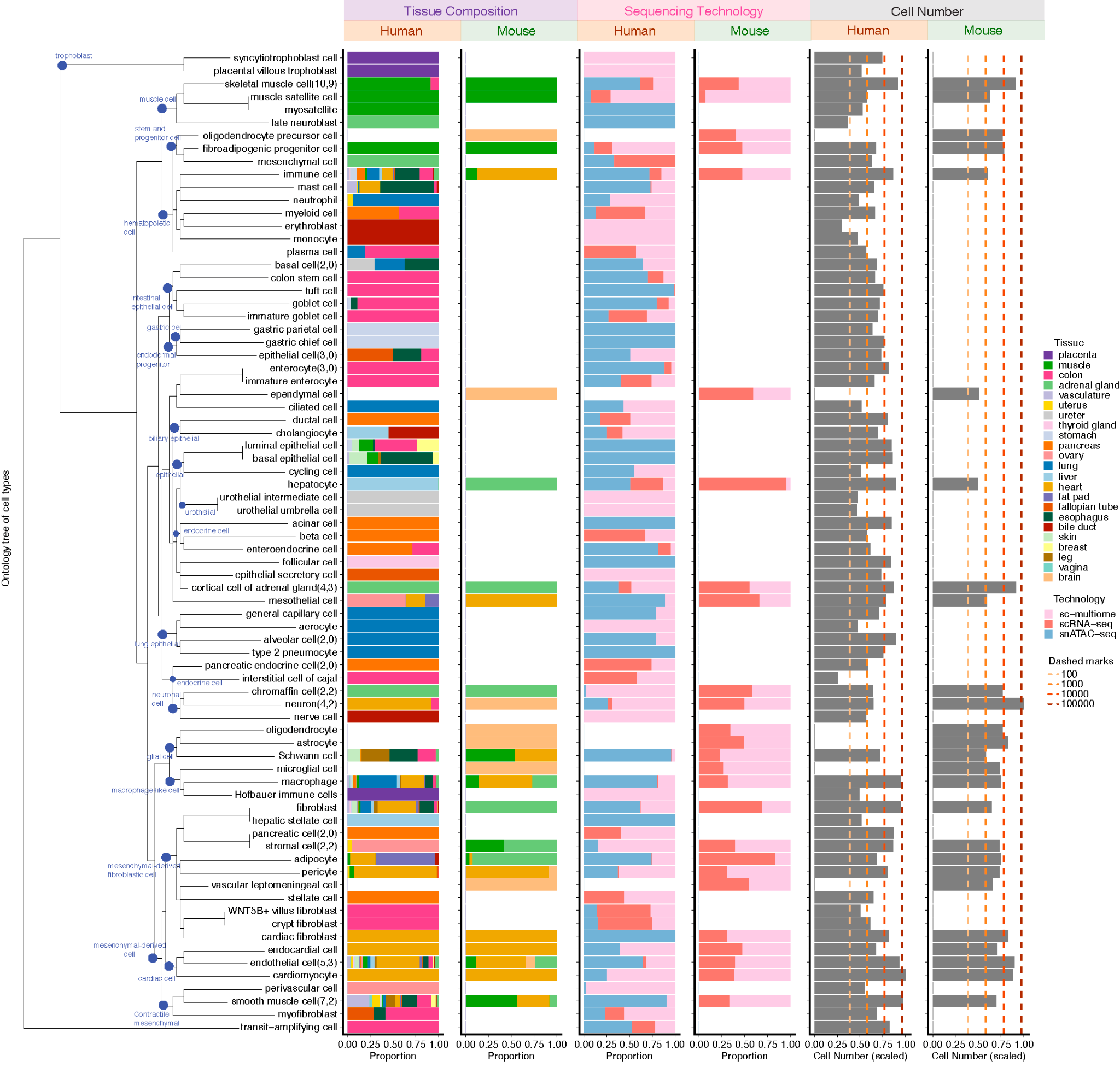

#### Supplementary Figure 1 | Overview of ENCODE Single Cell Datasets

**Supplementary Figure 1 | Overview of ENCODE Single Cell Datasets.** Cell types are harmonized and ordered according to hierarchical clustering based on pairwise distances derived from the Cell Ontology (CL) graph, as shown by the dendrogram on the left (ontology tree of cell types). The visualization includes 76 harmonized cell type labels used for cross-species comparison. Numbers in parentheses next to each cell type denote the counts of original subtype annotations contributing to the harmonized label (human, mouse). For each cell type, barplots summarize (from left to right): tissue composition in human and mouse, sequencing technology composition in human and mouse, and total cell numbers in human and mouse. Tissue proportions represent the fraction of cells from each tissue within a given cell type. Sequencing technology proportions represent the fraction of cells generated by scRNA-seq, snATAC-seq, or single-cell multiome profiling. Cell numbers are log10-scaled and normalized within species; dashed vertical lines indicate reference thresholds corresponding to 10^2^, 10^3^, 10^4^, and 10^5^ cells.

#### Supplementary Figure 2 | Summary of systematic approaches for silencer identification

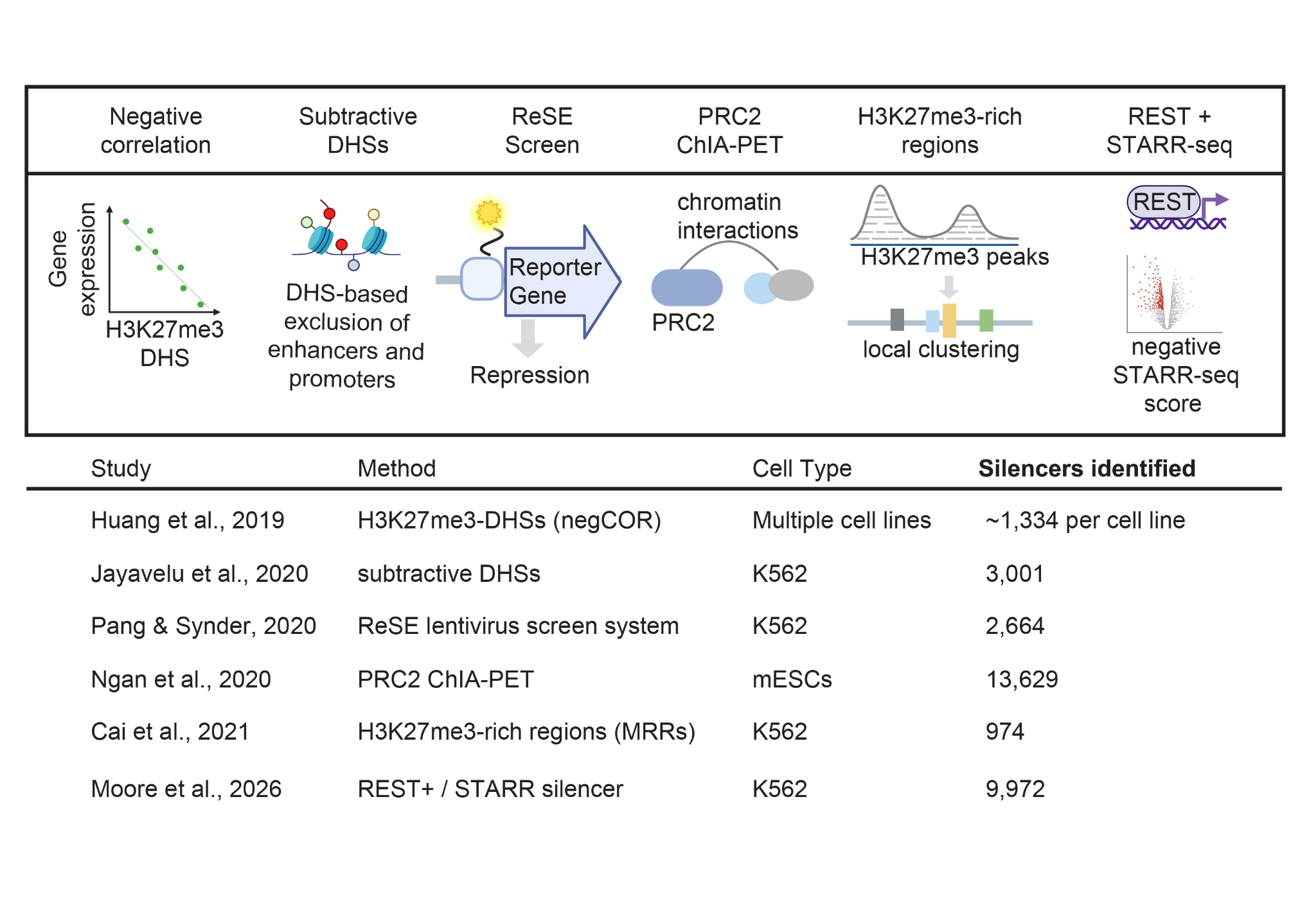

**Supplementary Figure 2 | Summary of systematic approaches for silencer identification.** Six representative methods are illustrated in this panel. Silencers can be identified based on: (1) H3K27me3-marked DNase I hypersensitive sites (DHSs) followed by negative correlation with gene expression; (2) subtractive approaches that exclude enhancer- and promoter-associated DHSs; (3) the ReSE lentiviral screening system; (4) PRC2 ChIA-PET, in which PRC2-bound loci are defined as silencers; (5) clustering of H3K27me3 signals to identify H3K27me3-rich regions (MRRs) as silencers; and (6) integration of REST transcription factor binding with negative STARR-seq activity. The cell type used for each method and the number of identified silencers is also indicated.

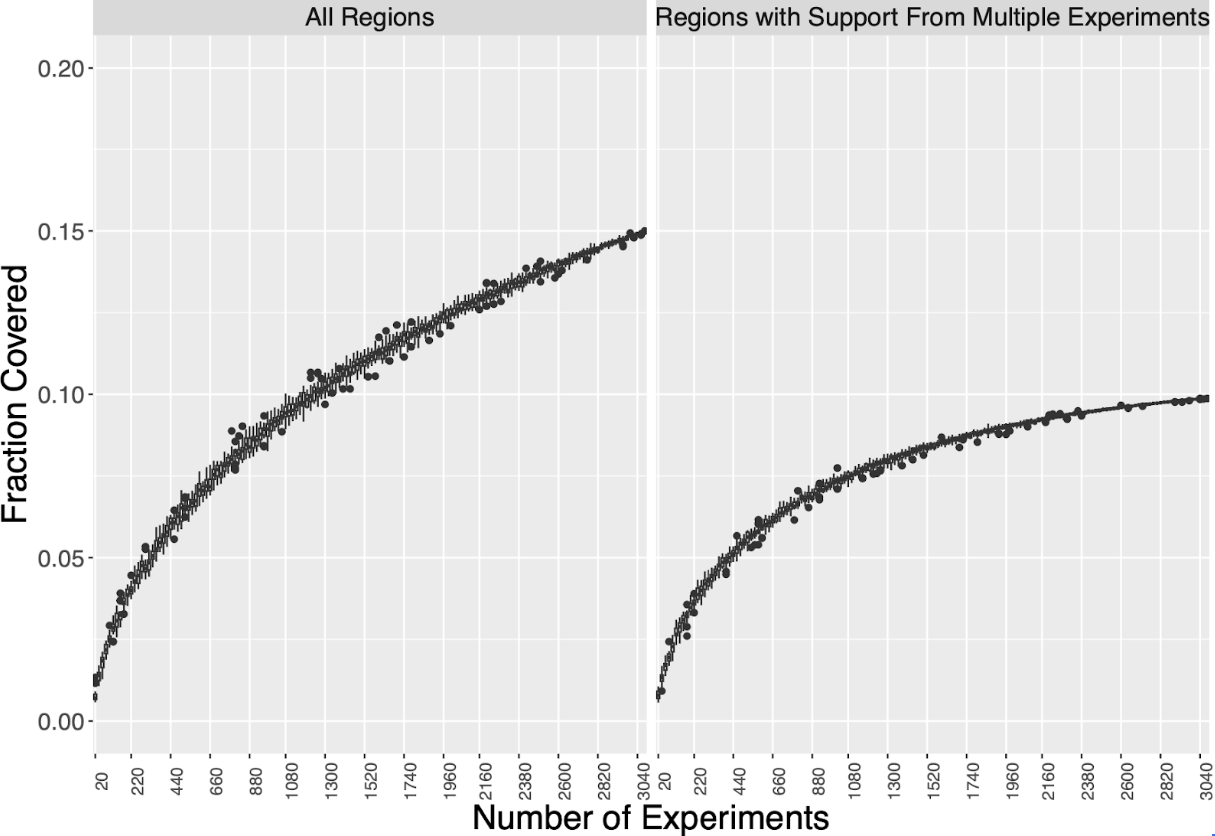

#### Supplementary Figure 3 | Fraction of the human genome bound by transcription factors

**Supplementary Figure 3 | Fraction of the human genome bound by transcription factors.** The fraction of the human genome bound by a TF, based on TF ChIP-seq. To avoid bias due to the order in which experiments were included, the ChIP-seq datasets were randomly subsampled to the number indicated on the x axis. ChIP-seq experiments were subsampled repeatedly, and for each subsampling peaks were restricted to the central 100 bp and merged to determine the fraction of the genome covered by at least one merged region. Left: Boxplot of genome coverage for each subset size of all experiments. Right: Boxplot of genome coverage for each subset, but restricted to regions which were covered by at least two experiments, to reduce the impact of false positive peak calls on the analysis.

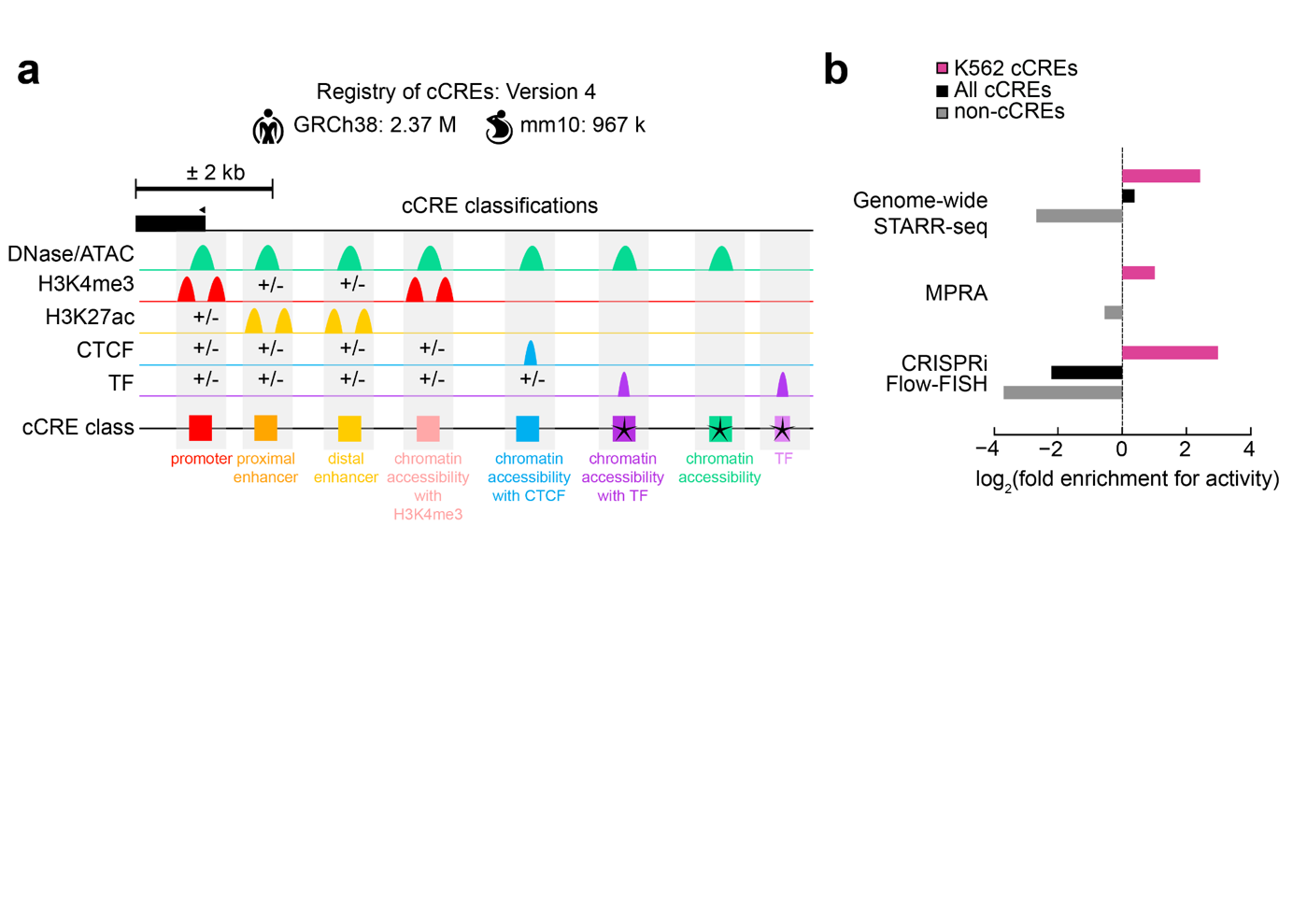

#### Supplementary Figure 4 | Overview of the cCRE catalog

**Supplementary Figure 4 | Overview of the cCRE catalog.** a, Overview of the ENCODE4 cCRE classification scheme. Candidate cis-regulatory elements (cCREs) are classified based on combinations of biochemical signals—chromatin accessibility (green), H3K4me3 (red), H3K27ac (yellow), CTCF binding (blue), and transcription factor binding (purple) — and their distance to annotated transcription start sites (TSSs). High signal levels are indicated by peaks. The ± symbols indicate that the corresponding signal may be present or absent and does not affect classification. Newly introduced cCRE classes in ENCODE4 are denoted by asterisks. In total, 2.37 million cCREs were identified in human (GRCh38) and 967,000 in mouse (mm10). b, Barplots depicting Log₂ fold enrichment of functional activity for regions tested by whole-genome STARR-seq (top), MPRA (middle), and CRISPRi FlowFISH (bottom). Regions are stratified by type: K562 cCREs (pink), cCREs defined in other cell types (black), and non-cCRE regions (gray). Enrichment is calculated as the fraction of active regions within each category relative to the overall fraction of active regions tested in the corresponding assay.

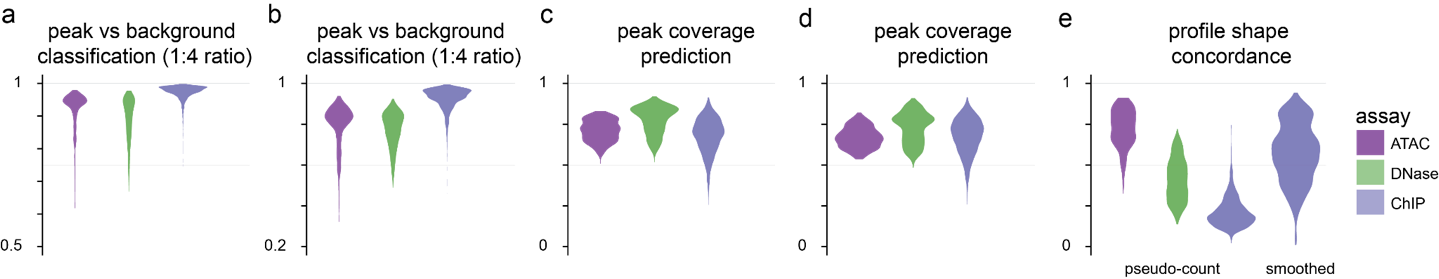

#### Supplementary Figure 5 | Cross-validation of BPNet and ChromBPNet models

**Supplementary Figure 5 | Cross-validated performance of BPNet and ChromBPNet models.** Violin plots summarize the distribution (across experiments) of the median performance over five cross-validation folds for models trained on TF ChIP-seq (BPNet) and chromatin accessibility (ChromBPNet; DNase/ATAC). For each fold, metrics were evaluated on held-out test chromosomes. **a,b.** Area under the receiver operating curve (auROC) (a) and area under the precision recall curve (auPR) (b) measure the performance of the models classifying high-confidence, reproducible peaks (positive set) against four times as many GC-matched background regions (negative set). Baseline random auROC = 0.5, auPR = 0.2. **c,d.** Peak-resolution coverage prediction performance was computed as the Pearson (c) and Spearman (d) correlations between predicted and observed total log(counts) across all peaks (1 kb bins at each peak). **e.** Profile-shape prediction was computed as the Pearson correlation between observed and predicted 1kb base-resolution probability profiles (after adding pseudocounts per position and/or smoothing (for ChIP-seq only due to extreme sparsity) at each peak. For each dataset/fold, we summarize the median of the profile-shape Pearson correlation over all peaks.

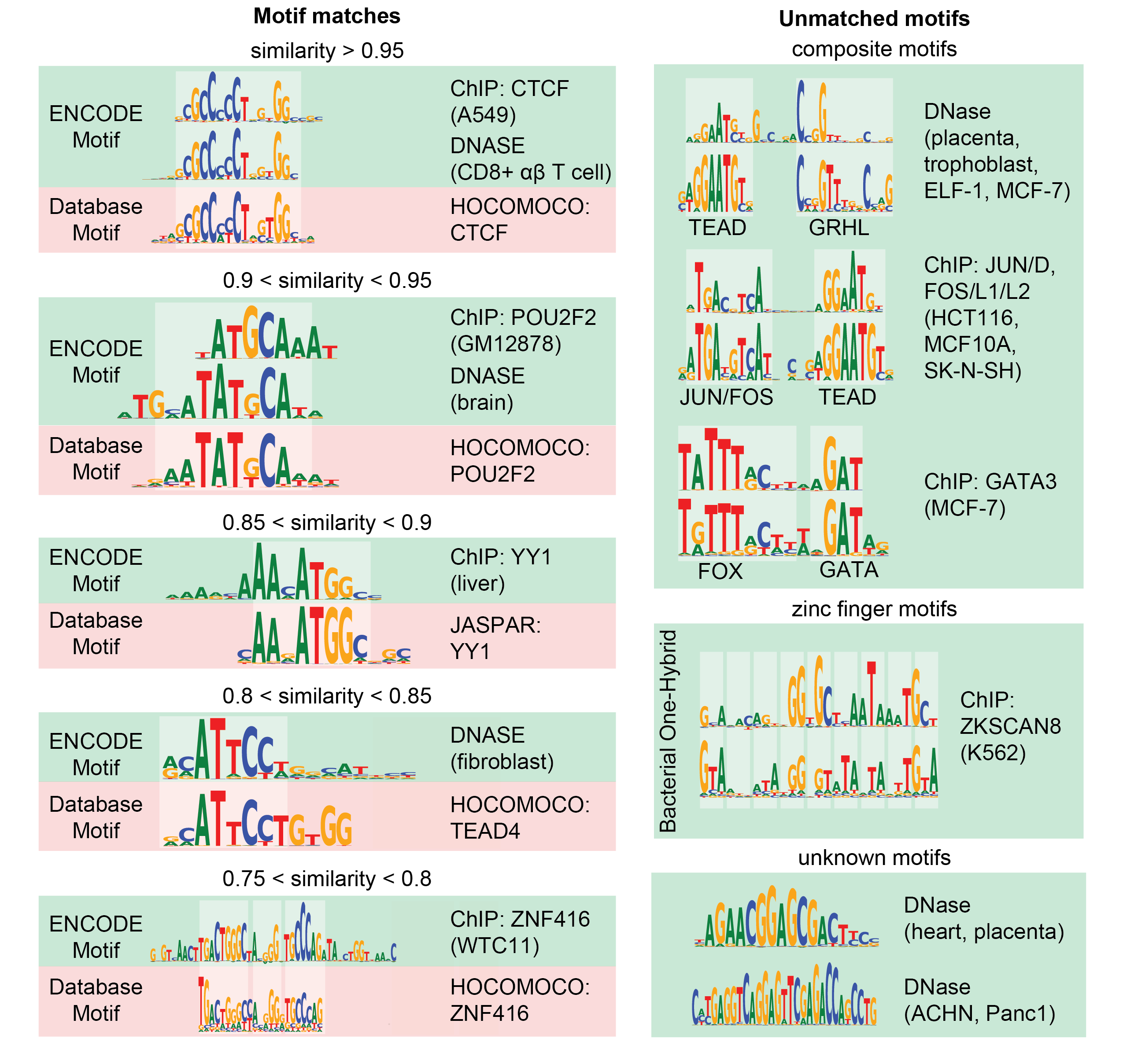

#### Supplementary Figure 6 | Examples of motif matches and novel motifs in the ENCODE motif catalog

**Supplementary Figure 6 | Examples of motif matches and novel motifs in the ENCODE motif catalog.** (Left column) Examples of motif matches across a range of similarity thresholds between 0.75 and 1.0. (Right column). Example motifs that are found only in the ENCODE motif catalog. Some of them are novel composite motifs of other known motifs and novel zinc finger motifs supported by alignment with the corresponding *in-vitro* bacterial-one-hybrid (B1H) binding profiles per finger.

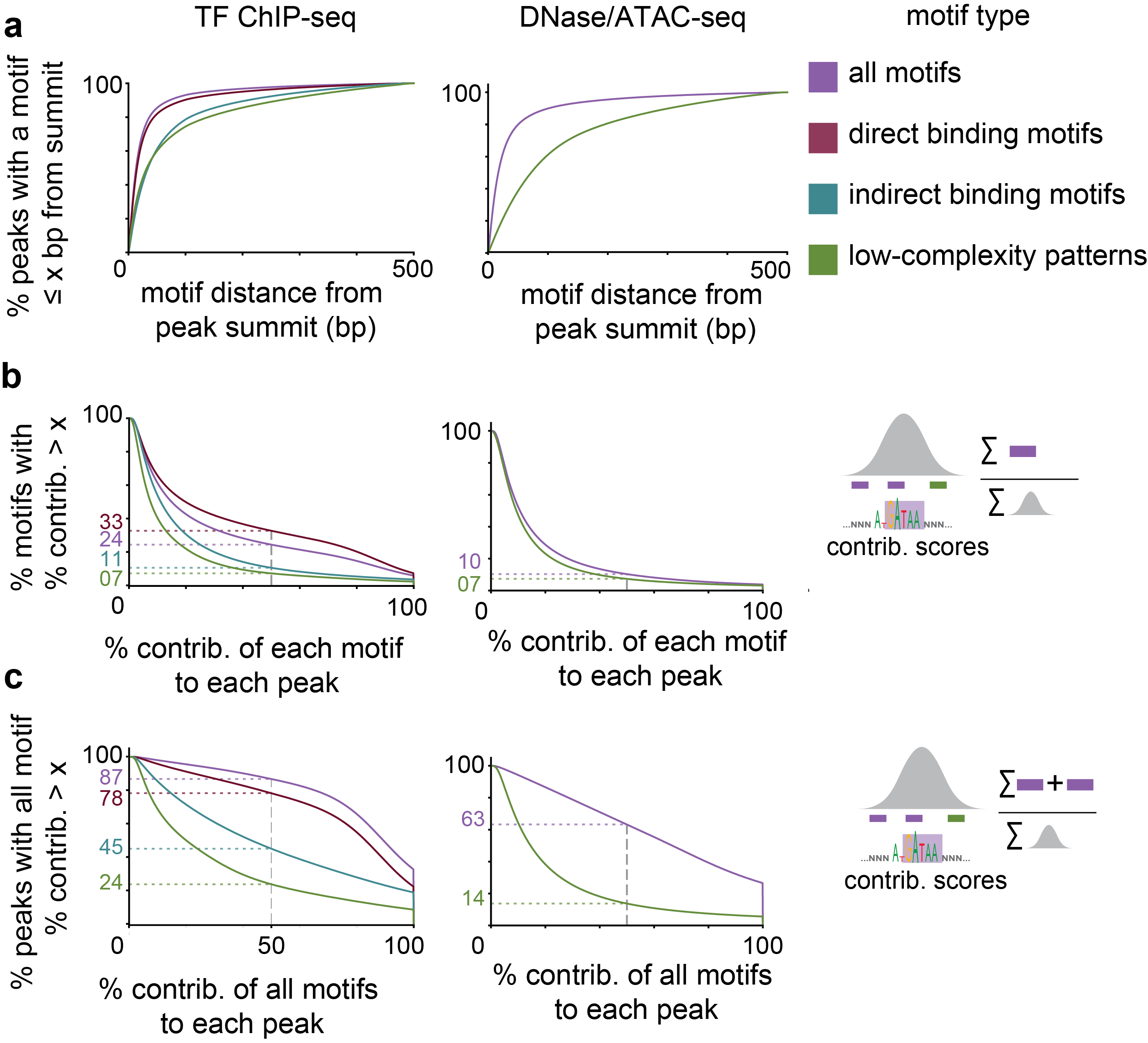

#### Supplementary Figure 7 | Localization and predictive contributions of model-derived motif instances in transcription factor binding and chromatin accessibility peaks

**Supplementary Figure 7 | Localization and predictive contributions of model-derived motif instances in TF binding and chromatin accessibility peaks.** Predictive motif instances are identified in every peak of each TF ChIP-seq and DNase-seq/ATAC-seq dataset using the corresponding BPNet or ChromBPNet deep learning model. Left and right panels correspond to TF ChIP-seq and DNase/ATAC-seq derived statistics respectively. **a**, Cumulative distribution of distance of predictive motif instances (P(distance <= x)) with respect to peak summits for different classes of motifs. When multiple motifs are present in a peak, the distance of the nearest predictive motif instance with respect to the peak summit for each class is used. The motif classes are as follows. all motifs: all motifs learned from each dataset except low complexity predictive patterns; direct binding motifs: for TF ChIP-seq datasets, these are motifs that match known cognate sequence preferences of the TF; indirect binding motifs: for TF ChIP-seq datasets, these are motifs that match cognate motifs of other TFs i.e. not the one targeted by the ChIP-seq experiment; low-complexity patterns: instances that match low complexity predictive patterns (see Methods). **b**, Complementary cumulative distribution of percentile contribution (P(% contrib > x)) of each predictive motif instance in peaks across all datasets for different classes of motifs. The guides show the % of instances of each motif class with > 50% contribution. For TF ChIP-seq, direct bound motifs have larger contributions than indirect bound motifs. **c**, Complementary cumulative distribution of percentile contribution (P(% contrib > x)) of all predictive motif instances in each peak across all datasets for different classes of motifs. The guides show the % of instances of each motif class with > 50% contribution.

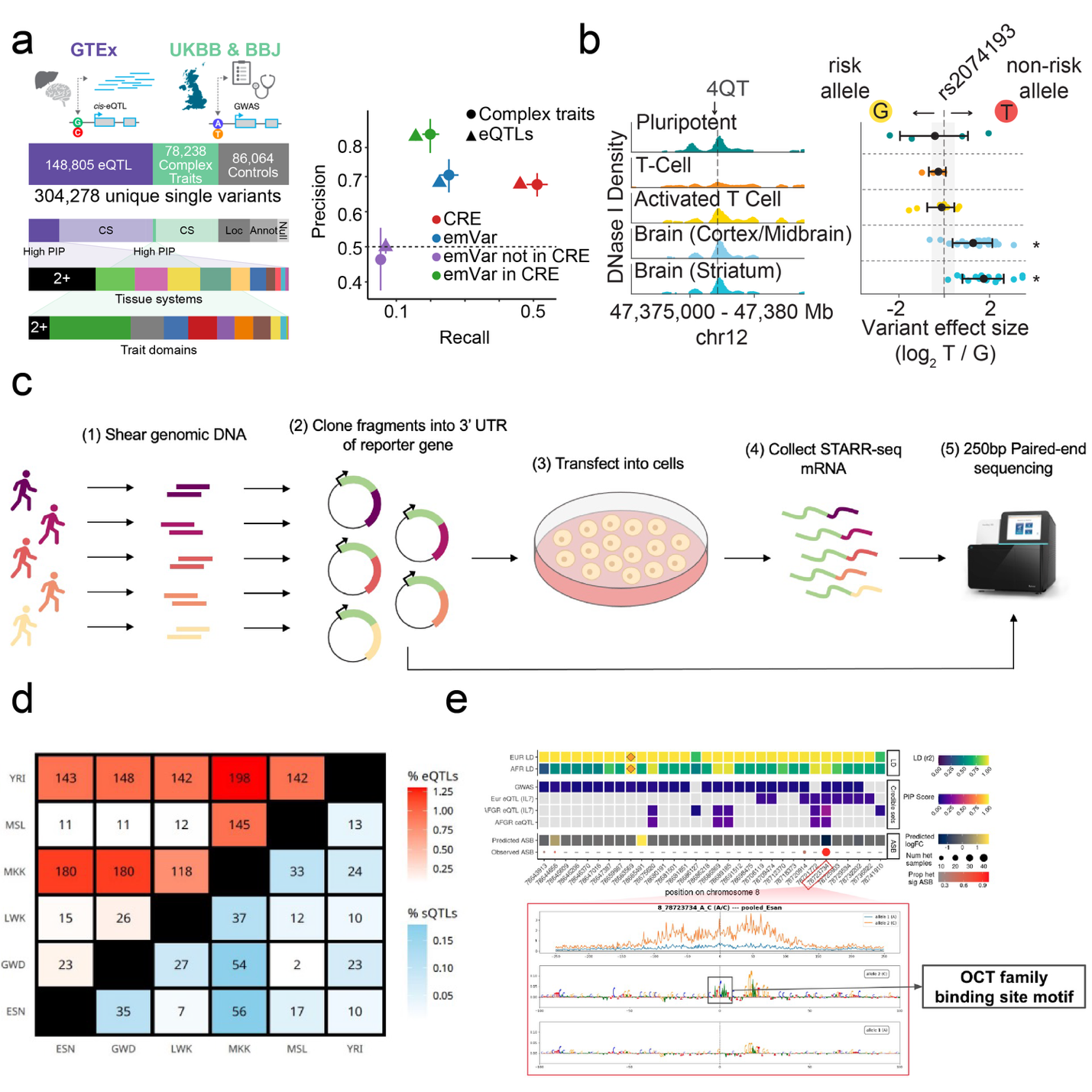

#### Supplementary Figure 8 | ENCODE approaches to estimate the effects of noncoding genetic variants

**Supplementary Figure 8 | ENCODE approaches to estimate the effects of noncoding genetic variants.** Here we detail the functional assays and analyses used in ENCODE 4 to estimate variant function. **a,** Massively Parallel Reporter Assays.^9^ (Left panel) Massively parallel reporter assays (MPRAs) were used to measure allele-specific expression-modulating effects across five different ENCODE cell lines of 221,412 unique fine-mapped trait-associated variants, consisting of GTEx eQTLs and GWAS variants from UK Biobank and Biobank Japan cohorts.^10^ (Right panel) A variant is called a significant expression modulating variant (emVar) if one or both alleles passes a log_2_ fold-change activity cut-off of 1 with a Bonferroni-adjusted activity p-value of 0.01 and activity is significantly different between alleles at an FDR-adjusted p-value of 0.1. (Right panel) The emVars, when combined with CREs (defined as accessible chromatin with an activating histone modification), showed higher precision than CREs albeit with lower recall, in fine-mapped disease variants and eQTLs. **b,** DNase I footprinting (Left panel) We performed in-vivo characterization of allelic imbalance using DNase I profiles across 1393 samples (491 distinct individuals). Variant calling, filtering, and statistical testing of allelic imbalance were performed using an improved protocol that corrects for mapping bias and copy number effects.^11,12^ Of the 1.7 million SNVs tested for allelic imbalance, 112,595 variants showed significant chromatin alterations (q-value ≤ 0.1). Allelic effects distribution across all variants highlights the symmetry in gain or loss in chromatin accessibility function. (Right panel) To investigate cell selective penetrance of regulatory variation, we used a data-driven approach to group samples into distinct cell archetypes and developed a statistical method to detect cell context-dependent variation, resulting in 124,971 variants imbalanced in at least one cell type. Here, we present an example of a cell context-dependent chromatin altering variant, *rs2074193*, a GWAS hit for migraine with aura.^13^ Although multiple cell types with an accessible DHS overlap this variant, the preference in chromatin accessibility towards the non-risk T allele is observed exclusively in brain samples. **c,** Genome-wide STARR-seq. We performed in the K562 cell line by generating pools of whole genomic DNA from five individuals per library. Alleles in regions with identified regulatory activity spanned common and rare variants and matched closely in their experimentally measured allele frequency and the expected population allele frequency. Variants were nominated as significant if they have a posterior probability > 0.90 for a Bayesian estimation of the genetic regulatory.^14^. Overall, out of 9,991,229 common and low-frequency variants, 483,023 were tested by these reporter assays, and we observed significant experimental evidence of allelic-level function for 7.9% of these variants. **d,** Quantitative trait loci. To increase the genetic diversity of variant effects, we used the African Functional Genomics Resource (AFGR) to perform QTL calling and allele-specific analyses.^15^ We observed 107,817 allele-specific expression (ASE) variants that were homozygous or not observed in 1000 Genomes European populations samples. Using the 100 individual ATAC-seq collections in AFGR, we detected allele-specific chromatin accessibility events for 299,247 unique variants. We observed a high replication rate of sharing in LCL eQTLs within African population subgroups and with the GEUVADIS European population (mashR $\pi_{1}$ranging from 0.63 to 0.94). **e,** DNA sequence models of function. ChromBPNet employs a bias-factorized model that regresses out chromatin background representing Tn5 enzyme bias in the non-peak profiles to glean the signal correctly. This model was used to annotate common and low-frequency variants (minor allele count >=5 in 1000 Genomes) for DNase I profiles in 104 samples and additional ATAC profiles in 4 ENCODE cell lines. Here, we present an example of a variant, *chr8:78723734:A:C*, which showed differences in ChromBPNet predicted activity between the two alleles, A and C, coinciding with the *Oct* family binding motif being altered by the variant.

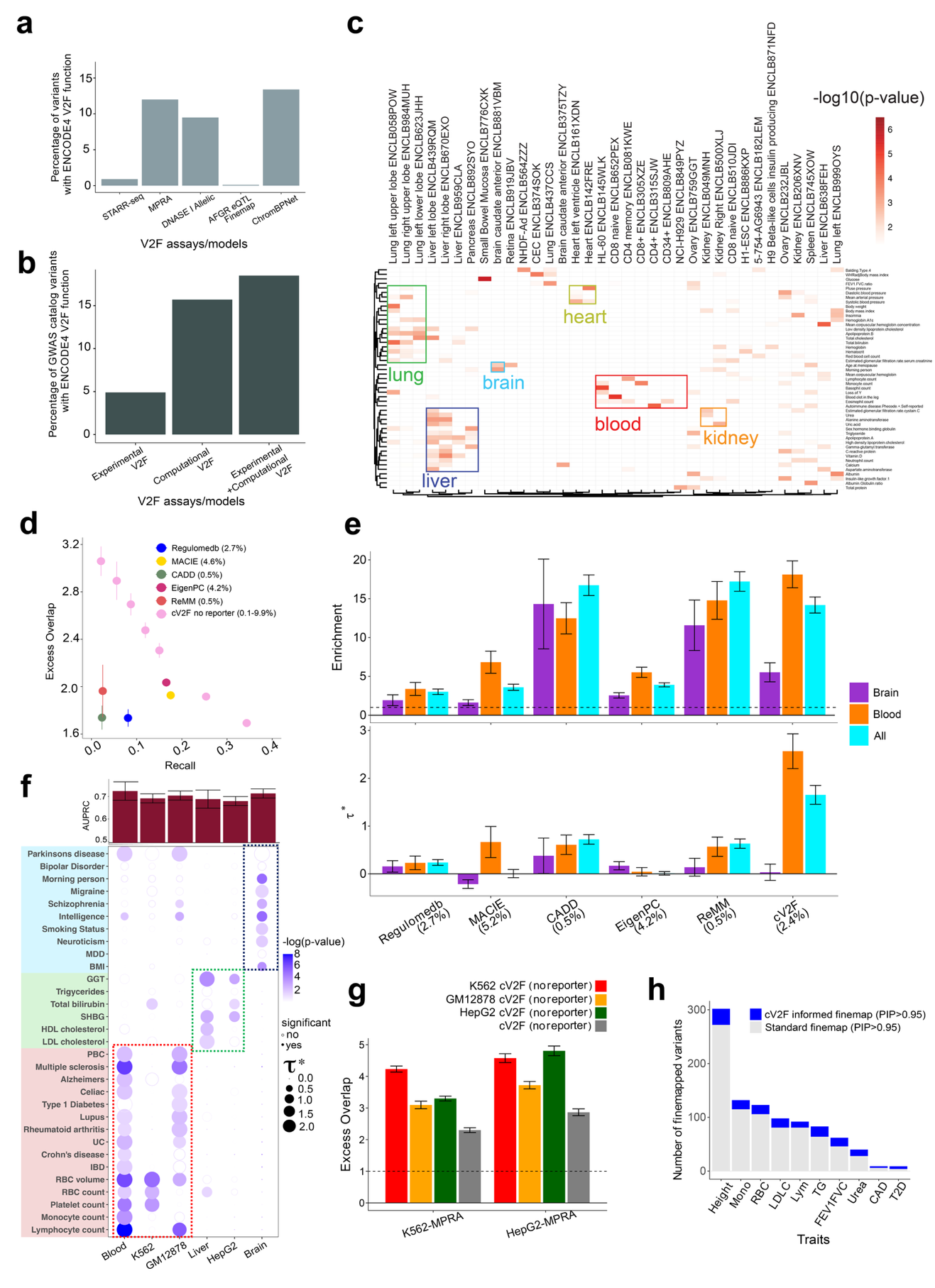

#### Supplementary Figure 9 | Variant-to-function (V2F) annotations for functional variant prioritization

**Supplementary Figure 9 | Variant-to-function (V2F) annotations for functional variant prioritization. a,** Percentage of common and low-frequency genetic variants (minor allele count >=5 in 1000 Genomes Project) with functional evidence from ENCODE4 experimental V2F assays and computational models. Experimental approaches include STARR-seq, MPRA, and DNase I allelic imbalance assays; computational approaches include fine-mapped eQTLs from African Functional Genomics Resource (AFGR), and ChromBPNet predictions. **b,** Cumulative percentage of GWAS Catalog variants annotated with V2F function using experimental assays alone, computational predictions alone, or the combination of both approaches. **c,** Tissue-specific enrichment of DNase allelic imbalance hits against fine-mapped variants across 50 complex diseases and traits. Heatmap displays −log_10_(p-value) of enrichment for each sample-trait pair, with samples grouped by tissue of origin (liver, lung, brain, blood, heart, kidney). **d,** Excess overlap and recall with respect to fine-mapped disease causal variants across 90 complex diseases and traits for cV2F (no reporter) subsetted at different thresholds and other existing scores (RegulomeDB, MACIE, CADD, EigenPC, ReMM). Error bars denote 95% confidence intervals. Percentages next to the method indicate the fraction of variants prioritized by each method. **e**, Heritability enrichment and standardized effect sizes of cV2F scores and other variant prioritization scores (RegulomeDB, MACIE, CADD, EigenPC, ReMM) - meta-analyzed across all traits, brain-related traits, and blood-related traits. The traits are selected to ensure they are not highly genetically correlated (rg < 0.7). Results are conditional on 97 baseline-LD v2.2 + 1 binarized primary cV2F annotations. Error bars denote 95% confidence intervals. See Supplementary Table 8 for the list of traits. **f**, Area under the Precision Recall curve (AUPRC) of cell-line and tissue-specific cV2F gradient boosting models based on held-out GWAS fine-mapping data. (Bottom panel) S-LDSC standardized effect sizes (*) of cell-line and tissue-specific cV2F scores for a set of related τ diseases and traits. Results are conditional on 97 baseline-LD v2.2 + 1 binarized primary cV2F annotations. Magnitude (𝛕*, dot size) and significance (−log_10_(P), dot color) are reported for disease signals for 31 blood, liver and brain-related traits. **g,** Excess overlap of binarized cell-line specific binarized cV2F-noReporter (model trained without MPRA features) variants for three cell lines- K562, GM12878, and HepG2 (colored), and primary cV2F-noReporter (gray) against MPRA positives versus tested variants in related cell lines (K562 and HepG2). Error bars denote 95% confidence intervals. **h,** Number of variants confidently fine-mapped (posterior probability of causality > 0.95) from standard fine-mapping and cV2F-informed functional fine-mapping of 10 selected diseases and traits covering the overall spectrum of relative improvement in fine-mapping across all 94 UKBB traits. See Supplementary Table 8 for the list of traits.

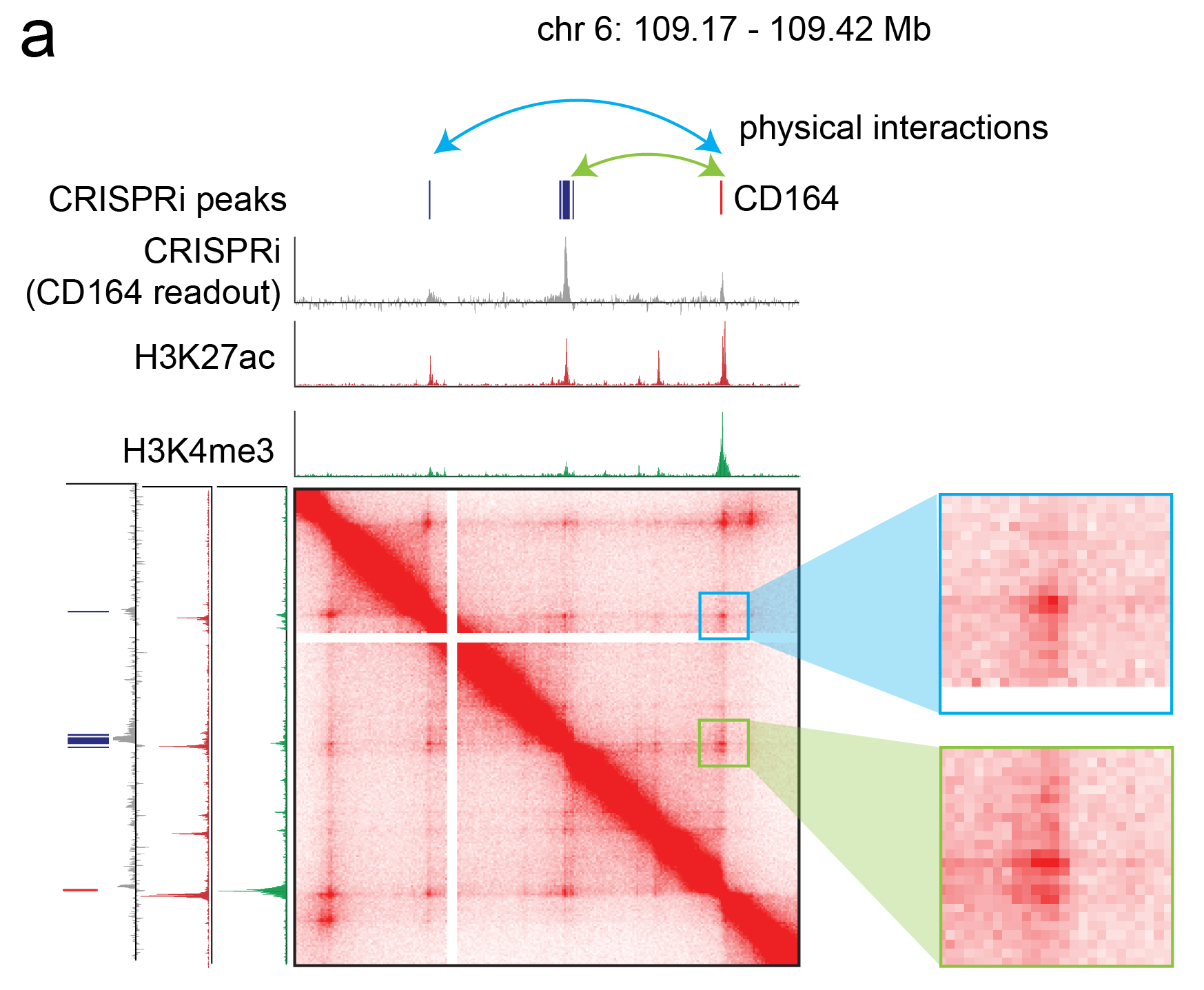

#### Supplementary Figure 10 | Example of overlapping physical interactions and regulatory interactions

**Supplementary Figure 10 | Physical looping interactions between enhancers and promoters detected by intact Hi-C overlap regulatory interactions detected by CRISPRi**. Intact Hi-C contact map in K562 cells is shown at the CD164 locus (chromosome 6:109.17-109.42 Mb) along with CRISPRi peaks and readout tracks, and ChIP-Seq tracks for H3K27ac and H3K4me3 histone modifications. Intact Hi-C identifies overlapping physical interactions between CRISPRi validated distal regulatory elements and the CD164 promoter.

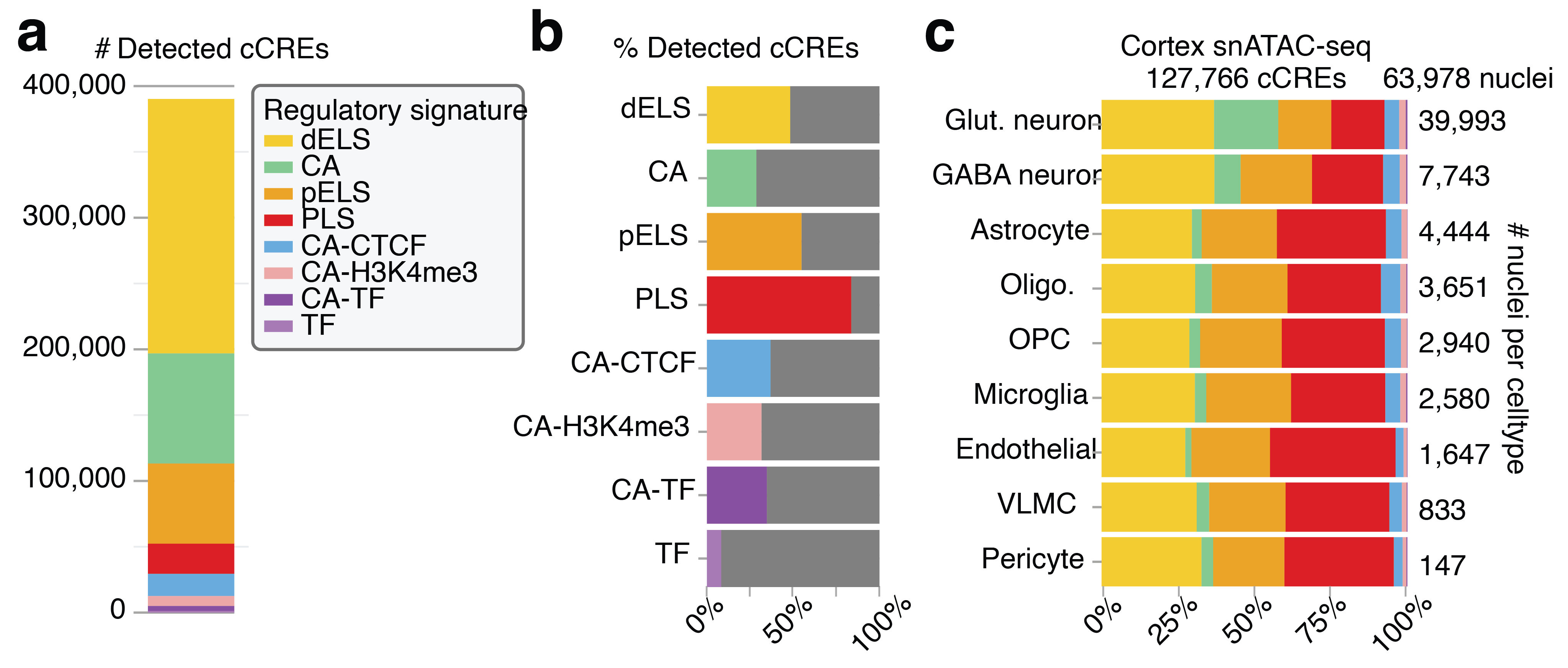

#### Supplementary Figure 11 | Distribution of cCRE annotations in pseudobulked mouse single cell ATAC-seq studies

**Supplementary Figure 11 | Distribution of cCRE annotations in pseudobulked mouse single cell ATAC-seq studies.** We found that 42% of ENCODE cCREs^16^ had chromatin accessibility in one or more of the mouse cell types studied in ENCODE. **a-b,** Breakdown of detected cCREs by type, with most promoter-like signature (PLS) detected. 390,146 out of 926,843 mouse cCREs were detected in one or more cell types in the snATAC-seq data. Most were distal enhancer-like sequences, and a substantial fraction of promoter-like sequences were also detected. **c,** Across all cell types, we observed a stable distribution of enhancer- and promoter-like sequences. However, a significant proportion of detected cCREs in glutamatergic neurons in the cerebral cortex are in accessible chromatin but not classified as traditional promoters or enhancers. Those likely are a result of the higher recovery of nuclei for glutamatergic neurons, allowing for more sensitive detection of regulatory elements that have yet to be well characterized by other assays. They may also indicate neuron-specific enhancers or promoters of genes that are not broadly expressed, or whose signals might be less prominent in bulk data. Single-nucleus ATAC-seq provides the resolution to uncover these subtleties, allowing us to parse the specific regulatory landscapes of different cell types and developmental stages.

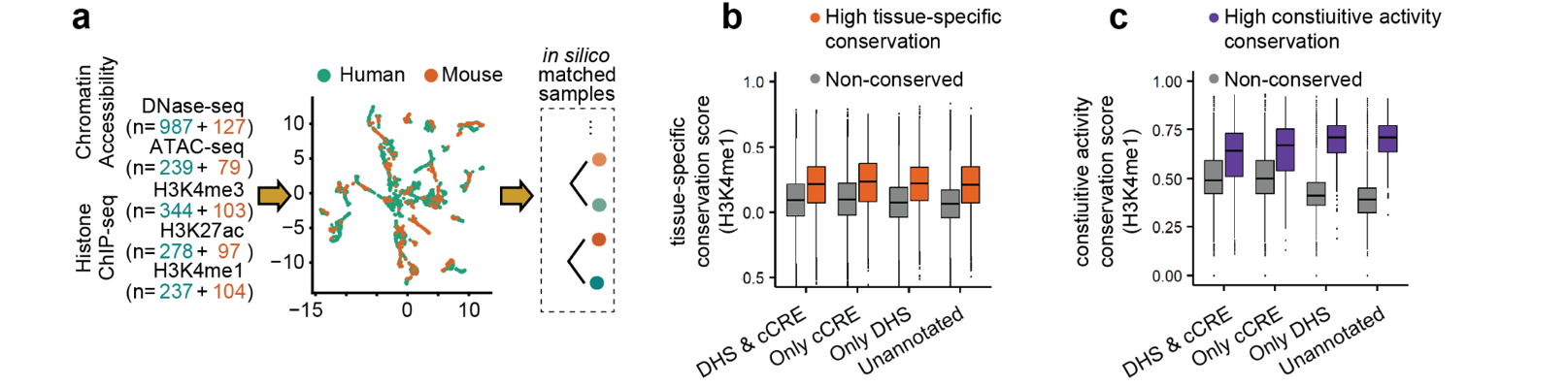

#### Supplementary Figure 12 | Sample matching for functional conservation and using FUNCODE to annotate new regulatory elements

**Supplementary Figure 12 | Sample matching for functional conservation and using FUNCODE to annotate new regulatory elements. a,** Construction of matched samples for scoring functional conservation. DNase-seq, ATAC-seq, and ChIP-seq profiles were co-embedded into a shared low-dimensional space to generate in silico–matched samples. Numbers of human and mouse samples are shown in parentheses. **b,** Identification of new regulatory elements using conservation of tissue-specific activity. Mouse DNA elements aligned to human DHSs or cCREs were grouped by mouse annotation status (annotated as both DHS and cCRE, only cCRE, only DHS, or unannotated). Within each group, elements were further classified as conserved (orange) or non-conserved (gray) based on the FUNCODE score for tissue-specific chromatin accessibility. Unannotated elements with conserved chromatin accessibility were predicted as putative regulatory elements. Boxplots show distributions of FUNCODE scores for tissue-specific H3K4me1, an orthogonal modality used to validate predictions based on chromatin accessibility. Unannotated mouse elements predicted to be regulatory elements exhibit higher conservation of H3K4me1 activity than annotated but non-conserved elements. **c,** Identification of new regulatory elements using conservation of constitutive activity. The FUNCODE score for constitutive chromatin accessibility was used to identify unannotated mouse elements with conserved constitutive activity as putative regulatory elements. These predictions were validated using conservation of constitutive H3K4me1 activity, analogous to **b.**

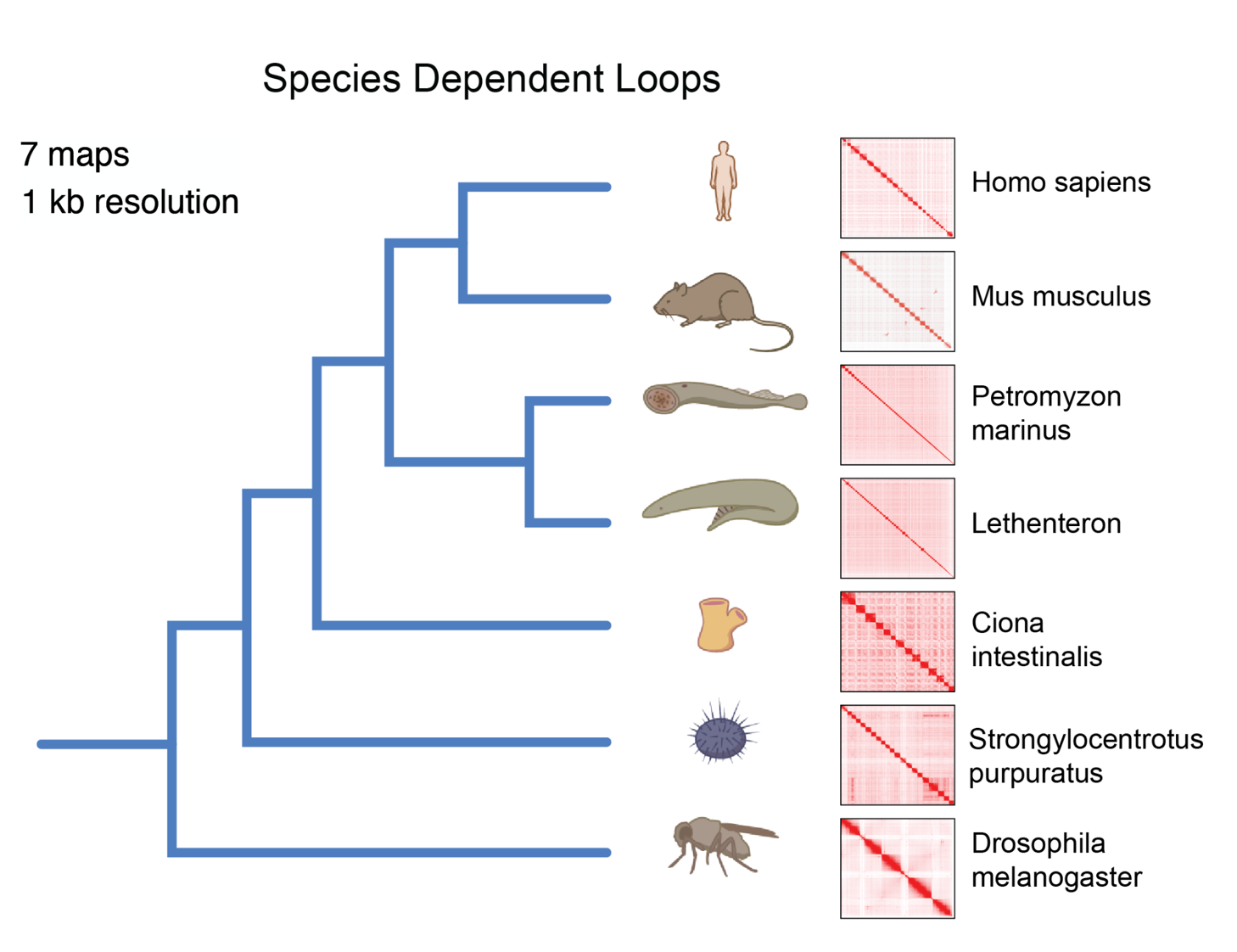

#### Supplementary Figure 13 | Long-range physical interactions evaluated over evolution

**Supplementary Figure 13 | Long-range physical interactions evaluated over evolution.** Physical interactions mapped across and beyond vertebrate evolution. In vertebrates, cohesin is arrested by inward-pointing CTCF binding sites. This behavior is not seen outside vertebrates.

### **Supplementary Tables**

#### Supplementary Table 1 | ENCODE 4 Data Summary

This table is provided as an external file.

#### Supplementary Table 2 | Overview of ENCODE 4 Single Cell Analysis

|  | **Single-cell multi-omics** | **Single-cell RNA-seq** | **Single-nucleus ATAC-seq** | **Total** |
| --- | --- | --- | --- | --- |
| **Total number of cells** | **584,168** | **269,369** | **650,694** | **1,504,231** |
| **Human cells** | **358,854** | **74,307** | **650,694** | **1,083,855 (72.05%)** |
| **Mouse cells** | **225,314** | **195,062** | **0** | **420,376 (27.95%)** |
| **Female cells** | **316,569** | **144,596** | **360,406** | **821,571 (54.62%)** |
| **Male cells** | **267,599** | **124,773** | **290,288** | **682,660 (45.38%)** |
| **Number of distinct cell types** | **105** | **71** | **70** | **120** |
| **Number of distinct tissue types** | **28** | **11** | **37** | **53** |
| **Number of distinct organs** | **14** | **7** | **18** | **23** |

#### Supplementary Table 3 | Datasets for ChIP-seq of tagged transcription factors

This table is provided as an external file.

#### Supplementary Table 4 | Deep learning models of nucleotide contributions

This table is provided as an external file.

#### Supplementary Table 5 | Enrichment of ChromBPNet variant effect predictions in variants with an allelic effect in MPRA

| Enrichment | sd (Enrichment) | Method (Cell line) | Assay (Cell line) |
| --- | --- | --- | --- |
| 11.8 | 1.4 | ChromBPNet-K562 | MPRA-K562 |
| 7.1 | 1.3 | ChromBPNet-GM12878 | MPRA-K562 |
| 6.1 | 1.1 | ChromBPNet-HepG2 | MPRA-K562 |
| 12 | 2.1 | ChromBPNet-K562 | MPRA-GM12878 |
| 15.5 | 2.6 | ChromBPNet-GM12878 | MPRA-GM12878 |
| 9.3 | 1.7 | ChromBPNet-HepG2 | MPRA-GM12878 |
| 10.6 | 1.8 | ChromBPNet-K562 | MPRA-HepG2 |
| 8.5 | 1.4 | ChromBPNet-GM12878 | MPRA-HepG2 |
| 10.4 | 1.8 | ChromBPNet-HepG2 | MPRA-HepG2 |

#### Supplementary Table 6 | Enrichment of ChromBPNet variant effect predictions in variants with an allelic imbalance in DNase hypersensitivity

| Enrichment | sd (Enrichment) | Method (Cell line) | Assay (Cell line) |
| --- | --- | --- | --- |
| 5.9 | 0.45 | ChromBPNet-K562 | DNaseAI-K562 |
| 3.5 | 0.35 | ChromBPNet-GM12878 | DNaseAI-K562 |
| 3.3 | 0.38 | ChromBPNet-HepG2 | DNaseAI-K562 |
| 6.78 | 0.6 | ChromBPNet-K562 | DNaseAI-GM12878 |
| 9.75 | 1.7 | ChromBPNet-GM12878 | DNaseAI-GM12878 |
| 5.58 | 0.5 | ChromBPNet-HepG2 | DNaseAI-GM12878 |
| 8.3 | 1.2 | ChromBPNet-K562 | DNaseAI-HepG2 |
| 8.2 | 1.4 | ChromBPNet-GM12878 | DNaseAI-HepG2 |
| 14.7 | 4.8 | ChromBPNet-HepG2 | DNaseAI-HepG2 |

#### Supplementary Table 7 | Annotation of GWAS Catalog variants with evidence of allele specific activity

This table is provided as an external file.

#### Supplementary Table 8 | Disease traits used in cV2F evaluation

This table is provided as an external file.

#### Supplementary Table 9 | Validation of cV2F using UK Biobank and Million Veterans Program traits

This table is provided as an external file.

#### Supplementary Table 10 | Percent improvement of cV2F-informed multi-ancestry polygenic prediction over a non-functional baseline

|  | Percent improvement in predictive performance | |  |
| --- | --- | --- | --- |
| Trait | All | Tissue-matched | Tissue |
| FEV1/FVC Ratio | 22.76% | 36.51% | Lung |
| Lymphocyte Count | 10.04% | 35.13% | Blood |
| eGFR | -11.00% | 23.72% | Kidney |
| LDL-C | 8.28% | 8.71% | Liver |

#### Supplementary Table 11 | Datasets for Evolutionary Hi-C Analysis

This table is provided as an external file.

### Case Studies

#### Case Study 1 | Pleiotropic contributions of DNA sequences to regulatory elements that control MYC expression

We illustrate how model-derived tracks—bias-corrected predicted signal, base-resolution contribution maps, and motif-instance annotations—enable mechanistic hypotheses about the cis-regulatory code at the *MYC* locus in K562 and HepG2 cells. We focus on the *MYC* promoter and two CRISPR-i validated distal enhancers (enhancer 1, enhancer 2) (**Supplementary Figure 14a**).

At enhancer 1 (**Supplementary Figure 14c**; details in **Supplementary Figure 15**), DNase-seq and ATAC-seq models in K562 recover a consistent accessibility profile only after correcting assay-specific enzyme biases. Sequence contribution maps from both models implicate a dominant GATA motif near the center of the enhancer—coincident with the local dip/footprint in accessibility—along with flanking GATA, SP and ETV motifs. MPRA models in K562 emphasized the same motifs but with stronger ETV effect sizes. Models of GATA1, SP1, and ETV6 ChIP-seq data in K562 supported these motif instances. The dominant central GATA motif was also predicted to strongly influence the occupancy of several other transcription factors (SP1, STAT6, ETV6, TCF7), consistent with a pioneer-like role for GATA1 in K562. By contrast, models of CTCF, GABPB1, KLF1, and SRF primarily highlighted their cognate motifs, showing limited coupling to accessibility and MPRA activity at this locus. Predictions were highly reproducible across ChIP-seq models of the same TFs despite differences in antibodies and data quality (**Supplementary Figure 16**).

At a more promoter-proximal enhancer 2 (**Supplementary Figure 14b**; details in **Supplementary Figure 16**), different motifs explain chromatin accessibility in K562 and HepG2 cells, consistent with the different transcription factors expressed in each. In K562, sequence contribution maps highlight a central GATA site (aligned with the strongest footprint) together with SP, AP-1 and ETS motifs, and TF binding models support these sites. In HepG2, the GATA motifs that were predictive in K562 are replaced by an HNF4 motif and a weak overlapping FOXA1/FOXA2 motif, while shared AP-1, SP and ETV sites remain predictive.

At the MYC promoter (**Supplementary Figure 14d**; details in **Supplementary Figure 17**), the nascent transcription (PRO-cap) models resolve two punctate TSSs with canonical initiation syntax (Inr with upstream TATA and SP/BRE).^17^ In contrast, accessibility models highlight an SP/CpG-rich grammar in both K562 and HepG2, with an additional K562-specific GATA site near the stronger TSS. TF binding models in both cell types support these motifs and point to additional TF-specific motifs such as non-canonical extended CTCF sites that only influence CTCF occupancy. Together, these models provide testable hypotheses of the pleiotropic DNA sequence features influencing diverse biochemical profiles at regulatory elements in different contexts.

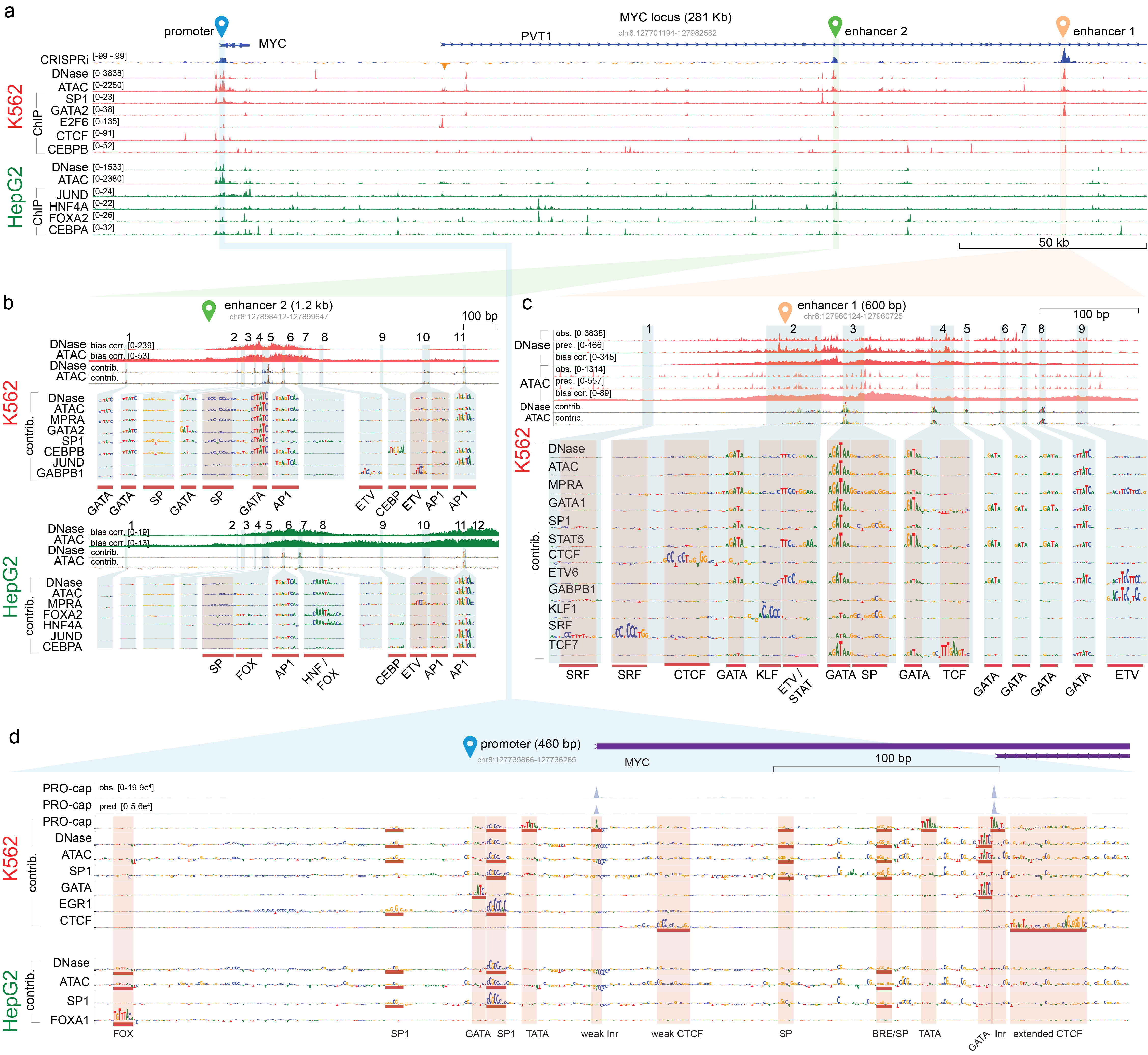

##### Supplementary Figure 14 | Deciphering pleiotropic regulatory sequence logic at the MYC locus with predictive sequence models

Deep learning models trained on diverse assays generate bias-corrected predicted signal, base-resolution sequence contribution maps, and motif-instance annotations, enabling mechanistic comparison of regulatory grammar across K562 and HepG2. (a) Genome browser view across the MYC/PVT1 locus (chr8:127,701,194–127,982,582) showing observed normalized signal tracks (y-axis units: MACS2 smoothed fold-enrichment) for chromatin accessibility (DNase-seq, ATAC-seq) and representative TF ChIP-seq experiments in K562 (red) and HepG2 (green). The CRISPRi tiling effect-size track (blue) highlights the MYC promoter and two distal enhancers (enhancer 1 and enhancer 2) as the strongest regulators of MYC expression in K562. (b) Zoom-in of enhancer 2 (chr8:127,898,412–127,899,647; ~162.1 kb downstream), accessible in both cell types. K562 (top): ChromBPNet predicts highly concordant bias-corrected (bias. corr) DNase- and ATAC-seq profiles (y-axis units: base-resolution fold-enrichment) and concordant contribution maps (contrib.), with annotated high-impact motifs (e.g., GATA, SP, AP-1, ETV/ETS, CEBP) supported by ChromBPNet, ReporterNet (MPRA), and selected BPNet TF ChIP-seq models. HepG2 (bottom): Predicted bias-corrected profiles and contribution maps differ from K562, indicating cell-type–specific logic: shared AP-1/SP/ETV sites remain predictive, K562 GATA contributions are lost, and HepG2-specific HNF/FOX motifs emerge, supported by BPNet models (e.g., HNF4A, FOXA2). (c) Zoom-in at K562-specific enhancer 1 (chr8:127,960,124-127,960,725, ~223.8 kb downstream of promoter) in K562. DNase-seq and ATAC-seq show strong K562 accessibility but discordant observed base-resolution profiles (obs.); assay-specific ChromBPNet predictions (pred.) without bias-correction recapitulate these differences. After bias correction, DNase/ATAC predicted profiles (bias corr.) and sequence contribution maps (contrib.) become highly concordant. Zoomed contribution maps from ChromBPNet, ReporterNet, and BPNet highlight a dominant footprint-aligned GATA site with additional GATA, ETV, SP, and TF-specific motifs. (d) Zoom-in at the MYC promoter (chr8:127,735,867-127,736,285). K562 (top): Observed (obs.) and ProCapNet-predicted (pred.) PRO-cap profiles resolve two punctate TSSs; ProCapNet sequence contribution maps (contrib.) highlight initiation syntax (Inr/TATA/SP-BRE), whereas ChromBPNet and BPNet TF ChIP-seq contributions emphasize distinct accessibility/occupancy determinants dominated by SP/CpG-like features. HepG2 (bottom): ChromBPNet sequence contribution maps are broadly similar to K562 but lack K562-specific GATA contributions.

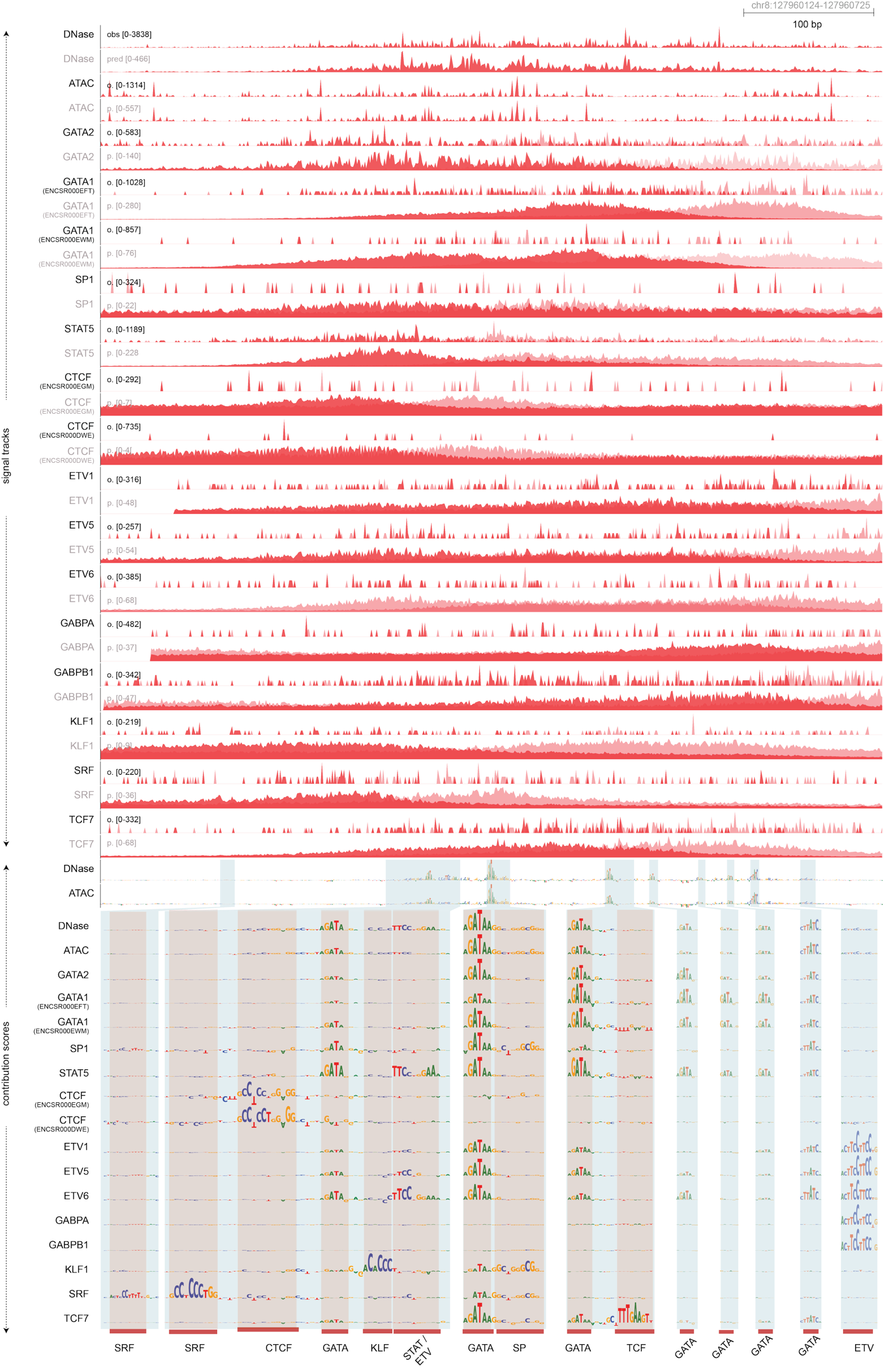

##### Supplementary Figure 15 | Model-derived predicted profiles and sequence contribution maps resolve reproducible sequence logic and reconcile assay-specific artifacts at MYC enhancer 1

Tracks across MYC enhancer 1 (chr8:127,960,124–127,960,725) in K562 showing observed (track labels in black) DNase-seq, ATAC-seq, and TF ChIP-seq base-resolution profiles (base-resolution fold-enrichment), alongside corresponding model predictions (track labels in light grey) and sequence contribution maps from ChromBPNet (DNase/ATAC) and BPNet (ChIP-seq). ChIP-seq tracks are shown stranded (positive/negative strand coverage in dark/light red). Despite sparse ChIP-seq signal, BPNet imputes smooth, high-resolution profiles and yields highly consistent predicted profiles and contribution maps across replicate experiments and/or related TFs where available, even when the observed profiles differ markedly. The lower panel summarizes ChromBPNet and BPNet contribution maps at enh1, with predictive motif instances annotated beneath the tracks, highlighting a dominant footprint-aligned GATA site and additional ETS/SP and TF-specific determinants that explain accessibility and TF occupancy.

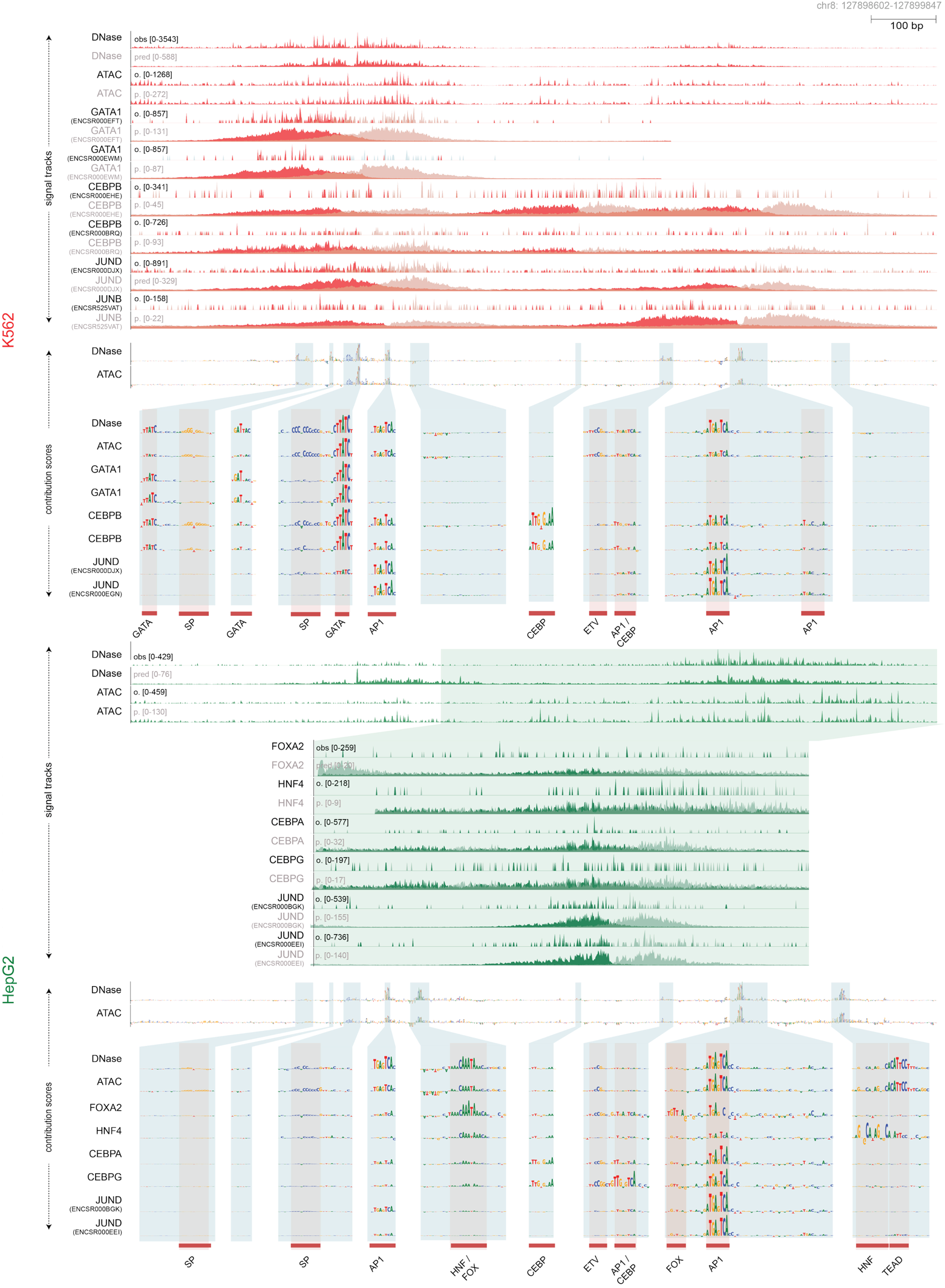

##### Supplementary Figure 16 | Model-derived profiles and contribution maps reveal cell-type rewiring of enhancer syntax at MYC enhancer 2

Tracks across MYC enhancer 2 (chr8:127,898,412–127,899,647) comparing K562 and HepG2 showing observed (track labels in black) DNase-seq, ATAC-seq, and TF ChIP-seq profiles (base-resolution fold-enrichment), alongside corresponding model predictions (track labels in light grey) and sequence contribution maps from ChromBPNet (DNase/ATAC) and BPNet (TF ChIP-seq). ChIP-seq tracks are shown stranded (positive/negative strand coverage in dark/light pink). Despite sparse and experiment-variable ChIP-seq signal, BPNet imputes smooth, high-resolution ChIP-seq profiles and produces more concordant predicted profiles and contribution maps across replicate experiments and/or related TFs where available. ChromBPNet reconciles assay-specific DNase/ATAC signatures via bias correction and yields coherent contribution maps within each cell type. In K562, contribution maps highlight a shared motif grammar dominated by a central footprint-aligned GATA site with additional SP, AP-1, and ETV motifs supported by BPNet models for relevant TFs. In HepG2, the enhancer remains accessible but the predicted profiles and contribution maps shift. The GATA motifs are no longer predictive in HepG2. Instead, we observe increased contributions from HNF/FOXA-family motifs alongside shared AP-1/SP/ETS features, consistent with cell-type–specific regulatory logic. Predictive motif instances are annotated beneath the contribution tracks.

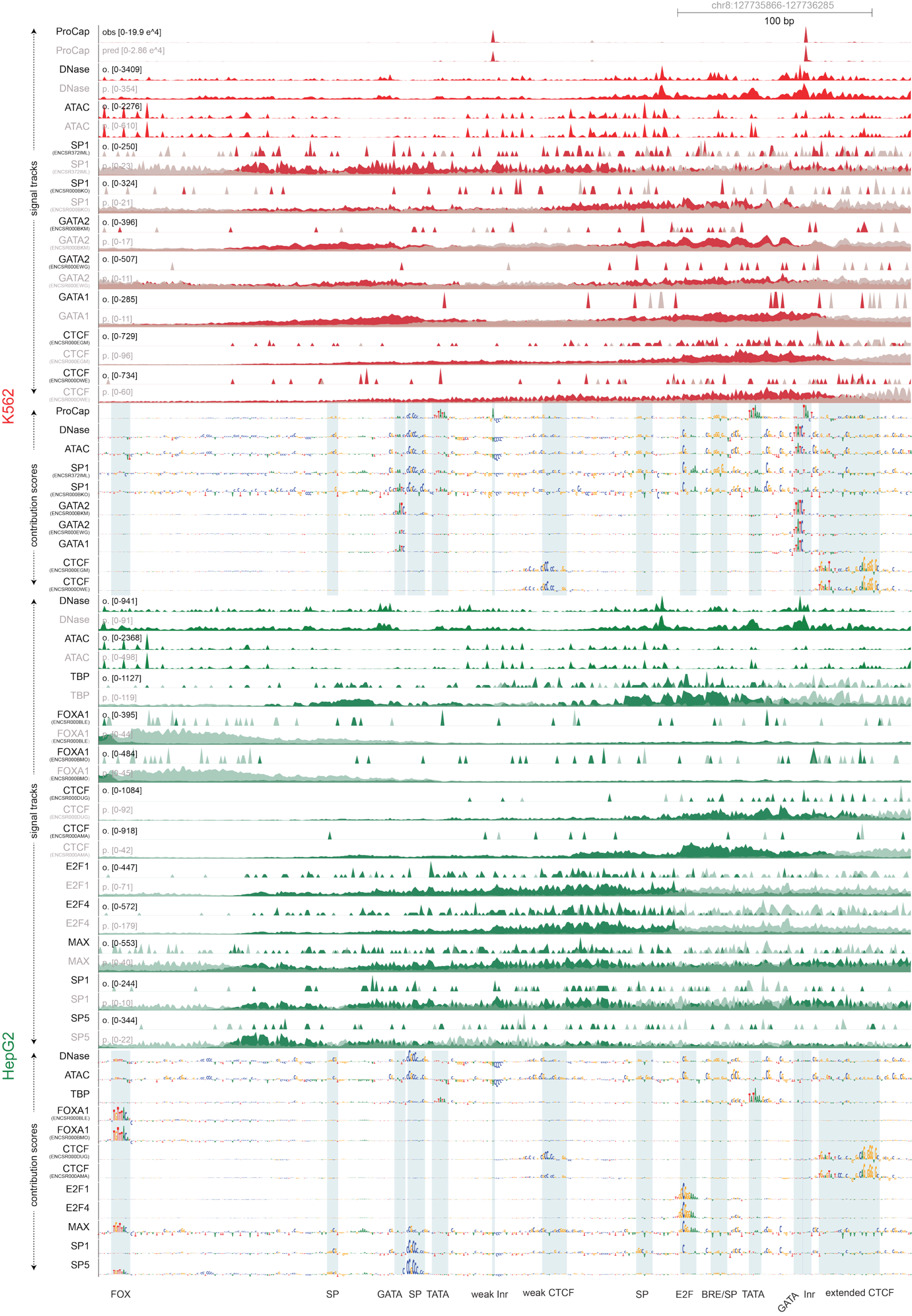

##### Supplementary Figure 17 | Model-derived profiles and contribution maps distinguish initiation, accessibility, and occupancy syntax at the MYC promoter

Tracks across the MYC promoter (chr8:127,735,867–127,736,285) comparing K562 and HepG2 showing observed (track labels in black) PRO-cap nascent transcription, DNase-seq/ATAC-seq accessibility, and TF ChIP-seq base-resolution profiles (fold-enrichment), alongside corresponding model predictions (track labels in light grey) and sequence contribution maps from ProCapNet (PRO-cap), ChromBPNet (DNase/ATAC), and BPNet (ChIP-seq). ChIP-seq tracks are shown stranded (positive/negative strand coverage in dark/light red). ProCapNet resolves two punctate TSSs in K562 with contribution maps highlighting canonical initiation syntax (e.g., Inr at the TSS with upstream TATA and SP/BRE elements). In contrast, ChromBPNet contribution maps emphasize a distinct promoter accessibility grammar dominated by SP/CpG-like features that is broadly shared between K562 and HepG2, with additional K562-specific GATA motifs near the stronger TSS. Despite variable observed ChIP-seq profiles, BPNet produces higher-resolution and more concordant predicted profiles and contribution maps across replicate/paralog experiments and highlights additional TF-specific sequence determinants (e.g., extended CTCF motifs influencing only CTCF binding) that are partially uncoupled from accessibility. Predictive motif instances are annotated beneath the contribution tracks.

#### Case Study 2 | Epistatic interactions between DNA sequences

To test whether the predictive sequence models capture non-additive (epistatic) motif syntax, we designed two MPRA libraries in K562: (i) a GATA–GATA pair library spanning diverse motif spacings and (ii) a motif–flank library probing interactions between a single GATA motif and its proximal flanks (**Supplementary Figure 18a**; **Supplementary Figure 19a**). We prioritized genomic loci whose model-derived contribution maps highlighted strong, predictive GATA instances, and assayed the corresponding wild-type sequences alongside systematic perturbations by MPRA. In the pair library, we combinatorially scrambled each GATA site individually or both sites together. In the motif–flank library, we scrambled the GATA motif, its flanking sequence, or both. We then compared MPRA-measured effects for all wild-type and perturbed constructs to in silico predictions from deep learning models trained in K562 on DNase-seq and ATAC-seq (ChromBPNet), GATA1/GATA2/TAL1 ChIP-seq (BPNet), and an independent variant MPRA dataset (ReporterNet) (**Supplementary Figure 18b–d**; **Supplementary Figure 19b**).

We quantified epistasis as the deviation of joint perturbation effects from additive effects of individual feature perturbations (**Supplementary Figure 18e-f**).^18-21^ Across both libraries, epistasis predicted by the DNase-based ChromBPNet model agreed well with MPRA measurements (pair library Pearson *r* = 0.55, **Supplementary Figure 18g**; motif–flank library *r* = 0.63, **Supplementary Figure 19c**). Models trained on ATAC-seq and MPRA showed similar concordance, with slightly lower agreement for TF ChIP-seq BPNet models (GATA1/2/TAL1) (**Supplementary Figure 18h**; **Supplementary Figure 19d**). Together, these results indicate that interaction rules and effect sizes learned from TF-occupancy and chromatin-accessibility models transfer to reporter activity.

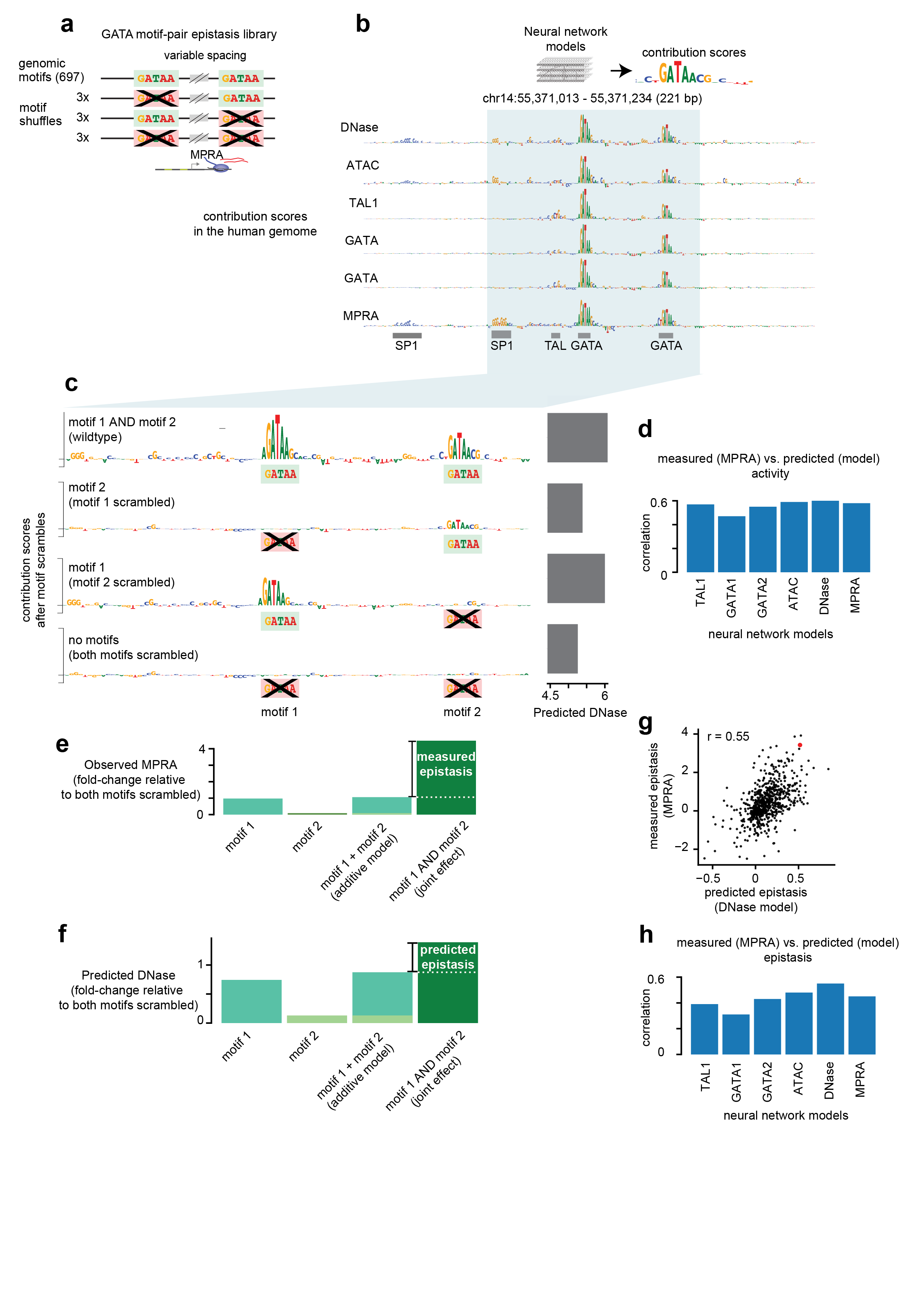

##### Supplementary Figure 18 | MPRA validation of GATA motif–pair epistasis predicted by predictive sequence models

**(a)** Design of the GATA motif–pair epistasis MPRA library. A total of 697 genomic loci containing two GATA motifs at variable spacings were selected, and each locus was assayed as the wild-type sequence together with combinatorial motif-scramble variants (motif 1 scrambled, motif 2 scrambled, and both motifs scrambled). **(b)** Example locus (chr14:55,371,013–55,371,234; 221 bp) illustrating model-derived base-resolution contribution scores across assays/models (ChromBPNet DNase/ATAC, BPNet models of TAL1/GATA1/GATA2, and a variant MPRA model), highlighting two GATA motifs and flanking TAL and SP1 motifs as key predictive sequence features. **c)** In silico motif-scramble analysis at the same locus. Contribution score tracks are shown for wild type (both motifs intact), each single scramble, and the double scramble, demonstrating that scrambling one GATA reduces predicted activity and also reduces the apparent contribution of the intact partner motif—consistent with non-additive interaction. The right-side bar summarizes the predicted DNase signal for each sequence. **(d)** Model performance for first-order activity effects: correlation between measured MPRA activity and predicted activity over the whole library for different models (TAL1, GATA1, GATA2 BPNet; ATAC- and DNase-ChromBPNet; MPRA model). **(e)** Definition of *measured epistasis* from MPRA data as the deviation from additivity: the joint effect of mutating both motifs compared to the sum of single-motif effects (effects are estimated as fold-change relative to the double-scramble baseline). **(f)** Analogous definition of *predicted epistasis* computed from the DNase-ChromBPNet model outputs using the same additivity framework. **(g)** Scatter plot comparing predicted epistasis (DNase-ChromBPNet) to measured epistasis (MPRA) across all tested loci, showing substantial agreement (Pearson *r* = 0.55). **(h)** Summary of epistasis prediction performance across models (correlation between MPRA-measured and model-predicted epistasis), showing that models trained on DNase/ATAC accessibility, TF ChIP-seq, and MPRA data all capture GATA–GATA interaction effects to varying degrees.

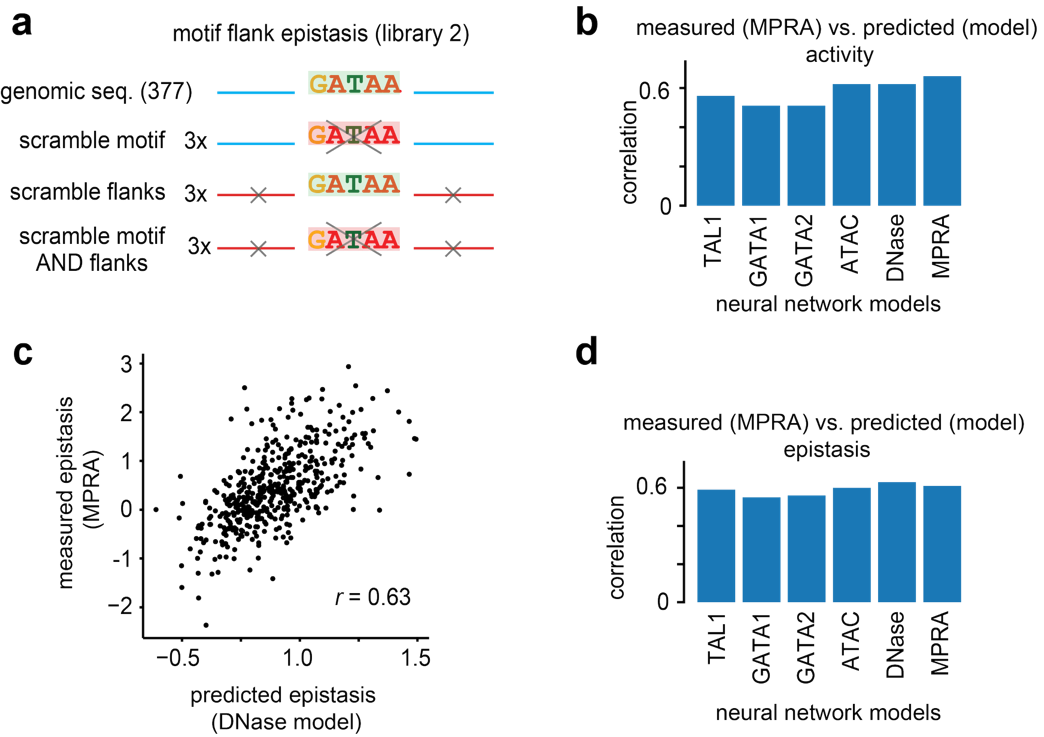

##### Supplementary Figure 19 | MPRA validates that predictive sequence models capture epistasis between GATA motifs and flanking sequence context

**(a)** Design of the motif–flank epistasis library: 377 genomic sequences containing a GATAA motif were assayed alongside matched perturbations in which the motif was scrambled, the flanking bases were scrambled, or both motif and flanks were scrambled (three independent scrambles per category). **(b)** Concordance between measured MPRA activity and model-predicted activity across models trained on TF ChIP-seq (TAL1, GATA1, GATA2), chromatin accessibility (ATAC, DNase), and variant MPRA data (ReporterNet). **(c)** Scatter plot comparing measured MPRA epistasis to predicted epistasis from the DNase-based ChromBPNet model (Pearson *r* = 0.63); epistasis is defined as deviation from additivity of joint versus single perturbations. **(d)** Model-wise correlations between measured and predicted epistasis, showing broadly consistent recovery of motif–flank interaction effects across assay-trained models.

#### Case Study 3 | Predicting noncoding variants causing common and rare disease

In this case study, we demonstrate the use of ENCODE-predicted variant effects to nominate gene regulatory mechanisms contributing to human disease at two loci.

The tyrosine-protein phosphatase non-receptor type 2 gene (PTPN2) is involved in a wide range of cellular processes. Mutations in PTPN2 are associated with human T-cell acute lymphoblastic leukemia (T-ALL). Noncoding genetic variation in the PTPN2 locus has also been associated with inflammatory bowel disease in genome-wide association studies (PMID:) and that result has been reproduced several times.^22,23^ The lead variant from genetic association is rs2542151. Using cV2F to fine-map the genetic association locus improved the posterior inclusion probability (PIP) for the intronic variant rs62097857 from 0.64 to 0.94 (**Supplementary Figure 20a**). That variant corresponds to predicted allele-specific transcription factor binding in enterocytes, goblet cells, and regulatory T cells from ENCODE machine learning models. Sequence motif analysis using the ENCODE motif catalog further predicts that the causal mechanism involves AP-1 binding.

Meanwhile, as an example application to rare disease, we assessed allelic effects of ultra-rare variants in short tandem repeats that were previously shown to cause rare monogenic thyrotropin resistance (**Supplementary Figure 20b**).^24^ Both variants showed significant allelic activity exclusively in normal and tumor bulk samples from thyroid cancer patients. ENCODE sequence contribution models predict that the variants impact a thyroid-specific enhancer via cooperative effects.

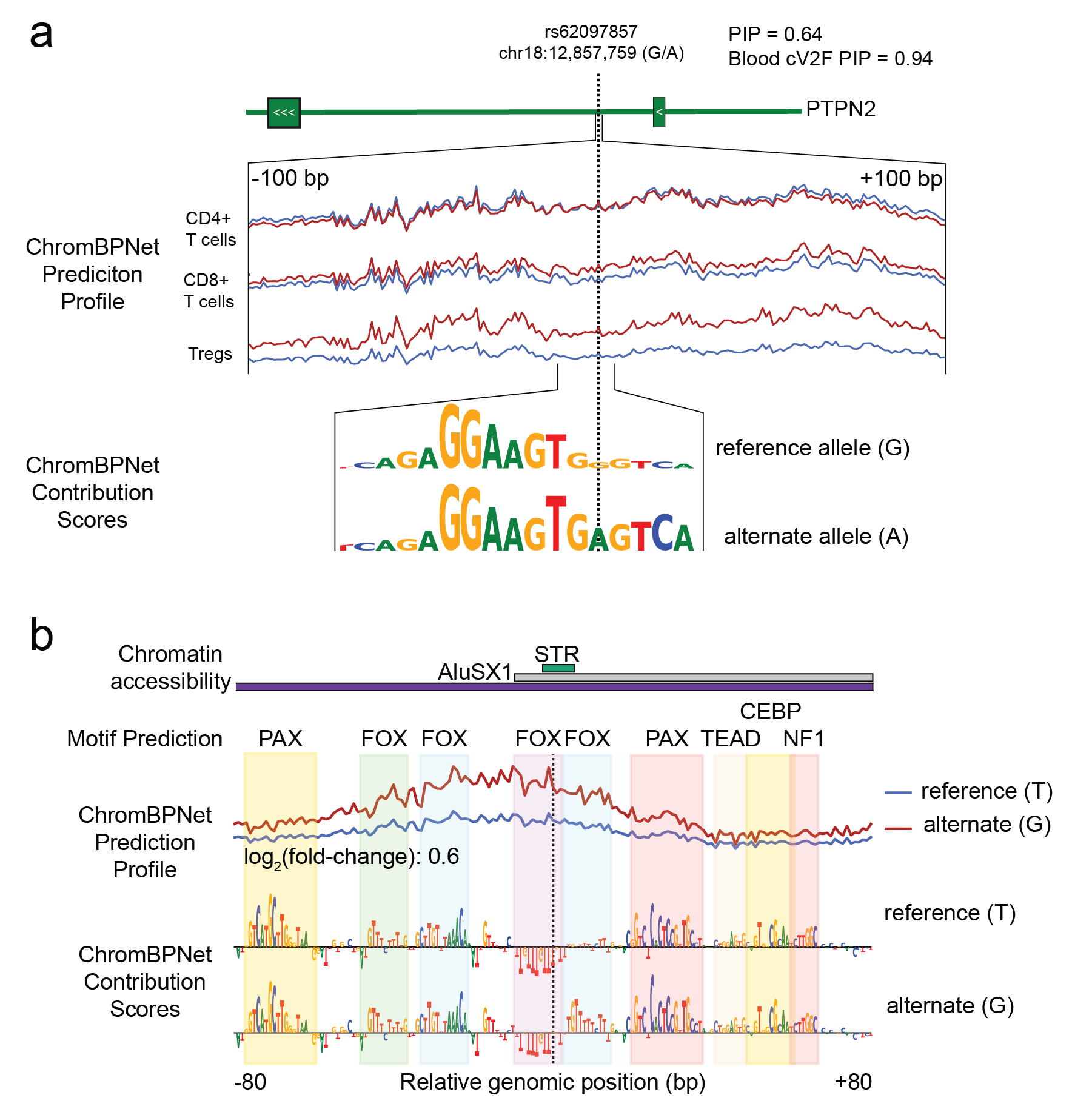

##### Supplementary Figure 20 | Fine mapping disease loci using ENCODE datasets

**(**a) cV2F scores nominate rs62097857 as a causal variant contributing to inflammatory bowel disease, improving the posterior inclusion probability (PIP) from 0.64 to 0.94. Machine learning models of the locus (ChromBPNet) predict that the alternate allele (red) has greater chromatin accessibility in Treg cells. That variant also corresponds to the predicted creation of an AP-1 transcription factor binding site in the locus. (b) An STR associated with rare monogenic thyrotropin resistance is predicted to cause increased chromatin accessibility. That effect corresponds to the predicted gain of a forkhead box transcription factor binding site (light blue).

#### Case Study 4 | Predicting noncoding variants causing common and rare disease

To illustrate how phased epigenetic (ChIP-Seq) and nucleomic (intact Hi-C) data can be combined with machine learning models and used to interpret the functional consequences of individual variants, we examined two loops in GM12878 lymphoblastoid cells. For each of these loops, a heterozygous SNP falls within a ChromBPNet predicted transcription factor binding motif at the loop anchor. The phasing is derived from intact Hi-C data, which was used to generate chromosome-length haploblocks.

We compared haplotype-resolved contact strength at the loop pixel by constructing homolog-specific 1 kb contact submatrices centered on the loop pixel and highlighting observed contacts within the central 5 × 5 bins to estimate per-homolog contact support.

In the first example (Supplementary Figure 21a), an AP-1–associated loop on chr4 contains a heterozygous T>C variant at chr4:104,719,749 within an AP-1 motif overlapping BATF and JUNB ChIP-seq signal. ChromBPNet CWMs for the motif present at this site indicate that the alternate allele disrupts the first position of the AP-1 core motif. Consistent with this prediction, the homolog carrying the motif-disrupting allele showed reduced contact frequency at the loop pixel, with 75 contacts on the unaffected homolog compared with 14 on the other, corresponding to a 5.3-fold allelic imbalance.

In the second example, a CTCF-associated loop on chr10 contains a heterozygous C>G variant at chr10:121,887,192, within a CTCF motif. According to ChromBPNet, the variant disrupts a high-information position in the CTCF core motif. The loop pixel again showed strong haplotype-specific contact imbalance, with 100 contacts on the unaffected homolog compared with 7 on the other, corresponding to a 14.2-fold difference (Supplementary Figure 21b). In both examples, the homolog carrying the motif-disrupting allele showed reduced contact frequency.

Together, these examples show how ENCODE resources can link individual sequence variants to allele-specific molecular phenotypes across regulatory readouts, including transcription factor ChIP-seq signal, sequence-model motif predictions, and chromatin looping.

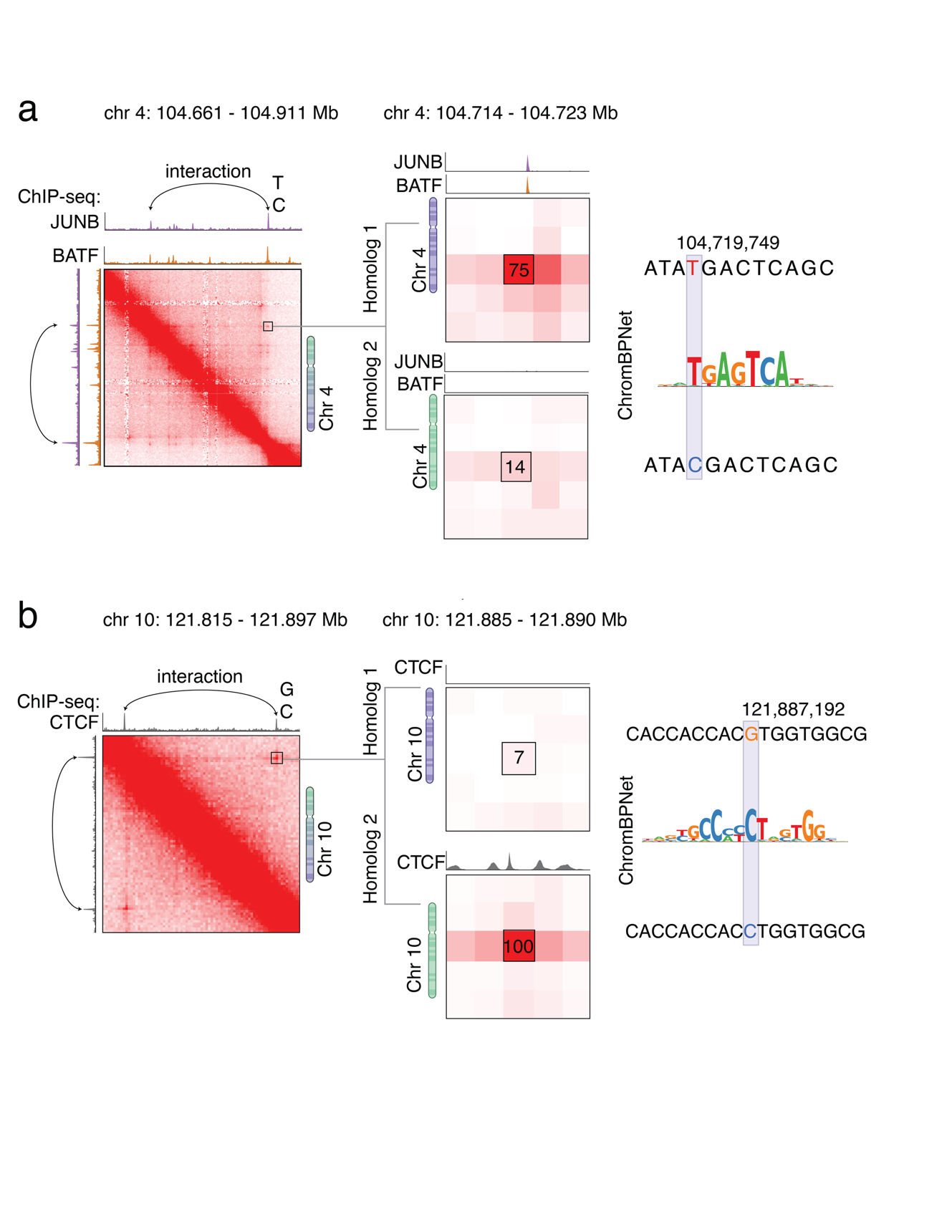

##### Supplementary Figure 21 | Disruption of binding sites drives allele-specific looping

**a,** On chromosome 4, we identified allele specific physical interactions corresponding to an AP-1 motif sequence. A T>C variant on chr4:104,719,749 disrupts an AP-1 motif identified by ChromBPNet and leads to a ~5.3-fold drop in loop contacts on the disrupted-motif homolog. **b**, On chromosome 10, we identified allele specific physical interactions that correspond to a CTCF motif sequence. A C>G variant on chr10:121,887,192 disrupts a CTCF motif identified by ChromBPNet and leads to a ~14-fold drop in loop contacts on the disrupted-motif homolog.

#### Case Study 5 | Dynamics of gene regulation across human crypt cell development

Integrating across the ENCODE datasets enables mapping regulatory dynamics of cell maturation in human tissues. For example, within the colon, intestinal stem cells continuously renew the epithelial lining of the intestine by differentiating into mature enterocytes, as well as specialized cell types including goblet, tuft, and enteroendocrine cells (**Supplementary Figure 22a**). To demonstrate how ENCODE datasets enable examination of regulatory states across differentiation, we defined a differentiation trajectory along the maturation pathway of absorptive epithelial cells (Fig. 1a). The expression of 38 TFs were correlated with motif activity of the same TFs along this trajectory (r > 0.6) (**Supplementary Figure 22b**), suggesting that these TFs may be regulators of intestinal differentiation. This analysis nominated TFs involved in intestinal stem cell maintenance, such as SOX9^25^ and ASCL2^26^, and regulators of differentiated enterocytes, such as HNF4A. We examined chromatin accessibility around the locus of one of these TFs, ASCL2, and observed several regulatory elements that become less accessible in more mature absorptive intestinal cells (**Supplementary Figure 22c**). Several of the regulatory elements around ASCL2 were also nominated as regulators of gene expression in ENCODE-rE2G models (**Supplementary Figure 22c**). Finally, we used deep learning sequence models of cell-type resolved chromatin accessibility profiles to annotate predictive TF motif instances in all accessible regions. For example, an accessible regulatory element near the ASCL2 gene showed predictive motif instances of SOX, FOX, and HNF1B TFs (**Supplementary Figure 22d**), which were also found in the initial correlative analysis as potential regulators of the immature or stem cell state (**Supplementary Figure 22b**). This provides an example of where these TFs may be binding to regulate the expression of ASCL2.

To define a trajectory of cell differentiation from colon stem cells to enterocytes, we started with data from the two multiome datasets where RNA-seq and ATAC-seq data was available from the same nuclei. We defined an LSI dimensionality reduction on the ATAC-seq data from these two samples using the ArchR^27^ function addIterativeLSI with 3 iterations and 20,000 variable features. We corrected batch effects using the ArchR function addHarmony^28^. We defined clusters from the harmony dimensions using the function addClusters and called peaks using the functions addGroupCoverages and addReproduciblePeakSet. We next defined a differentiation trajectory using addTrajectory with trajectory set to three large clusters of absorptive epithelial cells defined in the previous step. We defined motif activity scores using the functions addMotifAnnotations with motifSet = "cisbp" and addDeviationsMatrix. We then used the function getTrajectory to generate matrices of gene expression and motif activity along the differentiation pseudotime. We identified transcription factors whose expression was correlated with their motif activity along the differentiation trajectory using the function correlateTrajectories with varCutOff1 = 0.6, varCutOff2 = 0.6, and corCutOff = 0.6.

To plot accessibility tracks, we first exported group coverage BigWigs for each cell type using the function getGroupBW with normMethod = "ReadsInTSS", tileSize = 100, maxCells = 20000, and ceiling = 4. These group coverage BigWigs were loaded into the IGV genome browser for plotting. Thresholded bedpe files from ENCODE-rE2G models ([Dataset 1](https://www.encodeproject.org/files/ENCFF577BJW), [Dataset 2](https://www.encodeproject.org/files/ENCFF550BLY)) were used to plot predicted enhancer-gene links. Dot plots of ASCL2 expression were generated with the Seurat function DotPlot. For plotting results of the BPNet models, we plotted the results from the following pseudobulks in the UCSC genome browser: snATAC pseudobulk for human colon stem cell in adult/child and snATAC pseudobulk for human colon enterocyte in adult/child. We selected the pseudobulks from all samples to maximize coverage for model generation.

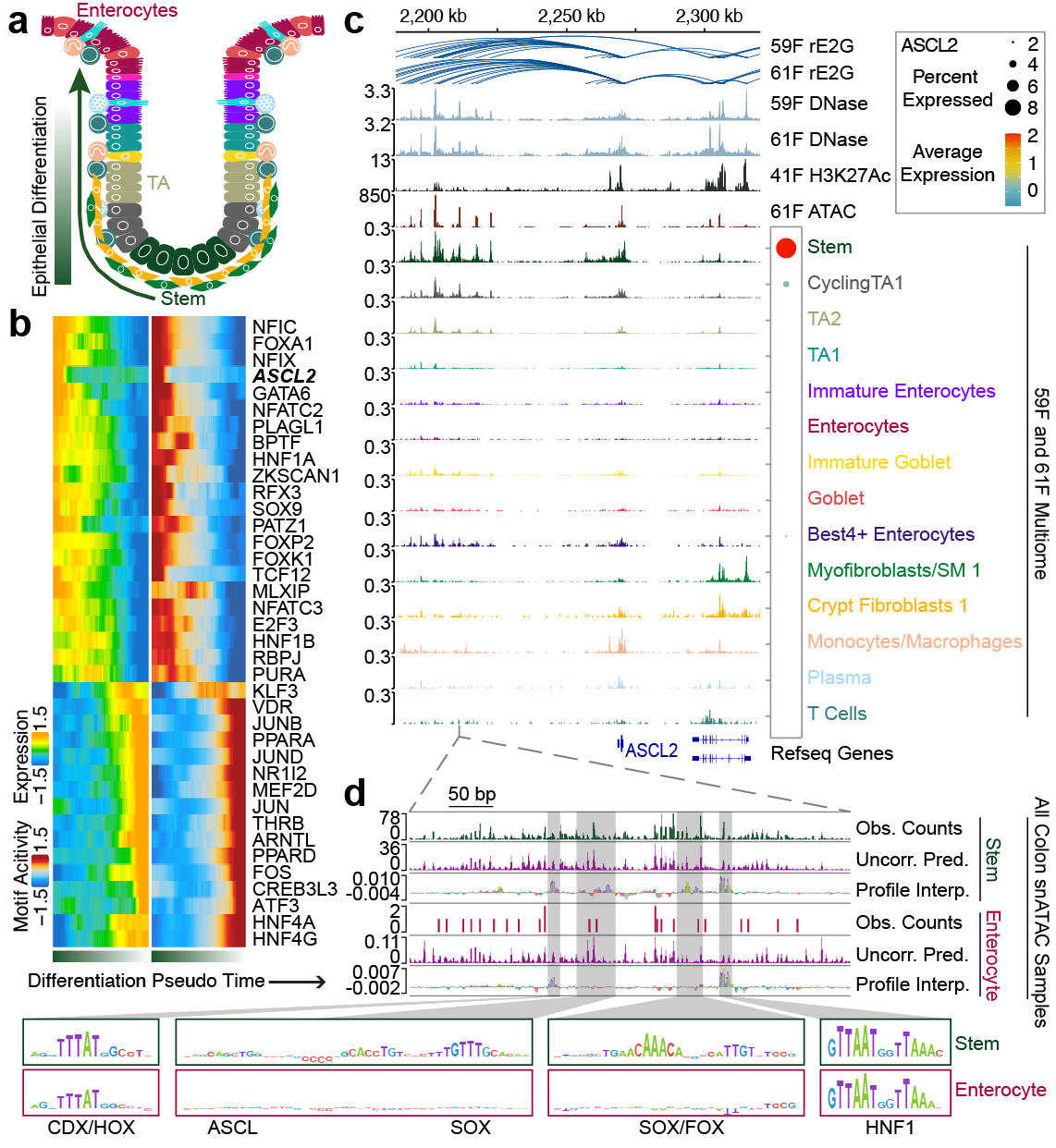

##### Supplementary Figure 22 | Using ENCODE data to dissect regulatory mechanisms during intestinal crypt cell development

(a) Schematic of epithelial differentiation in the colonic crypts. Cells are colored by cell type as indicated in (c). Not all cell types are shown. (b) TFs with correlation (r > 0.6) between their RNA expression and activity of the DNA motif to which they bind. RNA expression is plotted on the left and motif activity is plotted on the right. (c) Accessibility tracks around the ASCL2 locus for data from two bulk DNase donors, one bulk H3K27ac donor, one bulk ATAC-seq donor, and selected cell-types form the two snATAC multiome datasets. The expression of ASCL2 in different cell types in the multiome datasets is depicted in the dotplot on the right. Enhancer-gene links predicted by the ENCODE-rE2G models for two donors are depicted above the accessibility tracks. (d) Accessibility around ASCL2 at a single enhancer (chr11:2,211,013 - 2,211,514) for Stem cells and Enterocytes. For each cell type, the observed base-resolution scATAC-seq read count profiles, the predicted profiles from the BPNet, and the corresponding single-base pair contribution score profiles are shown. Contribution scores for selected regions are further zoomed in on at the bottom.

#### Case Study 6 | Changes in cCRE usage during transdifferentiation

To investigate chromatin dynamics at the cCREs during cellular differentiation, we used ChIP-seq data on nine histone modifications at twelve time points during the induced differentiation of human B-cell precursors into macrophages (**Figure 23a**, [Datasets](https://www.encodeproject.org/search/?type=Experiment&searchTerm=blaer)). We focused on a set of 110,544 accessible cCREs (ATAC+ cCREs), leveraging ATAC-seq data produced for this system.^29^ We employed a temporal Hidden Markov Model (tHMM) ^30^ to identify the major chromatin states—i.e., combinations of histone modifications—in which the cCREs can be found during transdifferentiation, and the transitions between these states during the process.

We found that six states adequately describe the epigenetic status of these 110,544 cCREs (**Supplementary Figure 23b; Supplementary Figure 23c**). Five of these states represent incremental deposition of histone marks, and we named them: Absent (A), with no marking, Basal (B), with low H3K4me1/me2 marking, Intermediate (I), with high H3K4me1/me2 marking and weak H3K27ac marking, Prominent (P), with high marking of H3K4me1/me2 and H3K27ac, and weak H3K9ac marking, Strong (S), with strong signal of all activation-associated marks, including H3K4me3, but H3K4me1. A sixth state, orthogonal to all these, is characterized by both high H3K4me2 and high H3K27me3, and we thus named it **Bivalent (Bv)**. State assignment varied greatly depending on cCRE classification: promoter-like or proximal enhancer-like (PLSs and pELSs) cCREs were enriched in H3K4me3-marked tHMM states, while these were infrequent in both distal enhancer-like (dELSs) and open chromatin (CA) cCREs, the latter enriched in Absent states compared to the former (**Supplementary Figure 23d**).

Overall, 50% of the cCREs remained in the same state during transdifferentiation and more than 86% were in no more than two states. However, these proportions depended very much on the cCRE classification. PLSs and pELSs were the most stable: 83% and 75%, respectively, remained in the same state and 98% and 95%, respectively, were in no more than two states (**Supplementary Figure 24a**). In any case, transitions between states showed a general balance between marking gain and loss, reflected in a similar proportion of cCREs assigned to each state at every time point (**Supplementary Figures 24c, 24b**). Dramatic transitions from unmarked (Absent) to strongly marked states (Prominent or Strong), occur gradually, and they can seldom be seen across consecutive time points (**Supplementary Figures 24d, 24c**).

The general pattern observed during transdifferentiation can be illustrated in the case of the β-catenin region. Consistent with the upregulation of the gene during transdifferentiation, we observed transitions from low to high chromatin marking for most analyzed cCREs (**Supplementary Figure 24d**).

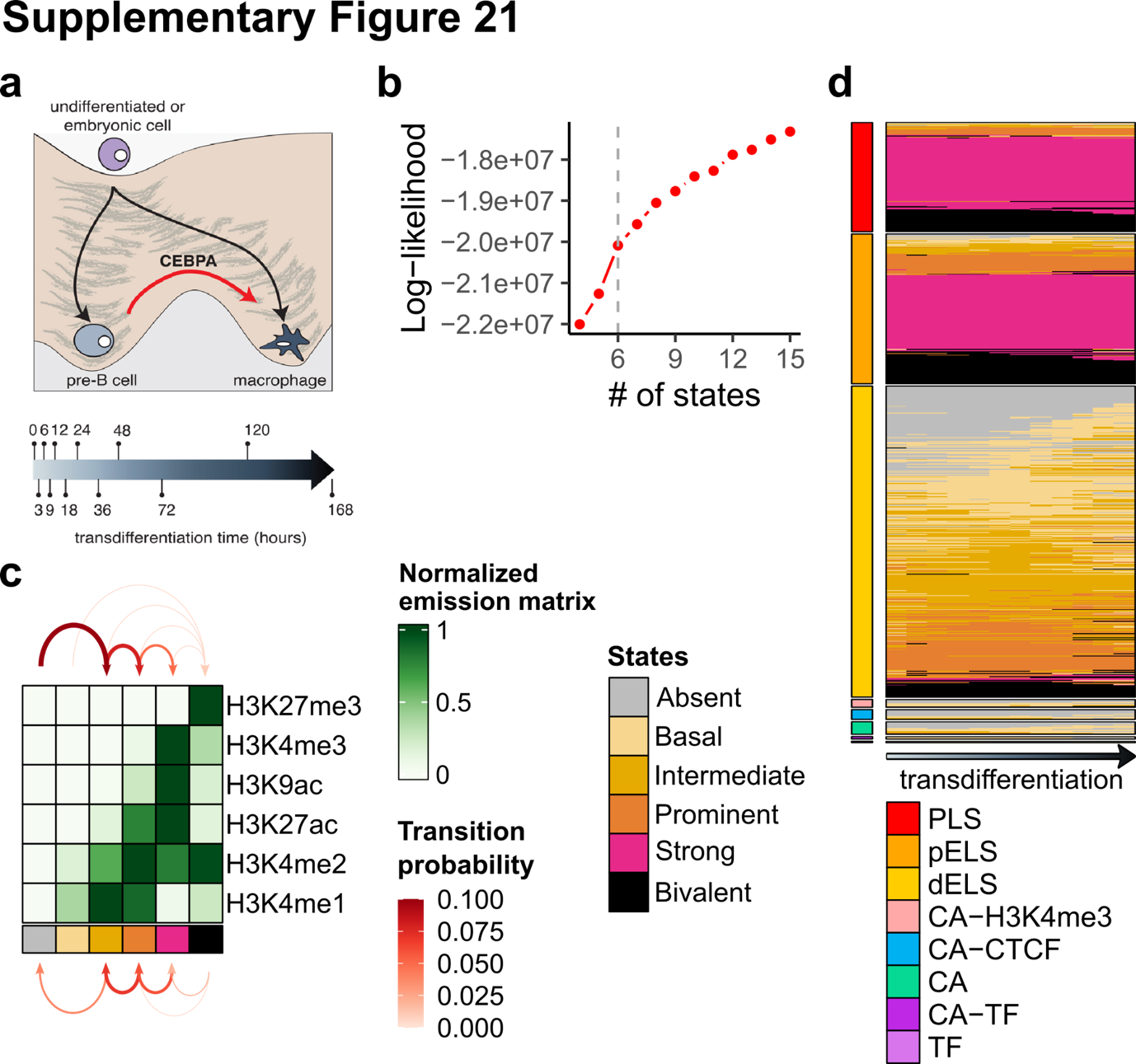

##### Supplementary Figure 23 | Recapitulating chromatin dynamics at genome and gene levels

**a,** Transdifferentiation of human B-cell precursors into macrophages induced by CEBPA expression. RNA-seq and ChIP-seq data on nine histone post-translational modifications were obtained at twelve time points during the process. Data available through ENCODE portal. **b,** Log likelihood values for tHMM models computed with between 4 and 15 states; The selected 6 states are indicated with a vertical gray dashed line. **c,** Chromatin states described as combinations of histone marks’ relative signal; arrows represent transitions between states for the same cCRE in consecutive time points, width proportional to absolute number of switching cCREs. **d,** Dynamics of chromatin states over transdifferentiation, each row corresponds to one cCRE (legend in panel **c**), and each column to a time point. ENCODE4’s classification is included to the left of the heatmap.

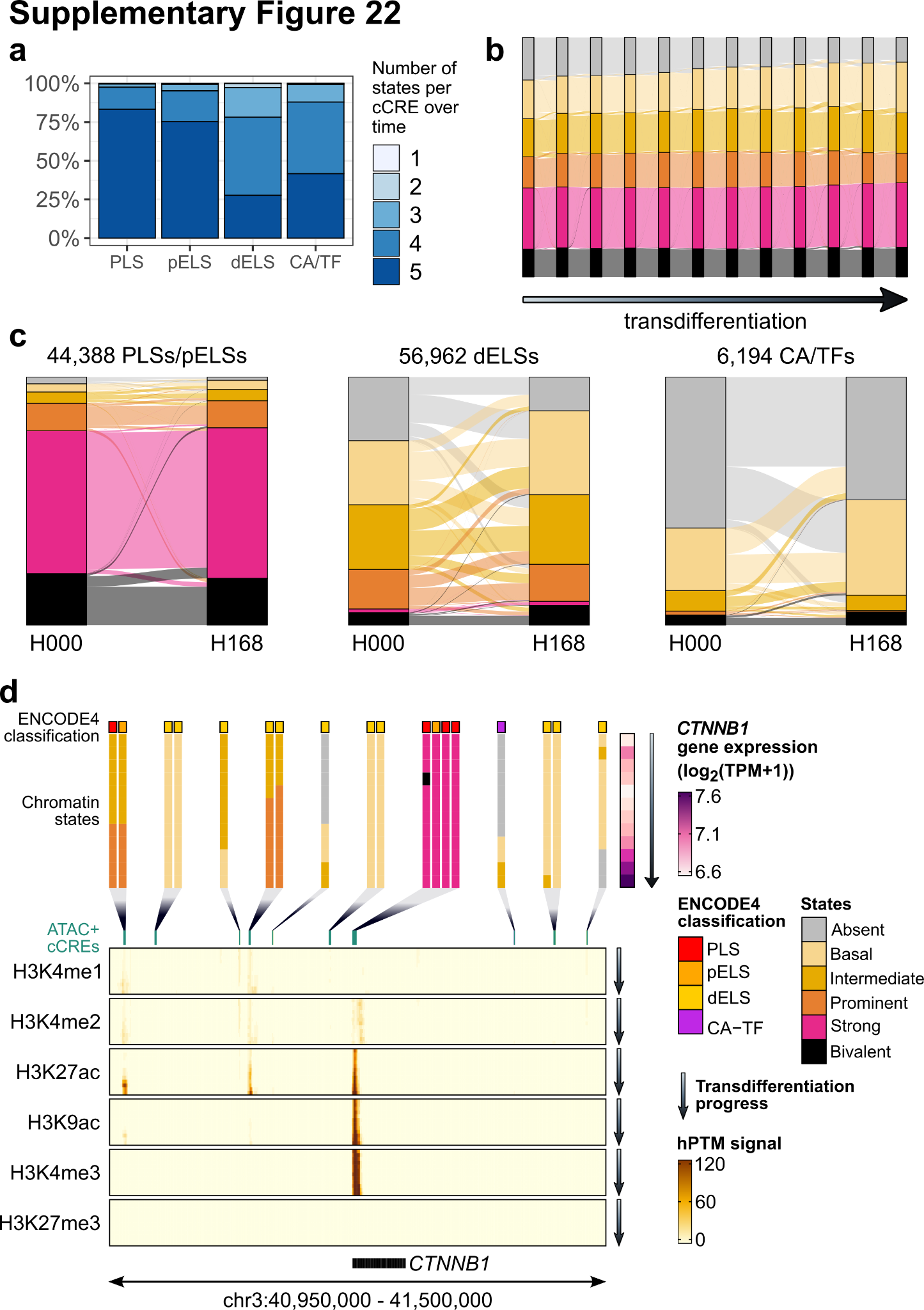

##### Supplementary Figure 24 | TSS proximity conditions chromatin signatures and their dynamics

**a,** Proportion of cCREs in a different number of states during transdifferentiation, depending on the ENCODE 4 cCRE category. CA/TF refers to a category comprising all cCREs that are not defined as PLSs, pELSs or dELSs. **b,** Sankey plot representing transitions between states in consecutive time points during transdifferentiation (time course indicated by the horizontal arrow, legend in **Supplementary Figure 23c**), as well as the proportion of cCREs assigned to each state during transdifferentiation. **c,** Sankey plots representing state transitions between first (0 h post-induction) and final (168 h post-induction) time points dividing cCREs into three categories i) TSS-proximal PLSs and pELSs, ii) marked TSS-distal dELSs, and iii) unmarked open chromatin regions/TF binding motifs (CA-H3K4me3’s, CA-CTCFs, CA-TFs, CAs, and TFs); the number of elements belonging to each category is indicated below each panel. **d,** Expression and chromatin marking at the E β-catenin locus. The expression of the gene (its genomic location indicated by a black rectangle at the bottom of the figure) along transdifferentiaion is indicated by a vertical bar at the top right on the figure. ChIP-seq for several histone modifications along transdifferentiation are shown in the body of the figure. ATAC+ cCREs are displayed and expanded into vertical bars at the top of the figure displaying their chromatin states along transdifferentiation. The box at the top of these bars indicated the ENCODE cCRE classification.

#### Case Study 7 | Stem cell differentiation toward the kidney lineage

The ENCODE stem cell differentiation resource provides a broad framework for interrogating epigenomic and chromatin-state remodeling as human pluripotent stem cells transition into multiple lineage derivatives (**Supplementary Figure 25a**, [Datasets](https://www.encodeproject.org/stem-cell-matrix/?type=Experiment&replicates.library.biosample.donor.accession=ENCDO222AAA&status=released&control_type!=*)). Across differentiated cell states representing ectodermal, mesodermal, and endodermal trajectories, these datasets enable the analysis of dynamic changes in histone-modification profiles and related regulatory features that accompany lineage commitment and progressive cell-state specification. This makes it possible to examine how regulatory programs are established, reinforced, or repressed during human differentiation in a comparative manner across organ systems. As one illustrative example, we focused on nephron lineage progression to assess how chromatin-state dynamics evolve as pluripotent stem cells acquire progressively more specialized kidney identities.

Human nephron organoids were generated from H9 human embryonic stem cells using a directed differentiation protocol that proceeds through a metanephric mesenchyme/nephron progenitor stage by day 8, followed by organoid formation and progressive maturation through day 21, day 35, and day 49. To define the epigenomic programs associated with nephron specification and temporal organoid maturation, ENCODE profiled these samples by Mint-ChIP-seq at the H9 embryonic stem cell stage (day 0), day 8 nephron progenitor cells, and nephron organoids collected at days 21, 35, and 49. Across these stages, Mint-ChIP-seq datasets were generated for H3K4me1, H3K4me3, H3K36me3, H3K9me3, H3K27me3, and H3K27ac, enabling assessment of promoter-, enhancer-, transcription-, and repression-associated chromatin-state transitions during differentiation. In keeping with ENCODE experimental standards, these datasets were generated with two biological replicates at each stage, providing a structured framework for comparing chromatin-state dynamics across sequential steps of human nephron differentiation and maturation.

Genome-wide chromatin-state profiles from this differentiation series are publicly accessible through the ENCODE portal, allowing users to query individual genes and visualize locus-specific histone modification patterns as a proxy for transcriptional regulatory status. In this nephron organoid model, directed differentiation induces nephron progenitor cells by days 8–9, followed by formation of early nephron structures by day 21 and progressive maturation through day 49.^31^ As an illustrative example, H3K4me3 Mint-ChIP-seq profiles at the SALL1 and AQP1 loci reveal distinct stage-specific promoter activation patterns (**Supplementary Figure 25b**). At the SALL1 promoter, H3K4me3 signal is weak in undifferentiated cells but becomes markedly broader and stronger at the nephron progenitor stage, consistent with transcriptional activation of a nephron progenitor program. This signal then progressively diminishes at day 21 and day 49, indicating that SALL1 activation is transient and associated with the progenitor state. In contrast, promoter-associated H3K4me3 at AQP1, a water channel and proximal tubular maturation marker, is weak in undifferentiated cells and nephron progenitor cells but becomes strongly enriched in late-stage organoids by day 49, consistent with progressive activation of a more mature tubular epithelial program.

As illustrated by this case study, the ENCODE portal provides a powerful framework for genome-wide interrogation of chromatin-state remodeling during human stem cell differentiation, enabling users to examine how gene activation and repression are dynamically established across developmental transitions using publicly accessible epigenomic datasets. In kidney organoids, which are often considered more similar to fetal than adult kidney states at earlier stages, longitudinal profiling nonetheless indicates that extended culture is accompanied by continued regulatory and transcriptional maturation rather than a static endpoint. These observations are consistent with prior studies documenting progressive maturation during kidney organoid differentiation,^32^ and highlight the value of this resource for assessing organoid maturity and for investigating mechanisms that promote further maturation. More broadly, ENCODE provides analogous differentiation datasets across multiple lineage trajectories derived from human pluripotent stem cells, making this platform a useful resource for investigators studying developmental gene regulation in a wide range of human cell and organoid models.

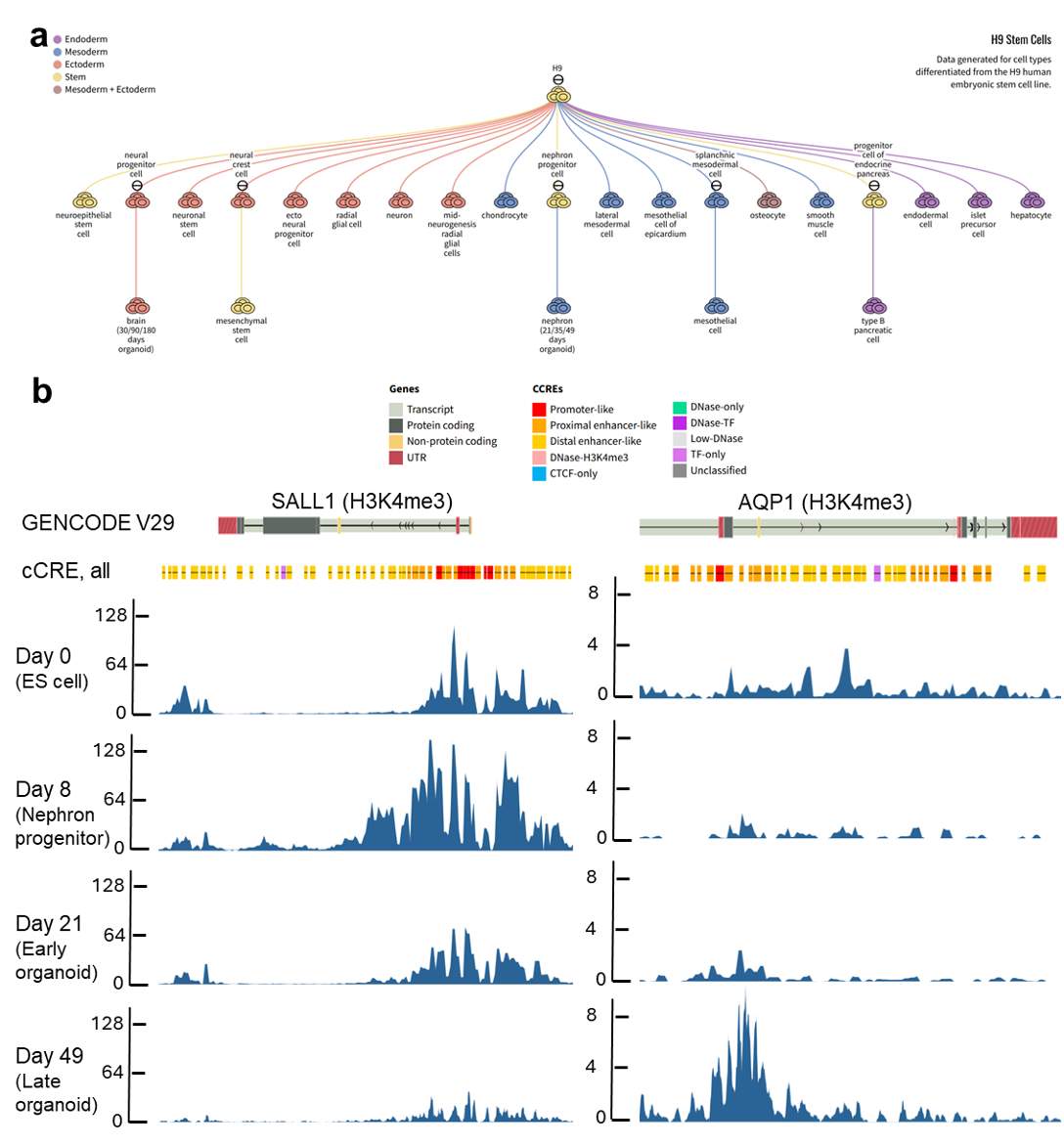

##### Supplementary Figure 25 | ENCODE data to explore longitudinal chromatin alterations during nephron organoid differentiation

**a,** A schematic summary of datasets available in ENCODE for multi-lineage differentiation from H9 human embryonic stem cells. **b,** Representative H3K4me3 Mint-ChIP-seq tracks at the SALL1 and AQP1 loci across nephron differentiation, including day 0 human embryonic stem cells, day 8 nephron progenitor cells, day 21 early nephron organoids, and day 49 late nephron organoids. GENCODE v29 gene annotations and ENCODE cCRE_all regulatory element annotations are shown above the signal tracks. These profiles illustrate stage-specific promoter-associated chromatin activation at a nephron progenitor marker (SALL1) and a proximal tubule maturation marker (AQP1).

#### Case Study 8 | Isoform switching during mouse development

Long read RNA sequencing reveals changes in transcript isoform usage during development. As one example, we identified multiple transcript isoforms for the class I fructose-bisphosphate aldolase gene *Aldoa*. *Aldoa* encodes a key glycolytic enzyme crucial for muscle function. Whereas the shorter isoform (*Aldoa[1,1,1]*) is expressed in all cell types, the longer isoform (*Aldoa[5,25,1]*) is preferred in myonuclei compared to other cell types in skeletal muscle (**Supplementary Figure 26a**). Comparing transcription start site usage in bulk muscle tissue across all mouse postnatal development reveals a switch from the shorter to the longer isoform between postnatal days 4 and 10 (**Supplementary Figure 26b**). Chromatin accessibility from pseudobulked single cell ATAC-seq highlights the specificity of the longer isoform in myonuclei compared to other cell types found in skeletal muscle, such as endothelial cells, fibro-adipogenic progenitor cells, and immune cells (**Supplementary Figure 26c**). This preference for *Aldoa[5,25,1]* in mature myonuclei may indicate an enhanced function of this isoform, given the high metabolic demand of developing myonuclei in comparison to other cell types.

Tissues at postnatal day (PND) 4, PND 10, PND 14, PND 25, PND 36, 2 months, and 18-20 months were collected from male and female C57BL6/J × CAST/EiJ F1 (“B6CASTF1/J”) hybrid mice. Bulk short- and long-read RNA-seq, microRNA-seq, single-nucleus RNA-seq, 10x Multiome (single-nucleus RNA-seq and single-nucleus ATAC-seq), DNase-seq, ChIP-seq, and Hi-C assays were performed on some or all tissues and timepoints. In addition to B6CASTF1/J, some assays were also performed on 5xFAD and 5xFAD/CAST F1 crosses at 8-14 months old. Five core tissues in B6CASTF1/J were chosen for comprehensive profiling by transcriptomic and single-cell assays: adrenal gland, left cerebral cortex, hippocampus, heart, and gastrocnemius.

A novel transcript annotation framework was then used to characterize the long-read RNA-seq data.^33^ Based on transcript annotations from the long-read RNA-seq analysis pipeline ([ENCPL239OZU](https://www.encodeproject.org/pipelines/ENCPL239OZU/)), the same triplet naming convention described in the main text was applied in mouse. Differential isoform expression testing between timepoints revealed a 5' isoform switch in skeletal muscle in *Aldoa* between Aldoa[1,1,1] and Aldoa[5, 35, 1].^34^ The relative abundance of each isoform among all the detected *Aldoa* transcripts, or percent isoform, was calculated as previously described.^34,35^ The accessibility at the TSS of each isoform was quantified using ArchR and pseudobulks
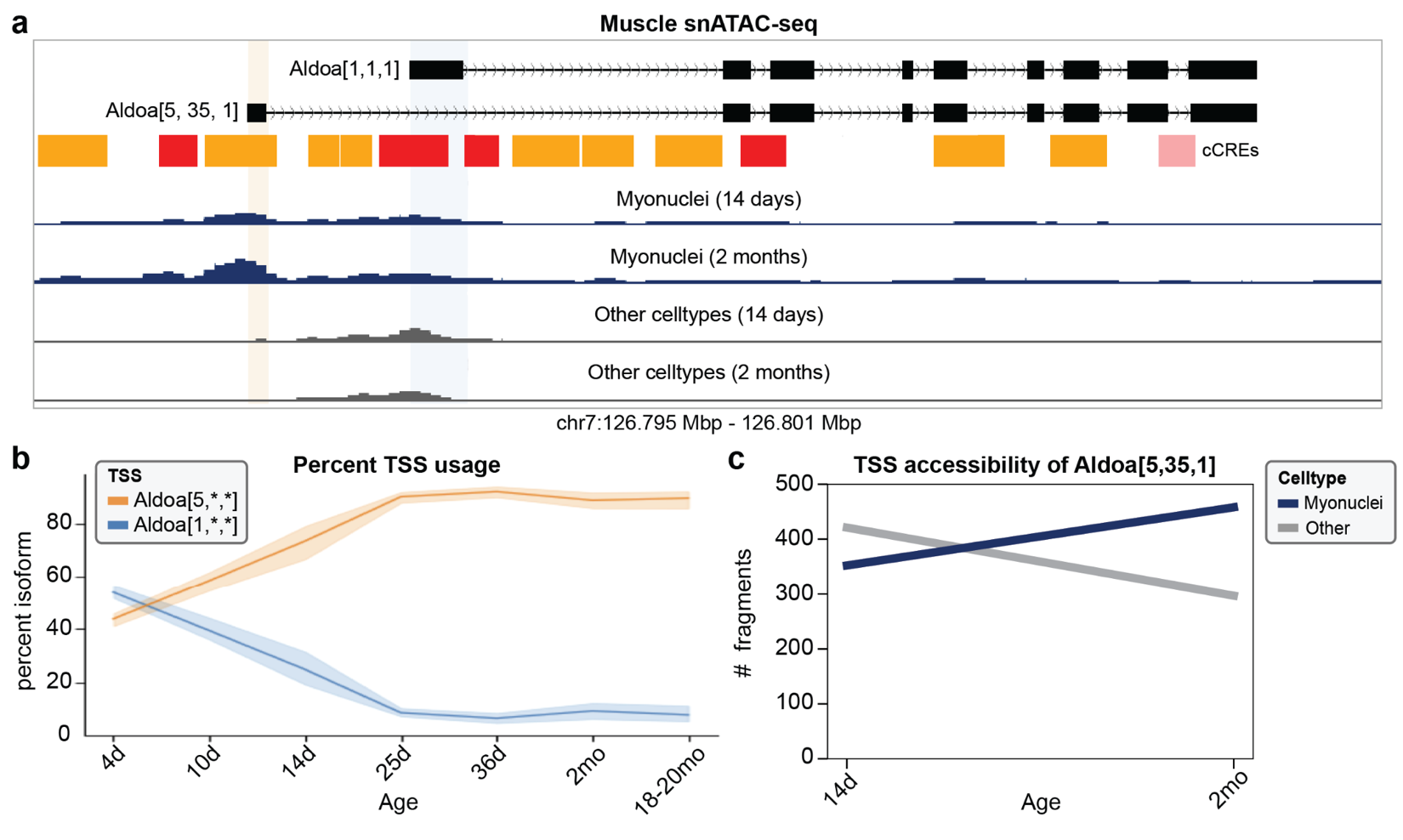
were calculated by sum across myonuclei and non-myonuclei cell types.

##### Supplementary Figure 26 | Changes in Aldoa isoform usage during mouse development

(a) Diagram of the short and long isoforms of Aldoa1 are shown at top. Below are representative RNA-seq signal tracks from myonuclei and other cell types collected from either 14 days or 2 months postnatal mice. (b) Percent of transcription start site usage in muscle over postnatal development for short (blue) and long (orange) isoform, as measured by RNA-seq. (c) Changes in chromatin accessibility at the promoter for the long isoform of Aldoa1 between 14-day-old and 2-month-old mice. Chromatin accessibility was measured using single-cell ATAC-seq. Blue shows accessibility in myonuclei and grey shows accessibility in other cell types.

### Supplementary Methods

#### Consensus Variant-to-Function Scoring and downstream applications

##### cV2F training and features

The consensus variant-to-function (cV2F) score is an integrative ensemble of 339 variant-level, element-level and baseline-LD (v2.2) features.^36^ The optimal combination of these features is generated by training a gradient boosting classification model on GWAS fine-mapped variants from 94 UK Biobank diseases and traits. We use for training data a set of positive variants that are confidently fine-mapped variants (PIP > 0.90) across 94 traits in UKBiobank as in ref. ^37^. We considered as negatives 1,227,661 variants that have very low fine-mapping posterior probability (PIP < 0.01) for all 94 UKBiobank traits. In order to ensure better matching between the positive and negative set of variants, we restrict our negative set of variants to the ones that are in the same LD block^38^ as a positive variant and up to 5 closest variants in terms of Minor Allele Frequency (MAF) as the positive variant. For tissue and cell-line level versions of cV2F, the feature sets were restricted to relevant tissues and cell types.

##### cV2F model

We used the Extreme gradient boosting (XGBoost) method implemented in the XGBoost software^39,40^ with the following model parameters: the number of estimators (25, 40, 50), depth of the tree (10, 15, 25), learning rate (0.05), gamma (minimum loss reduction required before additional partitioning on a leaf node; 10), minimum child weight (6, 8 ,10), and subsample (0.6, 0.8, 1); we optimized parameters by tuning hyper-parameters (a randomized search) with five-fold cross-validation. Two important parameters to avoid over-fitting are gamma and learning rate; we chose these values consistent with previous studies^77^, as in our previous work on AnnotBoost^41^ and DeepBoost^42^ frameworks.

The gradient boosting predictor is based on T additive estimators and it minimizes the loss objective function $L_{t}$ at iteration $t$.

$$L_{t}= \sum_{i=1..N} l \left( y_{i} , \hat{y}_{i} \left( t \right) \right)+ \gamma\left( f_{t} \right)\hat{y}_{i} \left( t \right)= \hat{y}_{i} \left( t-1 \right)+ f_{t}\left( x_{i} \right)$$

Where $f_{t}$ is an independent tree structure and $\gamma(f_{t})$ is the complexity parameter. The final prediction from the gradient boosting model therefore is given by

$$y_{i} = \sum_{t=1, 2, .., T} f_{t} (x_{i})$$

To avoid winner's curse and overfitting, we use fine-mapped SNPs on odd (respectively even) chromosomes as training data to make predictions for even (respectively odd) chromosomes, as in our previous work on AnnotBoost; thus, boosted annotations on a given chromosome are not informed by fine-mapped SNPs on that chromosome. We report the average AUPRC (Area under Precision Recall Curve) of odd and even chromosome classifiers. The boosted annotations produced as output of the classifier are probabilistic in nature because of the logistic loss. We have performed independent training and prediction generation for a model with all features (variant-level, element-level, baseline-LD), as well as different subsets of these features. The importance of each feature is assessed using Shapley scores^43^ and marginal AUPRC of each feature in the model.

##### cV2F-informed fine-mapping

We perform a functional fine-mapping of UKBiobank GWAS associations using the cV2F score following a similar approach as in Expression Modifier Score (EMS).^44^ The SuSIE fine-mapping model^45^ assumes a causal configuration vector $b = \sum_{l} b_{l}$where both $b$ and $b_{l}$ are vectors of length m, the number of variants in the locus to be fine-mapped, and $b_{l}$ has one entry equals to 1 and all other entries as 0. SuSIE defines a set of vectors (of length *m*) $\alpha_{1}, \alpha_{2}, ...., \alpha_{L}$ with $\alpha_{l}(v) = Pr ({|b}_{l}(v)| > 0 | X)$ is the posterior probability given the data $X$. Credible sets are computed for each 𝑙 from $\alpha_{l}$and credible sets that are not pure, meaning that contain a pair of variants with absolute correlation < 0.5, are pruned out. If $\alpha_{l}$ corresponds to a pure credible set, we re-weight each element of $\alpha_{l}$ with the binarized primary or tissue-specific score. Let $c_{1}, c_{2}, ...., c_{m}$be the cV2F scores for $m$ variants, then we generate updated posterior probabilities $\delta_{l}$ as

$$\delta_{l}(v) = c_{v}\alpha_{l}(v) / \sum_{u=1(i)m} c_{u}\alpha_{l}(u)$$

We use these updated cV2F-informed posterior probability $\delta_{1}, \delta_{2}, ...., \delta_{L}$to compute updated credible sets and posterior inclusion probabilities (cV2F.PIP). We compare these cV2F.PIP values with the standard PIP values obtained from the non-functionally-informed posterior probabilities $\alpha_{l}(v)$. For the tissue-specific approach, we replaced the cV2F score with tissue or cell-line cV2F score. For the analysis in the paper, we focused on blood and liver since several related biomarker traits with well-powered fine-mapping were available for those two tissues.

##### cV2F-informed polygenic risk score model

To generate cV2F-informed polygenic risk scores (PRS), we applied Priors-informed Regression for Inclusive Score Modeling (PRISM), a transfer-learning based framework that allows ancestry-aware integration of tissue-specific annotations for improved PGS transferability.^46,47^ We implemented PRISM by extending the iPGS approach, a penalized regression directly learned on the individual-level data that minimizes the following loss function:

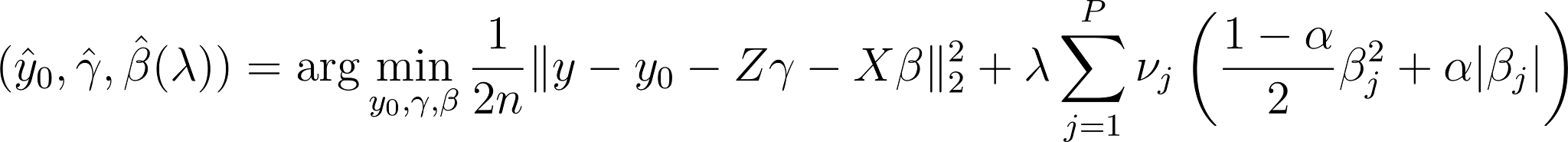
<https://www.codecogs.com/eqnedit.php?latex=(%5Chat%7By%7D_0%2C%5Chat%7B%5Cgamma%7D%2C%5Chat%7B%5Cbeta%7D(%5Clambda))%3D%5Carg%5Cmin_%7By_0%2C%5Cgamma%2C%5Cbeta%7D%5Cfrac%7B1%7D%7B2n%7D%5C%7Cy-y_0-Z%5Cgamma-X%5Cbeta%5C%7C_2%5E2%2B%5Clambda%5Csum_%7Bj%3D1%7D%5E%7BP%7D%5Cnu_j%5Cleft(%5Cfrac%7B1-%5Calpha%7D%7B2%7D%5Cbeta_j%5E2%2B%5Calpha%7C%5Cbeta_j%7C%5Cright)#0>

where $y$ is the phenotype, $Z$ is the covariates, and $X$ is the genetic predictors. Here, $\hat{y_{0}}$ is the estimated intercept of the regression model, and $\hat{\gamma}$, and $\hat{\beta}$ are the estimated coefficients for covariates and genetic predictors, respectively. We set the elastic net parameter $\alpha$ to 0.99. We grouped variants based on their predicted consequence and cV2F scores and assigned a penalty factor $\nu_{j}$ to each group. The detailed grouping scheme and penalty factor assignments are described in the companion PRISM manuscript.

We trained PRISM models using individual-level genetic data from 406,659 ancestrally diverse individuals in the UK Biobank (UKB),^48^ adjusting for age, sex, Townsend deprivation index, and the top 18 genotype principal components. We used cV2F scores that leverage fine-mapping results from African ancestry individuals in the Million Veteran Program to avoid data leakage.^49,50^ The estimated variant effect sizes from the PRISM models were then used to construct PRS for downstream evaluation.

We constructed tissue-matched and tissue-non-specific PRS given the type of cV2F score used. Tissue-matched PRS were derived from PRISM models using cV2F scores trained with annotations from samples corresponding to the trait’s primary tissue, whereas tissue-non-specific PRS were derived from PRISM models using cV2F scores trained with all available annotations.

To assess whether cV2F-informed PRS improved predictive performance, we constructed a baseline PRS in which cV2F scores were not used to inform penalty factor assignment during model training. We evaluated the predictive performance (*R^2^*) of each PRS in a held-out set of African ancestry individuals in UKB (n=1,154 for lymphocyte count, n=1,134 for eGFR, n=1,078 for FEV_1_/FVC ratio, and n=1,130 for LDL-C). We compared the R^2^ of tissue-matched and tissue-non-specific PRS to the baseline PRS and reported the percentage improvement in R^2^.

#### Predictive DNA sequence models and unified motif interpretation

##### Data preprocessing

Starting from uniformly processed ENCODE 4 Homo sapiens TF ChIP-seq, DNase-seq and ATAC-seq datasets, base-resolution signal tracks (available as observed signal profile) were generated from the 5′ ends of mapped reads following the removal of PCR duplicates and low-quality reads.

We first identified datasets as sequenced in either a paired- or a single-ended manner, and downloaded the corresponding “alignments” or “unfiltered alignments” BAM files, respectively. We filtered the “unfiltered alignments” files for low quality reads with samtools (-F 780 and -q 30). We merged all BAM files for a single experiment and rejected experiments that consisted of a mix of single- and paired-ended alignments.

For DNase-seq and ATAC-seq data, we followed the additional preprocessing steps described in ref. ^19^. We next prepared base-resolution signal tracks with the appropriate positive and negative strand shifts for each assay type. We defined the set of peak regions for DNase data by calling peaks with macs2, filtering to peaks with an FDR-adjusted p-value < 0.05, and then merging with the set of ENCODE cCREs matched to that dataset, if available. For ChIP-seq and ATAC-seq data, we downloaded the default peak set from the ENCODE portal.

##### Model training

For the BPNet models, we adapted a previously-described BPNet architecture and training procedure ([GitHub Repository](https://github.com/kundajelab/bpnet)).^18^ The model accepts a one-hot-encoded 2,114 bp input sequence and predicts strand-specific ChIP-seq signal profiles and total log counts over a central 1,000 bp window. The model predicts the residual signal relative to the control input tracks. The architecture comprises nine dilated convolutional layers with residual connections. Profile predictions are generated via an additional convolutional layer applied to the final dilated output, integrating the control signal profiles. Total counts are predicted by applying global average pooling, followed by a dense layer incorporating control total counts. The model is trained using a joint loss function that combines a single negative log-likelihood loss for the profile logits across both the strands and mean squared error for total counts.

Training data consisted of IDR-thresholded peaks and GC-matched non-peak regions in a 3:1 ratio. Outlier peaks with total signal exceeding 1.2x the total signal of the 99th percentile peak were excluded. Data augmentation included random jittering (up to 128 bp) and reverse-complement transformations. Models were trained using chromosome-based 5-fold cross-validation with no overlap between training, validation, and test sets.

We trained DNase and ATAC ChromBPNet models using the procedure described in Pampari et al. ChromBPNet models consist of a similar architecture to BPNet, with additional filters in each convolutional layer.^19^ Correcting for DNase and ATAC bias requires training a BPNet-like bias model on a set of background regions with a GC-content matched to the peak regions. These bias models learn the intrinsic sequence preference biases of the DNase I (used in DNase-seq) and Tn5 (used in ATAC-seq) enzymes. The bias models are used to predict enzyme bias profiles in peak regions. The predicted bias profiles are regressed out from the observed DNase/ATAC-seq profiles and a second BPNet-like neural network is used to fit a sequence model to the residual profiles and counts after regressing out bias. After training, the residual network, which predicts bias-corrected profiles and counts is used for all downstream prediction, interpretation and motif discovery stages described below. We used the same bias model in all ATAC bias-factorized ChromBPNet models. We used one of four bias models in most of the DNase models; in some cases, bias factorization with one of the reference models was not successful, which led us to train a custom bias model (i.e., on background regions of the same dataset for which we were training the ChromBPNet model).

##### Prediction

We calculated the predicted counts and profiles for BPNet models and for both the full ChromBPNet models and the bias-factorized ChromBPNet models by running a forward pass of the model on all of the input regions (training, test and validation). To calculate the per-base predicted counts on a per-region basis, we first converted the logit output by the profile head to a probability using a softmax function, and then we multiplied by the per-region counts. For the BPNet models, the total predicted counts are distributed across both strands by converting the logit outputs of both profile heads to a probability using a single softmax function and then multiplying the per-region counts. For BPNet models we also by default perform reverse complement averaging of the predictions. When calculating the model performance as described below, we compare the full ChromBPNet model predictions to the observed data, as the observed data includes enzymatic bias. However, inspecting the bias-factorized ChromBPNet predictions provides a view of the data unaffected by bias, which can be helpful in certain cases, such as when visually inspecting accessibility on a genome browser. Finally, we calculated normalized tracks for each dataset in the following way: (1) we calculated a normalization constant by averaging the counts in the background regions of the observed data; (2) we divided the counts in each of the predicted and observed tracks by this normalization constant.

##### Model interpretation and motif discovery

We interpreted the weights of each trained model by calculating the model’s predicted per-base contribution towards the final output in the peak and ENCODE ccRE regions of the dataset using DeepLIFT/ DeepSHAP.^51^ Specifically, we calculated the relative log fold change in output between the input peak sequence and a GC content-matched, dinucleotide-shuffled reference sequence. This score can be interpreted as where the sum of contribution scores of all bases and positions of a region is equal to the relative log fold change in predicted output for that region, compared to a GC-matched, dinucleotide-shuffled reference region. To assess the contribution of each base to the predicted counts, we calculated the log fold change in predicted counts for the peak region vs. the reference region. To assess the contribution of each base to the predicted profile shape, we calculated log fold change in the sum of predicted logits across the peak region vs. the reference region. We repeated this calculation across all 5 folds of each model and calculated the average scores across all folds over all peak regions.

Next, using the mean contribution scores, we performed motif discovery using TF-MoDISco,^52^ identifying contribution weight matrices (CWMs) of length 50 bp (initial core motif length 20 bp, add initial flank 5 bp, add final flank 10 bp) and 110 bp (initial core motif length 30 bp, add initial flank 20 bp, add final flank 20 bp) within a region of width 400 bp from the peak summit, and a maximum seqlet count of 50,000 for BPNet, and 30 bp (initial core motif length 30 bp, add initial flank 10 bp), within a region of width 500 bp from the peak summit, and a maximum seqlet count of 1,000,000 for chromBPNet, repeated for each output head type.

##### Motif instance calling

Using the motifs discovered, we identified the in-peak genomic occurrences of each motif using FiNeMo ([GitHub Repository](https://github.com/kundajelab/Fi-NeMo)), a GPU-accelerated motif hit caller for identifying occurrences of CWM motifs within deep learning contribution scores. For each set of TF-MoDisCo CWMs and their original contribution scores, we identified motif occurrences within a region of width 1,000 bp from the peak summits and at 3 different sensitivity thresholds (lambda = 0.6, 0.7, 0.8 for BPNet and 0.7, 0.8, 0.9 for chromBPNet).

##### Model performance and quality

We manually curated each BPNet and ChromBPNet model based on the predictive performance and discovered motifs. For BPNet models, we identified 1,617 models that passed the following criteria (“passed”): good model performance, measured by a mean Pearson correlation greater than 0.5 between the predicted and observed log counts in held out test chromosomes; contained one or more known binding motif of the ChIP-ed TF in the discovered motif set, based on a curated set of external TF binding motif databases (JASPAR 2026, HOCOMOCO V14, Codebook, and CIS-BP,^53-55^ using MotifCompendium).^56^ From the remaining models, we flagged 260 models as “archived” based on the following failure criteria: no direct binding motif of the ChIP-ed TF found, despite knowing one or more direct binding motif in the external curated TF binding motif database (excluding ZNFs); very low model performance, measured by a mean Pearson correlation below 0.3 between the predicted and observed log counts in held out test chromosomes; or all discovered motifs identified as low complexity, based on motif complexity metrics from MotifCompendium. The remaining 737 models were categorized as “unvalidated” and included in downstream analyses. While all 1,617 QC-passed models and 737 unvalidated models were used for MotifCompendium clustering, 15 of the QC-passed models were later archived because these experiments were either retracted due to potential antibody cross-contamination or were atypical datasets that used other untreated TF-ChIP experiments as controls, resulting in a total of 1,602 QC-passed models.

We identified 1,512 high-quality ChromBPNet models by running the interpretation and motif discovery steps on a randomly selected 30K subset of all peaks used in training and manually inspected each set of discovered motifs in both the counts and profile heads. If greater than ⅓ of the discovered motifs were identified as low complexity in the profile head, or if we identified a Tn5 or DNase motif in ATAC or DNase datasets, respectively, we retrained the model using a custom bias model and repeated the QC process. If greater than ⅔ of the discovered motifs in the counts head were identified as low complexity, we failed the model without retraining a custom bias model. Models that did not pass QC with a custom bias model were not uploaded. Additionally, models with a median Pearson correlation of less than 0.5 between predicted and observed logcounts across five folds of held-out test chromosomes were not uploaded.

##### Annotating and combining motifs

Using all 2,354 BPNet and 1,512 ChromBPNet models that passed quality checks, we collected all motifs that were discovered by all models and collapsed them into a single, non-redundant set of motifs, while maintaining key motif information, using a hierarchical clustering approach with MotifCompendium.^56^ First, motifs were filtered based on motif complexity scores. Next, motifs were annotated based on external TF binding motif databases.^53-55^ For BPNet-derived motifs, we further identified the binding motifs of the ChIP target TF and categorized these as “direct” binding motifs, while motifs attributed to other TFs not targeted by the ChIP experiment were categorized as “indirect” motifs. Direct motifs were further classified into three categories, based on confidence: (1) Class “A” direct motif: if the direct motif was identified by matching with an orthogonal, in vitro-based experiment’s PFM or a high quality in vivo-based PFM from the external TF binding motif databases; (2) Class “B” direct motif: if the motif was identified by matching with an orthogonal PFM of a paralog of the TF, or in case of a zinc finger protein, successfully reconstructed as a linear combination of each finger’s PFM from bacterial one-hybrid (B1H) experiments^57^; and (3) Class “C” direct motif: if there was no orthogonal evidence to rely on, simply selecting the highest contributing motif, as determined by its mean total contribution score.

Next, motifs were clustered hierarchically, keeping track of the motif’s origin. First: for BPNet, motifs were clustered per ChIP-ed TF, per motif category (direct A, B, C, indirect), per output contribution type (counts, profile); for ChromBPNet, motifs were clustered per tissue annotation, per output contribution type. Second: the centroids of the resultant clusters were further clustered across TF and motif category for BPNet, and across tissue annotation for ChromBPNet. Third: the resultant centroids were again further clustered across contribution types. Fourth: the resultant centroids were clustered one last time, into a unique set of clusters for all BPNet and ChromBPNet models, combined. At each level, clusters were named based on the constituent motifs’ ChIP-ed TF and TF family name if the cluster contained any BPNet-derived direct motifs, and if not, based on the closest annotation TF and TF family name annotation from external TF binding motif databases. Lastly, motif instances previously identified, which each originate from an individual motif, also inherited the hierarchy of clustering membership from its parent motif, creating a full, motif instance-to-4 cluster-level hierarchy.

##### Case study 2 -- MPRA library design

To select sequences containing active TAL and GATA motifs for the MPRA experiment, we intersected the ChIP-seq peaks of TAL1 ([Dataset](https://www.encodeproject.org/experiments/ENCSR000EHB/)), GATA1 ([Dataset](https://www.encodeproject.org/experiments/ENCSR000EFT/)), resulting in 9,989 overlapping loci. We then designed two separate MPRA libraries to assess GATA motif flanks and homotypic motif interactions respectively. For the flank library, we randomly sampled 300 sequences with exactly 1 GATA motif, 65 sequences with 2 motifs and 15 sequences with 3 motifs, where the minimum spacing between any pair of motifs was at least 15bp. For each wildtype sequence of this subset of peaks, we designed variants where either (1) the flanking positions surrounding each GATA motif were replaced with random nucleotides, (2) the GATA motif was scrambled by shuffling its nucleotides, or (3) both the GATA motif and its flanks were simultaneously perturbed with identical mutations as in (1)-(2). These perturbations were repeated 3 times per wildtype sequence, resulting in a total of 4,655 oligos (including wildtypes).

The homotypic GATA pair library was constructed similarly to the flank library but included 600 wildtype sequences with 2 GATA motifs and 100 sequences with 3 motifs (and 0 sequences with one motif, as we wanted to measure pairwise interactions), without any spacing threshold between the motif pairs. For each wildtype sequence, we designed variants where the nucleotides of each GATA motif were shuffled in isolation, as well as variants where all GATA motifs were scrambled simultaneously. This resulted in 8,200 oligos with 3 repeats per variant. Finally, for a small set of sequences sampled from both libraries, we designed a total of 1,305 control oligos where arbitrarily chosen regions were scrambled (as opposed to GATA flanks or neighboring motifs), allowing us to verify the absence of epistasis for neutral sequence. The final library oligos were 200 bp long windows, chosen by centering on the TAL1 ChIP-seq peak summits.

##### Case study 2 -- MPRA vector assembly

The MPRA libraries were constructed as previously described.^9^ 15,000 oligonucleotides were synthesized (Agilent Technologies) as 230 bp sequences consisting of 200 bp of genomic sequence and 15 bp adaptors on either end. Unique 20 bp barcodes were added via PCR using primers 2E-A082 and 2E-A202 (see below) and cycle conditions: 98 °C for 30 s, 6 cycles (98 °C for 10 s, 60 °C for 15 s, 65 °C for 45s), 72 °C for 5 min. The resulting products were subsequently incorporated into a SfiI-digested backbone pGL4.23∆luc:∆xbaI vector (Addgene #109035) via Gibson assembly. The oligo library was expanded by electroporation into 10-beta electrocompetent E.coli (New England Biolabs, catalog no. C3020K), transformed cells were immediately split across 10 independent cultures, and the desired number of expanded cultures were selected after outgrowth to reach an average of 300 colony-forming units (CFUs) per oligonucleotide sequence tested (4.5 million total). The resulting pGL4.23:MPRA∆ORF plasmid library was sequenced using Illumina 2x150bp chemistry to acquire oligo-barcode pairings. The library was linearized via AsiSI restriction digestion (New England Biolabs, catalog no. R0630S), and GFP with a minimal TATA promoter was inserted between the 200bp oligo sequence and 20bp barcode using Gibson assembly. Finally, the library was transformed in E.coli achieving 90 million CFUs and expanded for 16 hours at 30C to generate a final pGL4.23:MPRA library.

##### Case study 2 -- Cell culture and MPRA library transfection

K562 cells (ATCC, catalog no. CCL-243) were cultured in RPMI 1640 Medium, GlutaMAX™ Supplement (Life Technologies, catalog no. 61870127) + 10% fetal bovine serum (Fisher Scientific, catalog no. A3160402) to a density of 1 million cells per mL prior to transfection. One hundred million K562 cells were transfected per replicate using the Neon Transfection System 100 µl Kit (Thermo Fisher Scientific, catalog no. MPK10096) with three pulses at 1450 V for 10 ms and 5 µg of the MPRA library per ten million cells. Twenty-four hours after transfection, cells were harvested, rinsed twice with PBS and collected by centrifugation. Cells were mechanically homogenized in RLT buffer (Qiagen, catalog no. 75162) and dithiothreitol (Fisher Scientific, catalog no. 50-103-6063), and cell homogenates were subsequently frozen at -80 °C. There were 5 biological replicates with no more than two replicates performed on the same day with same day replicates using independently expanded batches of cells.

##### RNA isolation and MPRA RNA library generation

RNA was extracted from frozen cell homogenates using the Qiagen RNeasy Maxi kit and treated with SUPERase•In™ RNase Inhibitor (Invitrogen, catalog no. AM2696). Following treatment with Turbo™ DNase (Life Technologies, catalog no. AM2239), GFP transcripts were captured using Cytiva Sera-Mag SpeedBeads™ Carboxyl Magnetic Beads (Fisher Scientific, catalog no. 09-981-123) and a 100 µM mixture of three GFP-specific biotinylated primers 2E-A120, 2E-A123 and 2E-A126. After another round of SUPERase•In™ and DNase treatment, complementary DNA was synthesized using primer 2E-A019 and SuperScript™ III First-Strand Synthesis SuperMix (Life Technologies, catalog no. 18080400). GFP mRNA abundance was then quantified by qPCR using NEBNext® Ultra™ II Q5® Master Mix (New England Biolabs, catalog no.M0544S) to determine the cycle at which linear amplification begins for each replicate. Replicates and the plasmid library were diluted to approximately the same concentration based on the qPCR results, and samples were amplified with primers 2E-A801 and 2E-A802 with the following qPCR conditions: 98 °C for 20 s, 8 cycles (98 °C for 10 s, 62 °C for 15 s, 72 °C for 30 s), 72 °C for 2 min to amplify barcodes associated with GFP mRNA sequences for each replicate. Subsequent PCR (same conditions, 6 cycles) was used to add Illumina sequencing adaptors to the replicates. The resulting indexed MPRA sequencing libraries were spiked with 1% PhiX and sequenced on an Illumina NovaSeq S2 for 20 bp and dual 8 bp i5 and i7 index reads.

| **Primer number** | **Primer sequence** | **Vendor** | **Purpose** |
| --- | --- | --- | --- |
| 2E-A082 | GCCAGAACATTTCTCTGGCCTAACTGGCCGCTTGACG | IDT | 20bp barcode addition |
| 2E-A202 | CCGACTAGCTTGGCCGCCGACGCTCTTCCGATCT(N1:25252525)(N1)(N1)(N1)(N1)(N1)(N1)(N1)(N1)(N1)(N1)(N1)(N1)(N1)(N1)(N1)(N1)(N1)(N1)(N1)TCTAGAGGTTCGTCGACGCGATCGCAGGAGCCGCAGTG | IDT | 20bp barcode addition |
| 2E-A120 | CCTCGATGTTGTGGCGGGTCTTGAAGTTCACCTTG/3BioTEG/ | IDT | GFP mRNA pulldown |
| 2E-A123 | CCAGGATGTTGCCGTCCTCCTTGAAGTCGATGCCC/3BioTEG/ | IDT | GFP mRNA pulldown |
| 2E-A126 | CGCCGTAGGTGAAGGTGGTCACGAGGGTGGGCCAG/3BioTEG/ | IDT | GFP mRNA pulldown |
| 2E-A19 | CCGACTAGCTTGGCCGC | IDT | cDNA preparation from GFP mRNA |
| 2E-A801 | ACTGGAGTTCAGACGTGTGCTCTTCCGATCTCGCCCTGAGCAAAGA*C*C | IDT | qPCR amplification of GFP mRNA |
| 2E-A802 | ACTCTTTCCCTACACGACGCTCTTCCGAT*C*T | IDT | qPCR amplification of GFP mRNA |

##### Case study 2 -- MPRA data processing and analysis

Illumina reads from Oligo-barcodes sequencing of pGL4.23:MPRA∆ORF were processed using MPRAmatch. Briefly, MPRAmatch uses FLASH2^58^ to merge overlapping Illumina reads, MPRAmatch retrieves the barcodes and oligonucleotides from each complete sequence, and maps the oligonucleotides to the original oligonucleotide design file using Minimap2.^59^ A total of 14,994 out of the original 15,000 oligonucleotides were recovered (99.96% capture rate) from the pGL4.23:MPRA∆ORF library. Of all the barcodes recovered, ~92% were mapped to a single oligonucleotide with the remaining ~8% either mapping to multiple oligonucleotides or linked to a sequence that failed mapping QC.

Tag reads from the RNA and plasmid replicates were processed using MPRAcount. Barcodes were assigned a unique oligonucleotide for 93.6% of barcodes from the plasmid replicates and 95.5% of barcodes from K562 replicates. In total across all the replicates, 14,985 oligonucleotides were recovered with 14,908 oligonucleotides having an aggregated barcode count of greater than 20 (99.4%) in the plasmid and 14,769 oligonucleotides (98.5%) in K562.

A barcode level count table was processed in MPRAmodel, which uses DESeq2,^60^ to calculate the activity of all the oligonucleotides and produce p-values using a negative binomial model with pooled dispersion estimates. Library normalization was performed using the summit shift normalization method in MPRAmodel. This approach uses a predefined negative control group (ORF elements) to scale the log_2_(FoldChange) activity estimates for the entire library so the peak of the negative control distribution centers over zero.

##### Case study 2 -- Deep learning models

We utilized the single-task BPNet models on the TAL1 ([ENCSR000EHB](https://www.encodeproject.org/experiments/ENCSR000EHB/)), GATA1 ([ENCSR000EFT](https://www.encodeproject.org/experiments/ENCSR000EFT/)) and GATA2 ([ENCSR000BKM](https://www.encodeproject.org/experiments/ENCSR000BKM/)) and ChromBPNet models on ATAC-seq ([ENCSR868FGK](https://www.encodeproject.org/experiments/ENCSR868FGK/)) and DNase-seq ([ENCSR000EOT](https://www.encodeproject.org/experiments/ENCSR000EOT/)) in K562. In addition to the models, we also trained a ReporterNet model ([ENCSR701UIV](https://www.encodeproject.org/experiments/ENCSR701UIV/)) on a large-scale MPRA dataset measured in K562 from prior work. These MPRA data were mostly designed to assess the effects of variants associated with complex human traits sourced from UKBB and GTEX.^9,61^ The MPRA model shares the same architecture as the standard BPNet, with the exception that it lacks a profile head. Instead, it features a single prediction head that outputs the activity of each sequence, which is measured by the Log2 fold change between RNA and DNA. We used 200 bp flanking sequences from the MPRA construct, both upstream and downstream of the central 200 bp variant sequences for training the MPRA model. All models were trained with five-fold cross-validation (CV) and random reverse complement augmentation.

For the ChIP, ATAC, and DNase-seq models, we computed the marginalized effects of the MPRA oligos within the central 600 bp region, which encompassed the 200-bp variant sequences along with 200 bp flanking sequences upstream and downstream from the MPRA construct. We computed the marginalized effects by sampling 20 dinucleotide-shuffled sequences as additional flanking sequences beyond the central 600 bp region. The predictions were averaged across the 20 dinucleotide-shuffled sequences for each model fold and finally averaged across the 5 cross-validation folds. We used the log-count output heads for predictions and attributions.

### References for Supplementary Materials

1	The GTEx Consortium atlas of genetic regulatory effects across human tissues. *Science* **369**, 1318–1330 (2020). <https://doi.org/10.1126/science.aaz1776>

2	Rozowsky, J. *et al.* The EN-TEx resource of multi-tissue personal epigenomes & variant-impact models. *Cell* **186**, 1493–1511.e1440 (2023). <https://doi.org/10.1016/j.cell.2023.02.018>

3	Zhang, K. *et al.* A single-cell atlas of chromatin accessibility in the human genome. *Cell* **184**, 5985–6001.e5919 (2021). <https://doi.org/10.1016/j.cell.2021.10.024>

4	McShane, A. *et al.* Characterizing nascent transcription patterns of PROMPTs, eRNAs, and readthrough transcripts in the ENCODE4 deeply profiled cell lines. *bioRxiv* (2024). <https://doi.org/10.1101/2024.04.09.588612>

5	Bedi, K. *et al.* Isoform and pathway-specific regulation of post-transcriptional RNA processing in human cells. *bioRxiv* (2024). <https://doi.org/10.1101/2024.06.12.598705>

6	Rao, S. S. *et al.* A 3D map of the human genome at kilobase resolution reveals principles of chromatin looping. *Cell* **159**, 1665–1680 (2014). <https://doi.org/10.1016/j.cell.2014.11.021>

7	Guckelberger, P. *et al.* Cohesin-mediated 3D contacts tune enhancer-promoter regulation. *bioRxiv* (2024). <https://doi.org/10.1101/2024.07.12.603288>

8	Rebboah, E. *et al.* The ENCODE mouse postnatal developmental time course identifies regulatory programs of cell types and cell states. *bioRxiv* (2024). <https://doi.org/10.1101/2024.06.12.598567>

9	Siraj, L. *et al.* Functional dissection of complex and molecular trait variants at single nucleotide resolution. *bioRxiv* (2024). <https://doi.org/10.1101/2024.05.05.592437>

10	Siraj, L. *et al.* Functional dissection of complex trait variants at single-nucleotide resolution. *Nature* (2026). <https://doi.org/10.1038/s41586-026-10121-6>

11	Abramov, S. *et al.* Landscape of allele-specific transcription factor binding in the human genome. *Nat Commun* **12**, 2751 (2021). <https://doi.org/10.1038/s41467-021-23007-0>

12	Vierstra, J. *et al.* Global reference mapping of human transcription factor footprints. *Nature* **583**, 729–736 (2020). <https://doi.org/10.1038/s41586-020-2528-x>

13	Anttila, V., Wessman, M., Kallela, M. & Palotie, A. Towards an understanding of genetic predisposition to migraine. *Genome Med* **3**, 17 (2011). <https://doi.org/10.1186/gm231>

14	Majoros, W. H. *et al.* Bayesian estimation of genetic regulatory effects in high-throughput reporter assays. *Bioinformatics* **36**, 331–338 (2020). <https://doi.org/10.1093/bioinformatics/btz545>

15	DeGorter, M. K. *et al.* Transcriptomics and chromatin accessibility in multiple African population samples. *bioRxiv*, 2023.2011.2004.564839 (2023). <https://doi.org/10.1101/2023.11.04.564839>

16	Moore, J. E. *et al.* An expanded registry of candidate cis-regulatory elements. *Nature* (2026). <https://doi.org/10.1038/s41586-025-09909-9>

17	Cochran, K. *et al.* Dissecting the cis-regulatory syntax of transcription initiation with deep learning. *bioRxiv* (2024). <https://doi.org/10.1101/2024.05.28.596138>

18	Avsec, Ž. *et al.* Base-resolution models of transcription-factor binding reveal soft motif syntax. *Nat Genet* **53**, 354–366 (2021). <https://doi.org/10.1038/s41588-021-00782-6>

19	Pampari, A. *et al.* ChromBPNet: bias factorized, base-resolution deep learning models of chromatin accessibility reveal cis-regulatory sequence syntax, transcription factor footprints and regulatory variants. *bioRxiv* (2025). <https://doi.org/10.1101/2024.12.25.630221>

20	Kim, D. S. *et al.* The dynamic, combinatorial cis-regulatory lexicon of epidermal differentiation. *Nat Genet* **53**, 1564–1576 (2021). <https://doi.org/10.1038/s41588-021-00947-3>

21	Liu, B. B. *et al.* Dissecting regulatory syntax in human development with scalable multiomics and deep learning. *bioRxiv*, 2025.2004.2030.651381 (2025). <https://doi.org/10.1101/2025.04.30.651381>

22	Franke, A. *et al.* Replication of signals from recent studies of Crohn's disease identifies previously unknown disease loci for ulcerative colitis. *Nat Genet* **40**, 713–715 (2008). <https://doi.org/10.1038/ng.148>

23	Parkes, M. *et al.* Sequence variants in the autophagy gene IRGM and multiple other replicating loci contribute to Crohn's disease susceptibility. *Nat Genet* **39**, 830–832 (2007). <https://doi.org/10.1038/ng2061>

24	Grasberger, H. *et al.* STR mutations on chromosome 15q cause thyrotropin resistance by activating a primate-specific enhancer of MIR7-2/MIR1179. *Nat Genet* **56**, 877–888 (2024). <https://doi.org/10.1038/s41588-024-01717-7>

25	Blache, P. *et al.* SOX9 is an intestine crypt transcription factor, is regulated by the Wnt pathway, and represses the CDX2 and MUC2 genes. *J Cell Biol* **166**, 37–47 (2004). <https://doi.org/10.1083/jcb.200311021>

26	van der Flier, L. G. *et al.* Transcription factor achaete scute-like 2 controls intestinal stem cell fate. *Cell* **136**, 903–912 (2009). <https://doi.org/10.1016/j.cell.2009.01.031>

27	Granja, J. M. *et al.* ArchR is a scalable software package for integrative single-cell chromatin accessibility analysis. *Nat Genet* **53**, 403–411 (2021). <https://doi.org/10.1038/s41588-021-00790-6>

28	Korsunsky, I. *et al.* Fast, sensitive and accurate integration of single-cell data with Harmony. *Nat Methods* **16**, 1289–1296 (2019). <https://doi.org/10.1038/s41592-019-0619-0>

29	Stik, G. *et al.* CTCF is dispensable for immune cell transdifferentiation but facilitates an acute inflammatory response. *Nature Genetics* **52**, 655–661 (2020). <https://doi.org/10.1038/s41588-020-0643-0>

30	Visser, I. & Speekenbrink, M. depmixS4: An R Package for Hidden Markov Models. *Journal of Statistical Software* **36**, 1 – 21 (2010). <https://doi.org/10.18637/jss.v036.i07>

31	Morizane, R. *et al.* Nephron organoids derived from human pluripotent stem cells model kidney development and injury. *Nat Biotechnol* **33**, 1193–1200 (2015). <https://doi.org/10.1038/nbt.3392>

32	Gupta, N. *et al.* Modeling injury and repair in kidney organoids reveals that homologous recombination governs tubular intrinsic repair. *Sci Transl Med* **14**, eabj4772 (2022). <https://doi.org/10.1126/scitranslmed.abj4772>

33	Reese, F. *et al.* The ENCODE4 long-read RNA-seq collection reveals distinct classes of transcript structure diversity. *bioRxiv* (2023). <https://doi.org/10.1101/2023.05.15.540865>

34	Reese, F. & Mortazavi, A. Swan: a library for the analysis and visualization of long-read transcriptomes. *Bioinformatics* **37**, 1322–1323 (2021). <https://doi.org/10.1093/bioinformatics/btaa836>

35	Joglekar, A. *et al.* A spatially resolved brain region- and cell type-specific isoform atlas of the postnatal mouse brain. *Nat Commun* **12**, 463 (2021). <https://doi.org/10.1038/s41467-020-20343-5>

36	Fabiha, T. *et al.* A consensus variant-to-function score to functionally prioritize variants for disease. *bioRxiv*, 2024.2011.2007.622307 (2024). <https://doi.org/10.1101/2024.11.07.622307>

37	Gschwind, A. R. *et al.* An encyclopedia of enhancer-gene regulatory interactions in the human genome. *bioRxiv* (2023). <https://doi.org/10.1101/2023.11.09.563812>

38	Berisa, T. & Pickrell, J. K. Approximately independent linkage disequilibrium blocks in human populations. *Bioinformatics* **32**, 283–285 (2016). <https://doi.org/10.1093/bioinformatics/btv546>

39	Jerome, H. F. Greedy function approximation: A gradient boosting machine. *The Annals of Statistics* **29**, 1189–1232 (2001). <https://doi.org/10.1214/aos/1013203451>

40	Chen, T. & Guestrin, C. in *Proceedings of the 22nd acm sigkdd international conference on knowledge discovery and data mining.* 785–794.

41	Kim, S. S. *et al.* Improving the informativeness of Mendelian disease-derived pathogenicity scores for common disease. *Nat Commun* **11**, 6258 (2020). <https://doi.org/10.1038/s41467-020-20087-2>

42	Dey, K. K. *et al.* Integrative approaches to improve the informativeness of deep learning models for human complex diseases. *bioRxiv*, 2020.2009.2008.288563 (2021). <https://doi.org/10.1101/2020.09.08.288563>

43	Lundberg, S. M. & Lee, S.-I. A unified approach to interpreting model predictions. *Advances in neural information processing systems* **30** (2017).

44	Wang, Q. S. *et al.* Leveraging supervised learning for functionally informed fine-mapping of cis-eQTLs identifies an additional 20,913 putative causal eQTLs. *Nat Commun* **12**, 3394 (2021). <https://doi.org/10.1038/s41467-021-23134-8>

45	Wang, G., Sarkar, A., Carbonetto, P. & Stephens, M. A simple new approach to variable selection in regression, with application to genetic fine mapping. *J R Stat Soc Series B Stat Methodol* **82**, 1273–1300 (2020). <https://doi.org/10.1111/rssb.12388>

46	Tanigawa, Y. & Kellis, M. Power of inclusion: Enhancing polygenic prediction with admixed individuals. *Am J Hum Genet* **110**, 1888–1902 (2023). <https://doi.org/10.1016/j.ajhg.2023.09.013>

47	Tian, X. *et al.* PRISM: ancestry-aware integration of tissue-specific genomic annotations enhances the transferability of polygenic scores. *bioRxiv* (2025). <https://doi.org/10.1101/2025.11.13.688144>

48	Bycroft, C. *et al.* The UK Biobank resource with deep phenotyping and genomic data. *Nature* **562**, 203–209 (2018). <https://doi.org/10.1038/s41586-018-0579-z>

49	Gaziano, J. M. *et al.* Million Veteran Program: A mega-biobank to study genetic influences on health and disease. *J Clin Epidemiol* **70**, 214–223 (2016). <https://doi.org/10.1016/j.jclinepi.2015.09.016>

50	Verma, A. *et al.* Diversity and scale: Genetic architecture of 2068 traits in the VA Million Veteran Program. *Science* **385**, eadj1182 (2024). <https://doi.org/10.1126/science.adj1182>

51	Shrikumar, A., Greenside, P. & Kundaje, A. in *International conference on machine learning.* 3145–3153 (PMlR).

52	Shrikumar, A. *et al.* Technical note on transcription factor motif discovery from importance scores (TF-MoDISco) version 0.5. 6.5. *arXiv preprint arXiv:1811.00416* (2018).

53	Ovek Baydar, D. *et al.* JASPAR 2026: expansion of transcription factor binding profiles and integration of deep learning models. *Nucleic Acids Res* **54**, D184–d193 (2026). <https://doi.org/10.1093/nar/gkaf1209>

54	Vorontsov, I. E. *et al.* HOCOMOCO in 2024: a rebuild of the curated collection of binding models for human and mouse transcription factors. *Nucleic Acids Res* **52**, D154–d163 (2024). <https://doi.org/10.1093/nar/gkad1077>

55	Weirauch, M. T. *et al.* Determination and inference of eukaryotic transcription factor sequence specificity. *Cell* **158**, 1431–1443 (2014). <https://doi.org/10.1016/j.cell.2014.08.009>

56	Deshpande, S. *et al.* *A unified lexicon of predictive DNA sequence motifs from ENCODE transcription factor binding and chromatin accessibility assays*. (2025).

57	Najafabadi, H. S., Albu, M. & Hughes, T. R. Identification of C2H2-ZF binding preferences from ChIP-seq data using RCADE. *Bioinformatics* **31**, 2879–2881 (2015). <https://doi.org/10.1093/bioinformatics/btv284>

58	Magoc, T. & Salzberg, S. L. FLASH: fast length adjustment of short reads to improve genome assemblies. *Bioinformatics* **27**, 2957–2963 (2011). <https://doi.org/10.1093/bioinformatics/btr507>

59	Li, H. Minimap2: pairwise alignment for nucleotide sequences. *Bioinformatics* **34**, 3094–3100 (2018). <https://doi.org/10.1093/bioinformatics/bty191>

60	Love, M. I., Huber, W. & Anders, S. Moderated estimation of fold change and dispersion for RNA-seq data with DESeq2. *Genome Biol* **15**, 550 (2014). <https://doi.org/10.1186/s13059-014-0550-8>

61	Gosai, S. J. *et al.* Machine-guided design of cell-type-targeting cis-regulatory elements. *Nature* **634**, 1211–1220 (2024). <https://doi.org/10.1038/s41586-024-08070-z>
